# A Dimeric SINE Discovered in Shrew Mole is Structurally Similar to Primate Alu

**DOI:** 10.1101/2024.08.25.609555

**Authors:** Sergey A. Kosushkin, Nikita S. Vassetzky, Olga R. Borodulina, Dmitri A. Kramerov

## Abstract

Families of Short Interspersed Elements (SINEs) originate from tRNA, 5S rRNA, and 7SL RNA molecules in the genomes of multicellular organisms. Families of 7SL RNA-derived SINEs are very rare; however, they have been best studied in primates and rodents. The rodent B1 SINE is a monomeric element, whereas the primate Alu is composed of two 7SL RNA-derived monomers.We found that in contrast to other members of the family Talpidae (moles), which possess a tRNA-derived Tal SINE, the shrew mole *Uropsilus gracilis* contains 280,000 genomic copies of the previously unknown 7SL RNA-derived Urop SINE. Like Alu, Urop consists of two monomers connected by an A-rich linker. The origin of the Urop monomers, like that of the Alu and B1 monomers, was mediated by several essentially identical events - a long central deletion in the 7SL RNA, dimerization, and/or internal duplication. Urop copies can be divided into three subfamilies (a, b, and c), the latter being the most numerous and recent. Urop_c has more copies with poly(A) tails longer than 50 compared to other L1-mobilized SINEs. Urop and Alu illustrate an independent emergence in the evolutionary history of highly similar SINEs.

## Introduction

Short Interspersed Elements (SINEs) are non-autonomous mobile genetic retroelements less than 600 bp in length that are transcribed by RNA polymerase III (pol III) from an internal promoter.SINEs are characteristic of the vast majority of multicellular organisms, and the number of their copies in the genome can reach one million (Kramerov and Vassetzky 2011b). Their new genomic copies are generated by reverse transcription of SINE RNA by an enzyme of a Long Interspersed Element (LINE) present in the same genome. In placental mammals, most SINE families use L1 reverse transcriptase for their retrotransposition, which requires a poly(A) tail at the end of the SINE RNA. The integration of SINEs into new genomic loci often has no effect on host function, but sometimes it can either disrupt gene function, causing inherited diseases (Chen, et al. 2006), or introduce new regulatory mechanisms of gene transcription as well as pre-mRNA splicing and polyadenylation (Ferrigno, et al. 2001; Krull, et al. 2005; Chen, et al. 2009). A new SINE family that emerges in evolution becomes characteristic of all species that diverge from that lineage of host organisms. Thus, SINE families can be characteristic of an order or limited to a few mammalian families (Kramerov and Vassetzky 2011a).

Most SINE families originate from one of the tRNA species (Vassetzky and Kramerov 2013). 5S rRNA-derived SINE families are rare, while the formation of SINEs from 7SL RNA, a component of signal recognition particles, seems to be an exceptional event. Surprisingly, 7SL RNA-derived SINEs were the first to be discovered: Alu and B1 in the human and mouse genomes, respectively (Krayev, et al. 1980; Deininger, et al. 1981). Further research has shown that Alu is characteristic of the primate group (Deininger 2011), while B1 is present in the genomes of all rodents studied (Veniaminova, et al. 2007). In addition, not numerous 7SL RNA-derived SINEs (Tu types I, II, and m) have been found in the genomes of tree shrews (Nishihara, et al. 2002; Vassetzky, et al. 2003; Kriegs, et al. 2007). Tree shrews, primates and rodents belong to the Supraprimates clade, which makes the common origin of Alu, B1, and Tu SINEs highly probable.

A precursor SINE probably arose in the supraprimate ancestor after a 183-bp deletion in the central region of the 7SL RNA sequence (299 nt). This low-copy SINE is called free left Alu monomer (FLAM) in primates and proto-B1 (pB1) in rodents. In primates, another precursor, the free right Alu monomer (FRAM), arose after two shorter deletions, 155 and 11 nt, in the central part of 7SL RNA. A combination of FLAM and FRAM gave rise to the highly successful dimeric SINE Alu with over one million copies in the human genome. In rodents, specific 7─10-nt deletions within the same region generated pB1d7, pB1d9, and pB1d10 SINEs; and the subsequent 29-bp tandem duplication in pB1d9 (sometimes called quasi-dimerization in parallel with dimeric Alu) resulted in a highly successful B1 SINE in murids (half a million copies in the mouse genome). In tree shrews, the only detectable fossil SINE corresponds to the rodent pB1d SINEs (the deletion size here is 13 nt).

While the ancestral 7SL RNA-derived SINEs are monomeric, their more successful descendants are dimeric with two 7SL RNA units (Alu in primates), 7SL RNA-tRNA or tRNA-7SL RNA units (B1-dID and SP-D-Geo or B4, Men, and SINE type II in rodents and lower primates; for references see (Vassetzky and Kramerov 2013)). Tree shrews have both dimeric and trimeric SINEs (Tu types II and III), and their leftmost unit is derived from a tRNA (Nishihara, et al. 2002) or has a dual 7SL RNA and tRNA origin (Kriegs, et al. 2007). While murid B1 is a remarkable example of a successful monomeric 7SL RNA-derived SINE, this quasi-dimer has an internal duplication.

SINEs derived from 7SL RNA have not been found outside of supraprimates with the sole exception of SINE1-1_EBu (hereafter, Ebu1-1) described in the hagfish *Eptatretus burgeri*, whose genome contains 2363 copies of this element (Kojima 2020). Similar to other 7SL RNA-derived SINEs, Ebu1-1 has a deletion in approximately the same region of the 7SL RNA as other SINEs (see below). 7SL RNA-derived SINEs in supraprimates are mobilized by L1, which does not require any sequences other than the poly(A) tail. Ebu1-1 is mobilized by a hagfish LINE from the RTE clade and accordingly contains a bipartite recognition region from the RNA of this LINE and an (AAC)_n_ tail.

Here, our search for SINEs in sequenced mole genomes (Talpidae) unexpectedly revealed neither Tal, thought to be typical of the family, nor other specific tRNA-derived SINEs in the genome of the gracile shrew mole (*Uropsilus gracilis*). Instead, we found more than 280,000 copies of a 7SL RNA-derived SINE called Urop. This SINE is absent from the genomes of other moles, consistent with an early split of Uropsilinae from the main lineage of moles.

## Results

Previously, we described four families of tRNA-derived SINEs from the genomes of the order Eulipotyphla: Eri-1 and Eri-2 (Erinaceidae, hedgehogs and moonrats), Sor (Soricidae, shrews), and Tal (Talpidae, moles) based on a limited number of sequenced copies (Borodulina and Kramerov 2001). Since then, the sequenced genomes of many mammals, including rare and exotic species, have become available, allowing us to search for SINE sequences in eulipotyphlan genomes. First, we found an intrinsic tRNA-derived SINE (Sol) in the genome of *Solenodon paradoxus* (deposited in SINEBase (Vassetzky and Kramerov 2013)), which belongs to Solenodontidae, another Eulipotyphla family. Second, the unambiguous assignment of Eri1 and Eri2 SINEs to the family Erinaceidae was confirmed as well as Sor SINE to Soricidae, and Tal SINE to Talpidae (Fig. 1A).Third, we unexpectedly found no Tal or any other specific tRNA-derived (or 5S RNA-derived) SINE in the genome of the gracile shrew mole *Uropsilus gracilis* (Talpidae). However, more than 280,000 copies of a dimeric 7SL RNA-derived SINE named Urop were found in this genome (Fig. 1A). All copies of Urop were grouped into three subfamilies: a, b, and c (Fig. 1B). Urop_c was the most abundant subfamily (200,883 copies), Urop_a had 56,323 copies, and Urop_b had 24,864 copies. The Urop_a consensus looks most basal of three; unlike Urop_b and Urop_c, its sequences are more similar to the ancestral Urop-m (see below) (Fig. 1B). The left and right monomers are connected by A-rich linkers.

**Fig. 1.**
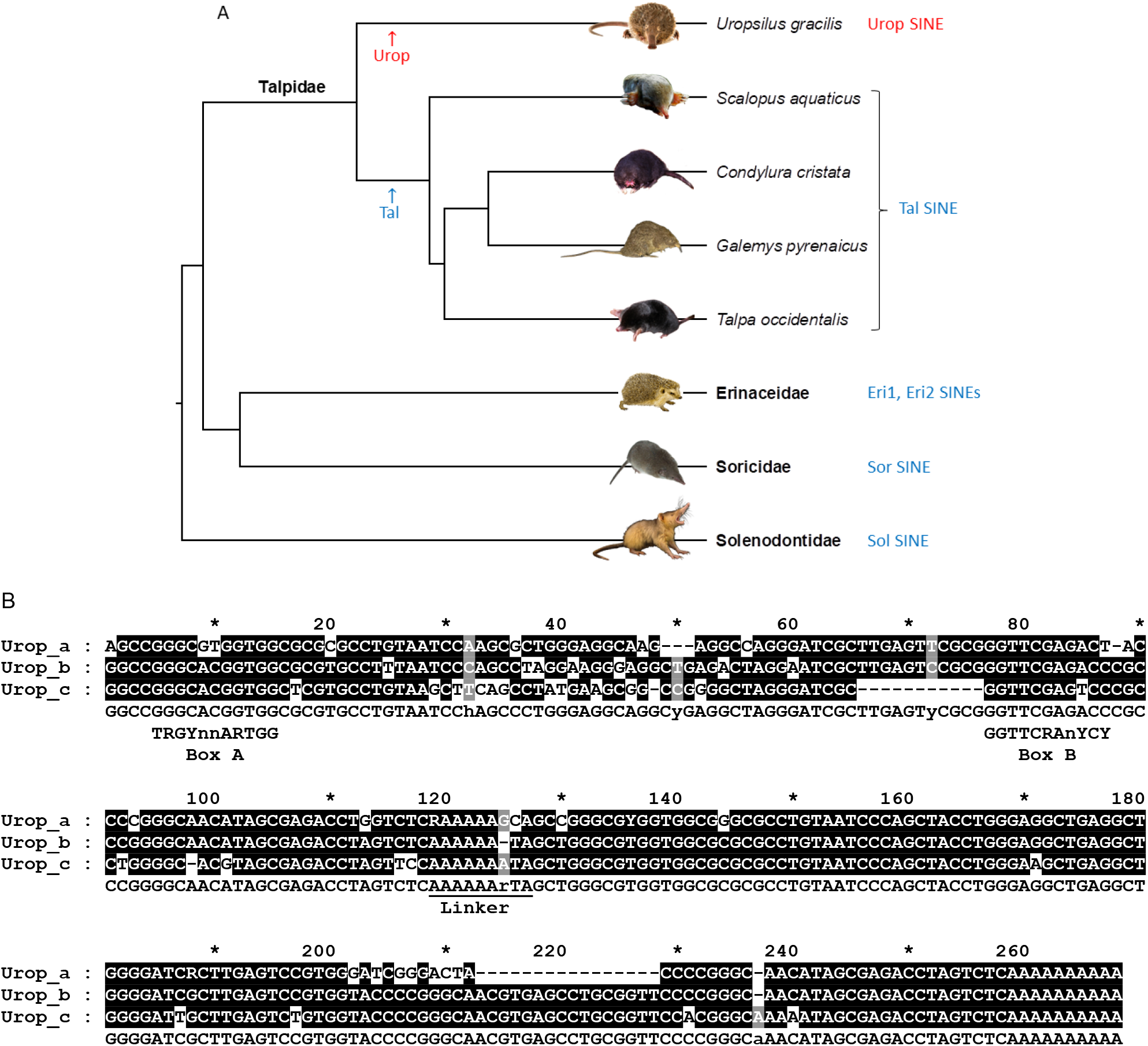
Urop, a SINE family from the genome of the shrew mole *Uropsilus gracilis*. **A**. SINE families identified in the genomes of four Eulipotyphla families. The names of tRNA-related SINEs are shown in blue, and the 7SL RNA-related Urop is highlighted in red. The Eulipotyphla tree shows five Talpidae species whose genomes have been sequenced. The putative time of origin of Tal and Urop SINEs are indicated by arrows. **B**. Alignment of the consensus sequences of the three Urop subfamilies. 7SL RNA-related monomers are connected by a linker (underlined). Consensus boxes A and B of the pol III promoters are shown below the similar SINE regions of Urop.

Fig. 2A shows an alignment of 7SL RNAs with related regions of the left (L) and right (R) monomers of each of the three Urop subfamilies, the L and R monomers of human Alu and their fossil precursors (FLAM and FRAM), rodent B1 precursors (pB1 and pB1D10), right and central Tu type I and Tu type II monomers of tree shrew, and Ebu1-1 of the hagfish. The consensus of each of the Urop monomers shows similarity to approximately the same 5’ and 3’ end regions of the 7SL RNA as other 7SL RNA-derived SINEs; in other words, they all originated from the 7SL RNA sequence after a long deletion in its central portion. The size (∼183 nt) and position of this deletion in the left Urop monomers and Urop_a_R is consistent with that in Alu_L, FLAM, pB1, and pB1d10/TuII-M.

**Fig. 2.**
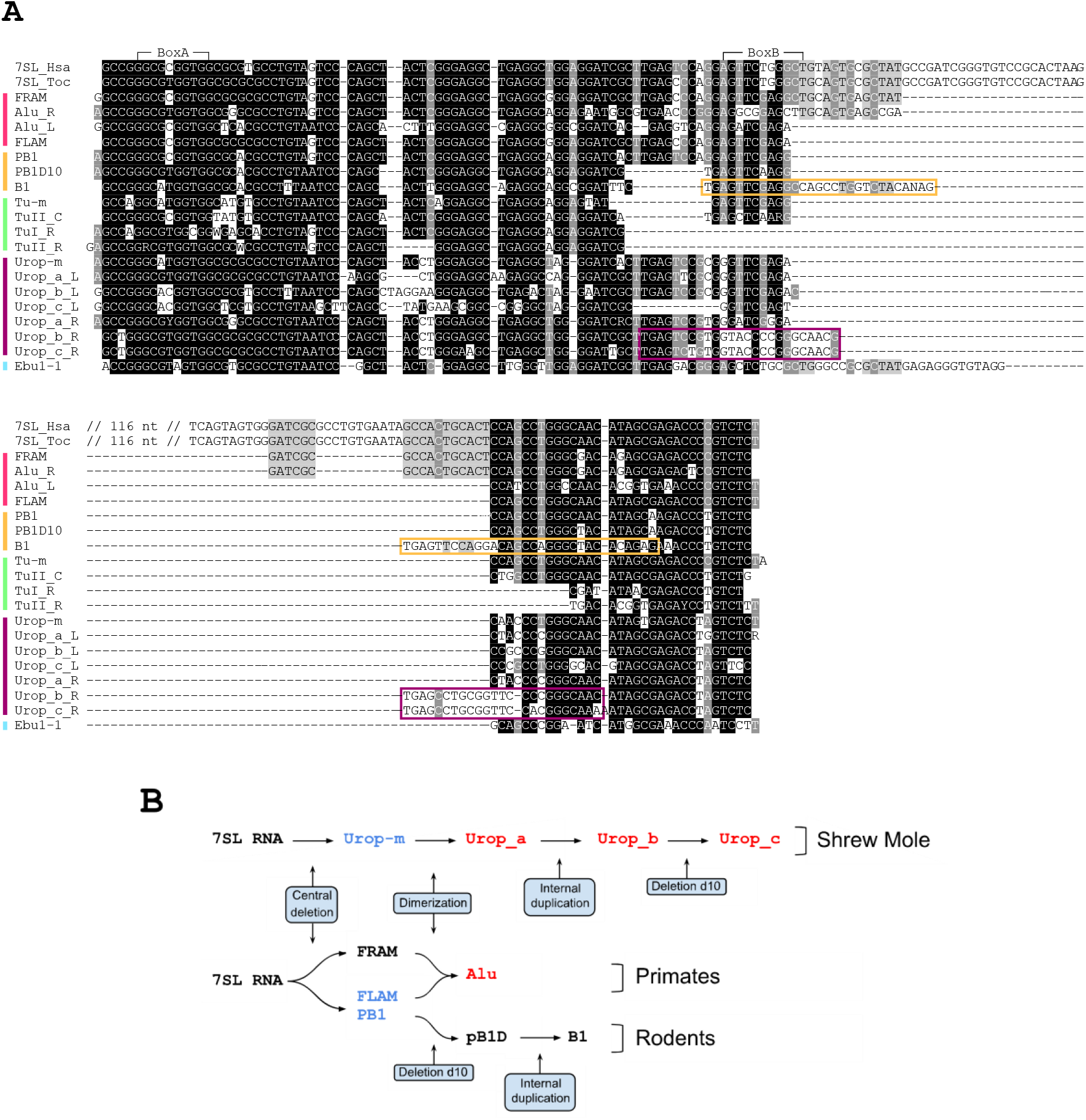
Urop, Alu and other 7SL-derived SINEs: structure, origin and evolution. **A**. Alignment of the human 7SL RNA sequences with the consensus sequences of the monomers of the three Urop subfamilies, human Alu, tree shrew Tu types I and II, mouse B1, and hagfish Ebu1-1. The names of the left, central, and right monomers are indicated by ‘_L,’ ‘_C,’ and ‘_R’ suffixes, respectively. The consensus sequences of the fossil human Alu (FLAM and FRAM), mouse B1 (pB1 and pB1d10), tree shrew Tu (Tu-m), and shrew mole Urop (Urop-m) are also included. Apart from the human 7SL RNA gene, the one from the mole *Talpa occidentalis* is given for reference (7SL_Hsa and 7SL_Toc, respectively). The central section of 7SL RNA (116 nt) missing in the considered SINEs was removed from the alignment. The position of boxes A and B are marked above. Direct repeats (duplications) in B1, Urop_b_R and Urop_c_R are boxed in orange and purple, respectively. **B**. Similar events underlying the evolution of SINEs from 7SL RNA in shrew mole, primates, and rodents.

The right monomers of Urop_b and Urop_c acquired a duplication in the region spanning 24 nt, which largely overlaps the 29-nt quasi-dimer in rodent B1 albeit shorter (boxed in purple and orange, respectively, Fig. 2A). Note also an additional 11-nt deletion in the left Urop_c monomer at a position similar to pB1D10, B1, Tu-m, and TuII_C. It is also worth to mention the absence of box B in certain right monomers of 7SL RNA-derived SINEs (Urop_b/c and Tu types II/III), which is not unexpected since a complete promoter (boxes A and B) is required in the 5’ part of a gene for pol III transcription initiation.

Apart from the dimeric Urop SINEs described above, we found approximately 2000 copies of individual Urop monomers in the *U. gracilis* genome (Fig. S1). Most of them (70%) were flanked by target site duplications (TSDs) that occurred after SINE integration into the genome. This indicates they were amplified from monomeric Urop copies and are independent SINEs rather than truncated dimers, which are also abundant in the *U. gracilis* genome. Most likely, it was the fusion of pairs of monomeric Urop that gave rise to retropositionally more active subfamilies of dimeric Urop. The Urop-m consensus is most similar to the two Urop_a monomers, and also closely resembles both pB1 and FLAM (Fig. 2A). A schematic representation of all the above discussed 7SL RNA-derived SINEs is given in Fig. S2.

Figure 2B illustrates the sequence of events (deletions, duplications, and dimerizations) that generated broadly similar 7SL RNA-derived SINEs in the shrew mole, primate, and rodent genomes. The scheme highlights the parallelism in the evolution of 7SL RNA-derived SINEs in three different groups of mammals.

The average copy similarity is 83% for Urop_a and Urop_b and 88% for Urop_c. The distribution of copies by nucleotide substitutions in them for the three subfamilies plotted in Figure 3A indicates the following. (1) The Urop SINE family arose long ago, apparently shortly after the *Uropsilus* lineage diverged from other Talpidae, which occurred about 50 Mya (Springer, et al. 2018). (2) Most Urop copies were integrated into the genome many millions of years ago. (3) The Urop_c subfamily is younger than the other two; judging from Figure 3A, there have been two waves of Urop_c amplification. Thus, the data suggest that Urop represents an old SINE family whose activity peaked long ago. However, the Urop_c subfamily included two fractions, each consisting of approximately 300 copies, whose sequences were 98% similar to their consensus sequences (Fig. S3). These data suggest recent retropositional activity of certain Urop_c copies. In addition, we extracted some Urop_c copies with particularly long pure poly(A) tails (Fig. S4 shows 16 Urop_c copies with A_21─90_). In L1-mobilized SINEs, long poly(A) tails are known to be characteristic of recently integrated and potentially active copies; these poly(A) tails shorten and acquire nucleotide substitutions over time (Roy-Engel, et al. 2002; Odom, et al. 2004; Vassetzky, et al. 2021;Kosushkin, et al. 2022). Thus, Urop_c copies with long pure poly(A) tails have been amplified and incorporated into the genome relatively recently, suggesting the recent activity of Urop_c subfamily.

**Fig. 3.**
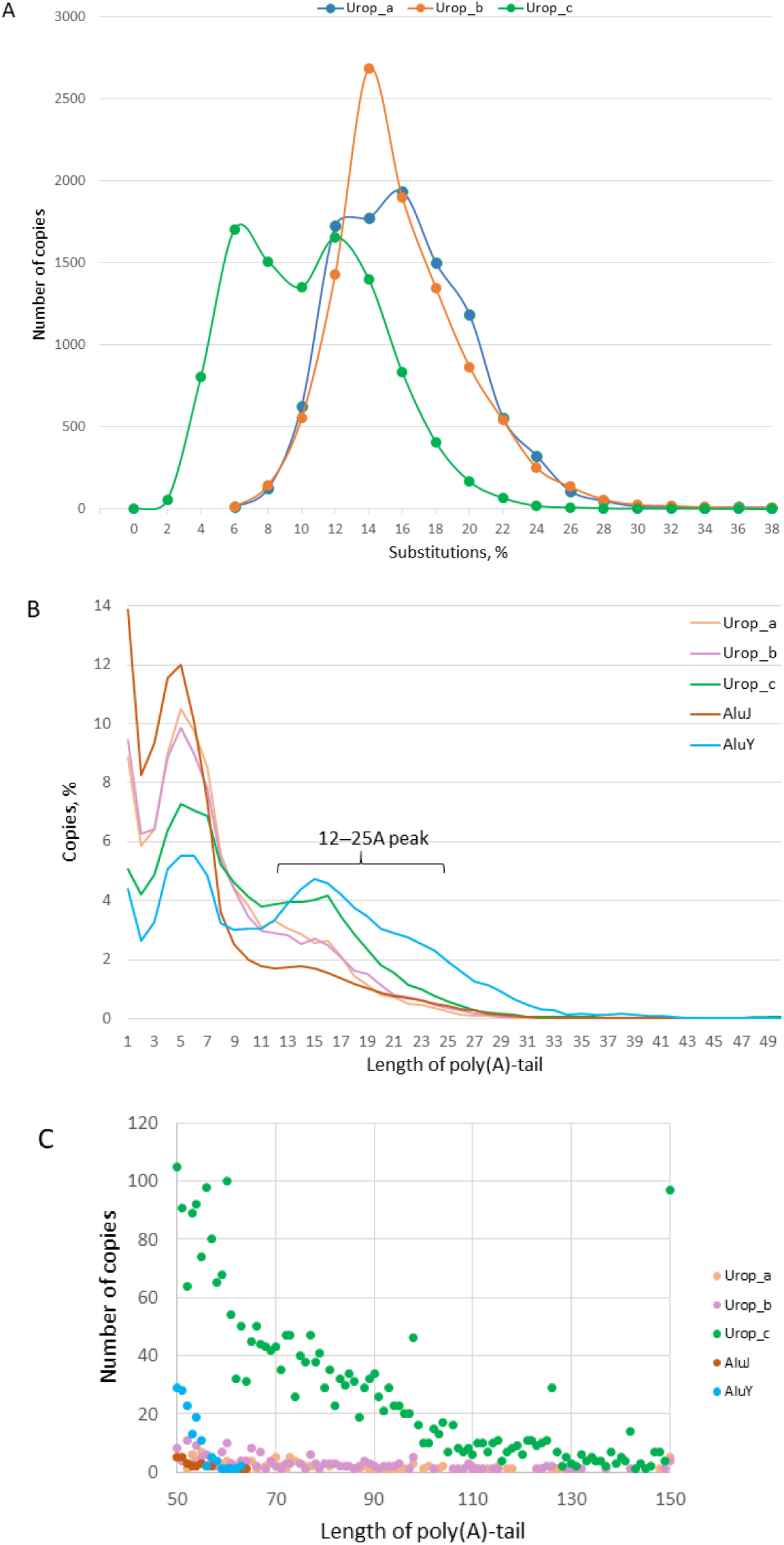
Sequence analysis of all copies of the three Urop subfamilies with a focus on poly(A) tails in the *Uropsilus gracilis* genome. **A**. Distribution of Urop_a, Urop_b, and Urop_c copies based on similarity to the subfamily consensus sequence. **B**. Distribution of copies of Urop_a, Urop_b, and Urop_c by the length of their poly(A) tails. Similar data obtained for the old (J) and young (Y) human Alu subfamilies are shown for reference. **C**. Distribution of Urop_a, Urop_b, and Urop_c copies with extra-long (A_≥50_) tails. Similar data obtained for the old (J) and young (Y) human Alu subfamilies are shown for reference. The point at x=150 shows the number of copies with a tail equal to or longer than A_150_.

We analyzed the poly(A) tails in 149,255 copies of Urop_c and in 123,730 copies of human AluY. Pure 12─25-long A-tails are readily identified in Urop_c similar to the young AluY subfamily, as visualized by the additional peak in the plots (Fig. 3B). This finding confirms the presence of a significant number of young copies in the Urop_c subfamily. Subsequent analysis of pure extralong (A_>50_) tails revealed at least twice shorter pure A-tails in AluY compared to Urop_c, which exceeded 150 bp at several dozen loci (Fig. 3C). Furthermore, the number of Urop_c copies with A_>50_ is much higher (2750) than that of such AluY copies (140). It is worth noting that copies of Urop_a and Urop_b with very long tails (A_50-150_) also occur although in much smaller numbers compared to Urop_c (Fig. 3C). A sample of Urop_c copies with extra-long tails (A_139-178_) is shown in Fig. S5. It is not unlikely that certain aspects of Urop template slippage during DNA replication or reverse transcription underlie such elongation of poly(A) tails relative to Alu. In addition, it is likely that the poly(A) tails of at least some Urop copies shorten much more slowly than those of young Alu, and nucleotide substitutions occur in them at a lower rate. The biochemical processes underlying the differential elongation and shortening of SINE poly(A) tails in *Uropsilus gracilis* and humans remain to be elucidated.

## Discussion

SINEs have evolved throughout the evolution of genomes. When one of the cellular RNAs is reverse transcribed and cDNA integrated into the genome, its genomic copies can gradually acquire properties that make them capable of amplification (Roy-Engel 2012). Despite the extreme rarity of such events, most genomes of higher eukaryotes, especially those of mammals, contain large numbers of SINE copies. The period of activity of SINE families is limited and their taxonomic distribution is restricted to a range of related taxa, from family to order (although relict copies of SINEs active in the ancestors of higher-ranked taxa are not uncommon).

Here we have described the Urop SINE family from the genome of the shrew mole *Uropsilus gracilis*. It is highly likely that Urop is also present in the genomes of a dozen other living species of the genus *Uropsilus*, which constitute the subfamily Uropsilinae of the family Talpidae. The split of the extant *Uropsilus* into species occurred no later than 8 Mya (Wan, et al. 2013), while the lowest estimated age of old Urop copies is 33 My (assuming that the nucleotide substitution rate in *Uropsilus* is as high as in rodents (Gibbs, et al. 2004)). In addition, there are no copies of Tal in the *Uropsilus gracilis* genome, which is a characteristic SINE in other subfamilies of the Talpidae. The data obtained suggests that Urop appeared in the Uropsilinae, while Tal appeared in the common ancestor of the other Talpidae after the divergence of these two lineages (Fig. 1A). To date, this is a unique example of such a contrasting distribution of different SINE types (tRNA- and 7SL RNA-derived) within the same family; this may confirm the elevation of Uropsilinae to a family, as proposed elsewhere (Bannikova 2019).

The finding of a 7SL RNA-derived SINE so similar to human Alu in a very distant mammal is surprising. Previously, 7SL RNA-derived SINEs have been described in supraprimates (e.g., Alu in primates, B1 in rodents, and Tu types I and II in tree shrews), all of which share a common ancestry. Another example, Ebu1-1 in hagfish, only underscores the extraordinary origin of SINEs from 7SL RNA.

Unlike most other 7SL RNA-derived SINEs, all three Urop subfamilies (a, b, and c), like Alu, are composed of two 7SL RNA-derived monomers joined by an A-rich linker. The right and left monomers in both Alu and Urop are similar but not identical (except for the Urop_a monomers).

The 183-nt deletion in the central part of the 7SL RNA that preceded the formation of the left Urop monomer is nearly identical to the deletion in all successful 7SL RNA-derived SINEs (and the 3’ boundary is the same even in the very distant hagfish Ebu1-1) (Fig. 2). This striking similarity of deletions that occurred in completely different lineages suggests that the molecular mechanisms of their generation and/or functional selection are the same. Perhaps these deletion boundaries are determined by the secondary structures of the 7SL RNA. Under this premise, the boundaries of the 183-nt deletion (marked with red arrows in Fig. S6) lie side by side in the folded 7SL RNA. These regular, albeit extremely rare, deletions in the 7SL RNA have probably allowed the emergence of amplification-competent SINEs.

Although Alu is probably the best known example of a SINE, independent lineages of 7SL RNA-derived SINEs are extremely rare compared to SINEs of other origins: to date, such SINEs have only been identified in supraprimates and hagfish. Moreover, mammals are the best-studied class with respect to SINEs. In this context, the discovery of an independent 7SL RNA-derived family of SINEs was very unexpected.

We believe that the similarity of Alu and Urop SINEs is due to their independent origin from 7SL RNA in primates and shrew moles rather than to horizontal SINE transfer between these mammals. Unlike other mobile genetic elements, there is no clear evidence for horizontal transfer of SINEs. Although the possibility of SINE horizontal transfer has been suggested several times (Kordis and Gubensek 1995; Piskurek and Okada 2007; Han, et al. 2021), subsequent studies have seriously questioned it (Gogolevsky, et al. 2008; Fawcett and Innan 2016; Platt, et al. 2018).

Overall, the independent emergence of dimeric 7SL RNA-derived Alu and Urop is a vivid example of parallel evolution. Interestingly, higher primates and shrew moles have only 7SL RNA-derived SINEs and lack specific tRNA-derived SINEs. (Relict SINEs such as Ther1 and Ther2 are not considered.) At the same time, the vast majority of mammalian species have one or more families of tRNA-derived SINEs in their genomes. In rodents, in addition to the 7SL RNA-derived SINE B1,there are tRNA-derived SINEs ID and B2 (the latter is only characteristic of mouse-like rodents). It is possible that the high retropositional activity of the dimeric 7SL RNA-derived SINEs Alu and Urop prevented the appearance and spread of tRNA-derived SINEs in the genomes of higher primates and shrew moles.

By analyzing the level of nucleotide substitutions in Urop copies relative to the consensus sequences of the subfamilies, we concluded that Urop_a and Urop_b are older than the Urop_c subfamily. Among Urop_c, we found groups with the most similar (98%) nucleotide sequences, indicating their relatively recent integration into the genome (about 2.5 Mya); it is possible that such copies are still retropositionally active. Long poly(A) tails in L1-mobilized SINEs indicate their relatively recent integration into the genome and possible retropositional activity (Roy-Engel, et al. 2002; Odom, et al. 2004; Vassetzky, et al. 2021). We have shown a remarkable fraction of copies with rather long poly(A) tails (A_13─25_) among Urop_c, as in the young AluY subfamily. Odom *et al*. (2004) suggested that the most retropositionally active Alu copies should have extra-long (A_>50_) poly(A) tails. Unexpectedly, there were 20 times more copies with extra-long poly(A) tails among Urop_c than among AluY; moreover, the tail length of such Urop_c was at least twice that of AluY. Such long poly(A) tails in certain Urop copies may indicate specific mechanisms controlling poly(A) tail elongation and shortening in *Uropsilus gracilis*.

In conclusion, 7SL RNA-derived SINEs emerged independently in three lineages: supraprimates, shrew moles, and hagfish. In *Uropsilus*, SINE evolution likely involved the following events. A long deletion in the central part of the 7SL RNA produced a functional but ineffective Urop-m SINE. Its dimerization resulted in a much more efficient retrotransposon. An internal duplication within the right monomer and minor modifications produced the most efficient Urop subfamily. While Urop retrotransposition peaked in the past, not many Urop copies remained active until recently. Similar events (and hence the same underlying mechanisms) but not identical scenarios in two other lineages led to the emergence of 7SL RNA-derived SINEs (in particular, primate Alu with essentially the same structure). These findings open new avenues for understanding the evolution of SINEs.

## Methods

Genome assemblies of *Uropsilus gracilis* (UroGra_v1_BIUU), *Talpa occidentalis* (MPIMG_talOcc4v2.1), *Galemys pyrenaicus* (Gpyr_1.0), *Condylura cristata* (ConCri1.0), *Scalopus aquaticus* (ScaAqu_v1_BIUU), and *Homo sapiens* (T2T-CHM13v2.0) were downloaded from NCBI Genomes (https://www.ncbi.nlm.nih.gov/datasets/genome/). Copies of the SINE were found using the ssearch36 program from the 3.6 release of the FASTA package (Pearson and Lipman 1988) with at least 65% identity and 90% length overlap with the query sequence. AluY sequences were extracted from the human genome using the RepeatMasker annotations (https://hgdownload.soe.ucsc.edu/goldenPath/hs1/bigZips/).

Sequence manipulations were performed using SeqKit (Shen, et al. 2016) and standard Linux programs. BED genomic intervals were operated using bedtools v2.31.0 (Quinlan and Hall 2010). Multiple sequence alignments were generated using MAFFT (Yamada, et al. 2016) and edited using GeneDoc (http://www.nrbsc.org/gfx/genedoc/index.html). Consensus sequences were generated from multiple sequence alignments in GeneDoc or using the cons program from the EMBOSS package (http://emboss.open-bio.org/html/use/pr02s04.html). The mean similarity to the consensus sequence was determined for 10000 randomly selected sequences from each subfamily using the esl-alistat program (hmmer.org). The A-tail lengths were determined by alignment of a SINE sequence with the corresponding consensus without tail and counting the length of the downstream pure poly(A) sequence.

RNA secondary structure folding was performed on the mfold server using the default settings http://www.unafold.org/mfold/applications/rna-folding-form.php.

## References

Bannikova A. 2019. Molecular evolution and the problems of phylogenetic reconstruction of true insectivores (Mammalia: Eulipotyphla). Doctoral Thesis, Moscow State University.

Borodulina OR, Kramerov DA. 2001. Short interspersed elements (SINEs) from insectivores. Two classes of mammalian SINEs distinguished by A-rich tail structure. Mamm Genome 12:779–786.

Chen C, Ara T, Gautheret D. 2009. Using Alu elements as polyadenylation sites: A case of retroposon exaptation. Mol Biol Evol 26:327–334.

Chen JM, Ferec C, Cooper DN. 2006. LINE-1 endonuclease-dependent retrotranspositional events causing human genetic disease: mutation detection bias and multiple mechanisms of target gene disruption. J Biomed Biotechnol 2006:56182.

Deininger P. 2011. Alu elements: know the SINEs. Genome Biol 12:236.

Deininger PL, Jolly DJ, Rubin CM, Friedmann T, Schmid CW. 1981. Base sequence studies of 300 nucleotide renatured repeated human DNA clones. J Mol Biol 151:17–33.

Fawcett JA, Innan H. 2016. High Similarity between Distantly Related Species of a Plant SINE Family Is Consistent with a Scenario of Vertical Transmission without Horizontal Transfers. Mol Biol Evol 33:2593–2604.

Ferrigno O, Virolle T, Djabari Z, Ortonne JP, White RJ, Aberdam D. 2001. Transposable B2 SINE elements can provide mobile RNA polymerase II promoters. Nat Genet 28:77–81.

Gibbs RA, Weinstock GM, Metzker ML, Muzny DM, Sodergren EJ, Scherer S, Scott G, Steffen D, Worley KC, Burch PE, et al. 2004. Genome sequence of the Brown Norway rat yields insights into mammalian evolution. Nature 428:493–521.

Gogolevsky KP, Vassetzky NS, Kramerov DA. 2008. Bov-B-mobilized SINEs in vertebrate genomes. Gene 407:75–85.

Han G, Zhang N, Jiang H, Meng X, Qian K, Zheng Y, Xu J, Wang J. 2021. Diversity of short interspersed nuclear elements (SINEs) in lepidopteran insects and evidence of horizontal SINE transfer between baculovirus and lepidopteran hosts. BMC Genomics 22:226.

Kojima KK. 2020. Hagfish genome reveals parallel evolution of 7SL RNA-derived SINEs. Mob DNA 11:18.

Kordis D, Gubensek F. 1995. Horizontal SINE transfer between vertebrate classes. Nat Genet 10:131–132.

Kosushkin SA, Ustyantsev IG, Borodulina OR, Vassetzky NS, Kramerov DA. 2022. Tail Wags Dog’s SINE: Retropositional Mechanisms of Can SINE Depend on Its A-Tail Structure. Biology (Basel) 11.

Kramerov DA, Vassetzky NS. 2011a. Origin and evolution of SINEs in eukaryotic genomes. Heredity (Edinb) 107:487–495.

Kramerov DA, Vassetzky NS. 2011b. SINEs. Wiley Interdiscip Rev RNA 2:772–786.

Krayev AS, Kramerov DA, Skryabin KG, Ryskov AP, Bayev AA, Georgiev GP. 1980. The nucleotide sequence of the ubiquitous repetitive DNA sequence B1 complementary to the most abundant class of mouse fold-back RNA. Nucleic Acids Res 8:1201–1215.

Kriegs JO, Churakov G, Jurka J, Brosius J, Schmitz J. 2007. Evolutionary history of 7SL RNA-derived SINEs in Supraprimates. Trends Genet 23:158–161.

Krull M, Brosius J, Schmitz J. 2005. Alu-SINE exonization: en route to protein-coding function. Mol Biol Evol 22:1702–1711.

Nishihara H, Terai Y, Okada N. 2002. Characterization of novel Alu- and tRNA-related SINEs from the tree shrew and evolutionary implications of their origins. Mol Biol Evol 19:1964–1972.

Odom GL, Robichaux JL, Deininger PL. 2004. Predicting mammalian SINE subfamily activity from A-tail length. Mol Biol Evol 21:2140–2148.

Pearson WR, Lipman DJ. 1988. Improved tools for biological sequence comparison. Proc Natl Acad Sci U S A 85:2444–2448.

Piskurek O, Okada N. 2007. Poxviruses as possible vectors for horizontal transfer of retroposons from reptiles to mammals. Proc Natl Acad Sci U S A 104:12046–12051.

Platt RN, 2nd, Vandewege MW, Ray DA. 2018. Mammalian transposable elements and their impacts on genome evolution. Chromosome Res 26:25–43.

Quinlan AR, Hall IM. 2010. BEDTools: a flexible suite of utilities for comparing genomic features. Bioinformatics 26:841–842.

Roy-Engel AM. 2012. A tale of an A-tail: The lifeline of a SINE. Mob Genet Elements 2:282–286.

Roy-Engel AM, Salem AH, Oyeniran OO, Deininger L, Hedges DJ, Kilroy GE, Batzer MA,Deininger PL. 2002. Active Alu element “A-tails”: size does matter. Genome Res 12:1333–1344.

Shen W, Le S, Li Y, Hu F. 2016. SeqKit: A Cross-Platform and Ultrafast Toolkit for FASTA/Q File Manipulation. PLoS One 11:e0163962.

Springer MS, Murphy WJ, Roca AL. 2018. Appropriate fossil calibrations and tree constraints uphold the Mesozoic divergence of solenodons from other extant mammals. Mol Phylogenet Evol 121:158–165.

Vassetzky NS, Borodulina OR, Ustyantsev IG, Kosushkin SA, Kramerov DA. 2021. Analysis of SINE Families B2, Dip, and Ves with Special Reference to Polyadenylation Signals and Transcription Terminators. Int J Mol Sci 22.

Vassetzky NS, Kramerov DA. 2013. SINEBase: a database and tool for SINE analysis. Nucleic Acids Res 41:D83–89.

Vassetzky NS, Ten OA, Kramerov DA. 2003. B1 and related SINEs in mammalian genomes. Gene 319:149–160.

Veniaminova NA, Vassetzky NS, Kramerov DA. 2007. B1 SINEs in different rodent families. Genomics 89:678–686.

Wan T, He K, Jiang XL. 2013. Multilocus phylogeny and cryptic diversity in Asian shrew-like moles (Uropsilus, Talpidae): implications for taxonomy and conservation. BMC Evol Biol 13:232.

Yamada KD, Tomii K, Katoh K. 2016. Application of the MAFFT sequence alignment program to large data-reexamination of the usefulness of chained guide trees. Bioinformatics 32:3246–3251.

