## Supplementary material for "A Dimeric SINE Discovered in Shrew Mole is Structurally Similar to Primate Alu": Fig. S1

```
                                                      *        20         *        40         *        60         *        80         *       100         *       120         *
Urop-m                           :                              AGCCGGGCRTGGTGGCGCGCGCC----TGTAATCCCAGCTACC-TGGGAGGCT-----GAGGCTAGGGATC---ACTTGAGTCCGY-GGGTTCGAGACTACC
PVHY01015140.1_34556-34880       : GCACATTAGACTCACAAAGTTTCATACCAGCTGGGTGCAGTGGTGCACGCC----TGTAATCCCAGCCACC-TGGGAGGCT-----GAGGCTAGGGATC---ACTTGAGTCTGT-GGGTTCGGAACTAAC
PVHY01031403.1_6020-6344         : TGAGATTCCTCTTTAAAAGTCATATAGGAGCCGGGCATGGTGGTGCATGCC----TGTAATCCCAGTTACC-CGGGAGGCT-----GAGGCTGGGGATC---ATTTGAGTCCGT-GGGTTCCAGACAACC
PVHY01000151.1_192297-19262      : TTTCCTTTTCAAAAAAAAAAATTATTATTGCCAGGCATAGTGGTGTGCGCC----TGTAATCCCAGCTACC-TGGGAAGCT-----GAAGCTAGGGATC---ACTTGAGTCCAT-GGGTTCAAGACAACC
PVHY01004557.1_27327-27651       : TTTGTTAAGCTAAAGAAACAGGAAAAAAAGCCAGGCATGGTGGCACGCCCC----TATAATCCTAGCTACC-TGGGAGGCT-----GAGGCTAGGGATC---ACTTGAGTTTGC-AGGTTTGGGACTACC
PVHY01003029.1_38740-39064       : AACAGGACTATGAAAAGTAGGGAGTATGAGTCAGGCATGGTGGTGTGTGCC----TGTAATCCCAACTACC-TGGGAGGCT-----GAGGCAAGGGATC---ACTTGGGTCCAT-GGGTCCAAGACAACC
PVHY01000107.1_284017-28434      : ATAAATAAGGACATTTAAGTGACCCTGAAACTGGGTGTGGTGGCTCGTATC----TGTAATCCCAGCTACC-TGGGAGGCT-----GAGACTAGGGATC---ACTTGAGTCCAC-AGGTTCAAGACTACC
PVHY01021119.1_21558-21881       : AATTTTTTTAAAAATAAATTTTATTTGGAGTCGGAAGTGGTGGCTGCGTC----TGTAATCCCAGCTACC-TGGGAGGCT-----GAGACTAGGGATC---ACTTGAGTCCGC-GGGTTCAAGAAAACC
PVHY01004698.1_89996-90320       : TTATTCATAATAAAAAAAATGTATTCATAGCCAGGCATGGTGGCACACACA----TGTACTCCCAGCTACC-TGGGAGGCT-----GAGGCTAGGGATC---ACTTGAGTCCAC-AGGTTTGAGACAACC
PVHY01009889.1_9618-9942         : TGAAACTGAGTCAAGATTTAATAAAATTAGCTGGGTGCGCTGGTGTGTGCC----TGTAATCCCAGTTACC-TGGGAGGCT-----GAGGCTGGGGATC---ACTTGAGTCCAT-GGGTTTGAGATAACC
PVHY01007153.1_59040-59364       : AAATGAGAGTAGCACTCAAGAGACTATAAGCCGGGTGTGGTGGCACACACC----TGTAATTCTAGCTACCTTGGGAGGTT-----GAGGCTAGGGATC---ACTTTAATTCGC-AGGTTCAAAACTACC
PVHY01015370.1_10638-10960       : TAATTCATTTTTCAAAACAACTGAAAGTAGCTGGGCATGATGGTGCATGCC----TGTAATCCCAGCTACT-TGGGAGGCT-----GAGGCTAGAAATC---ACTTGATTCCGC-AGGTTCGAGACTACC
PVHY01000416.1_6821-7140         : GGAGAGCAATTGATTTCAATCTCTATTCTGGTAGACATGGTGGTGCACACC----TGTAATCCCAGCTAGC-TGGGAGGCT-----GATGTTGGGGATC---ACTTGAGTCTCC-GGGTTCGAGACTATT
PVHY01003821.1_68789-69114       : AAGACTTCATTTTAAAAAGTCAGAATAAAGCTGGGCGCGGTGGTACACACC----TGTAATCCCGGCTACC-TGGGAGACT-----GAGGCTAGGGATC---ATTCGAGTCCATGGGTTCGAGACAACC
PVHY01000470.1_10163-10473       : TACATTAGTTGCAAATAATCTGAAATCCAACAAGACATGGTGGCACATGCC----TATAATCCCAGTTACC-TGGGAGACT-----GGGGCTGAGGATC---ACTTGAGTCCAT-GGGCTCAAGACAACC
PVHY01005138.1_68052-68367       : GTATCAGTTTAAAAGTTTCTCTCTGCTAATCTAGGTATGGTGGTGCACGCC----TGTAATCCCAGCTACC-TGGGAGACT-----GAGGCTAGGGATC---ACTTGAGTCCAT-GGGTTCGAAATAA-C
PVHY01016211.1_26091-26418       : TGTACAATCAAGTAAAAAAAAAAAAGCTAGCCGGGCGTGGTTGCACGTGCC----TGTAATCCCAGCTACC-TGGGAGGCT-----GAGGCTAGGGATC---ATTTGAGTCTGC-GGATTTGGGACTA-T
PVHY01003071.1_16040-16363       : ATAAATAAAAGTTAAAATTCCCATATAAAGTCAGGCATGGTGGTGTGTGCC----TGTAATCCCAGTTACC-TGGGAGGCT-----GAAGTTGGAAATC---ACTTGAGTCCAC-GGATTTGAGACAACC
PVHY01012321.1_2445-2767         : TCCCCCTCACCCTGAAAAATGACTATGTAACCAAGCATGGTGGCACATGCC----TGTAATCCCAGCTACC-TGGGAGGCT-----GAGGTTAGGGATC---ACTTGAGTCAGT-GGGTTTGAGACAG-C
PVHY01008062.1_45206-45521       : TATTTGTGCTTTGTCTATAATTATATCTTATAAGGCATGGTGGCATGCACC----TGTAATTCCAGTTACC-TGGGTGACT-----GAGACTGGGTATC---ACTTGGGTCTGA-GGGTTCAAGACAATC
PVHY01000361.1_41634-41961       : AGAATATAACATAAAAACATCAAGACCAGGCCAGGTGTGGTGGCACATGCCACCACATAATCCCAGTTACC-TGGGAGGCT-----GAGGCTGGGGATC---ACATGAGTCCGT-GGTTTTGAGGCAACC
PVHY01002065.1_106129-10644      : CTAACATCTCCATAAAATATGTAGAACTAGCCAGGCATGGTGGCA-GCACC----TGTAATTCCAGTTACC-TGGGAGGTT-----GAAGCTATGAATC---TCTTGAGTCCAT-GGGTTTGAGACAACC
PVHY01000597.1_141223-14154      : TTGCTCCCTTAAAAAGTTACAACTTGCAAAGCTGGTGTGGTGGTGTGCACC----TGTAATCCCAGTTACC-TGGGAGACT-----GGGGATGGGAATC---ACTTGATTCCGT-GGGTTCAAGACAACC
PVHY01035533.1_6734-7055         : TTTTCAATATCTCTTTAAATACTACTCCTGCCGGGCACGATGGTGTGCACC----TGTAATCCCAGCT--C-TGGGAGGCT-----GAGGCCAGGGATC---GCTTGAGTCTGC-AGGCTCTAGACTACC
PVHY01000497.1_225978-22630      : CAGTGGATGTATAAGATTTCAAAATTAAGGCCAGGCATGATGGTGCATGCC----TGTAATCACAG-TACC-TGGGAGGCT-----GAGGCTCAGTATC---ACTTGAGTTTGT-AGGTTGAAGTCAACC
PVHY01023319.1_6386-6705         : TGCCAACAAAATAAAAAGTCAGTTAGCAAGCTGGGAGCGGTGGTGTGTGCC----TGTAATCCC----ACC-TGGGAGGCT-----GAGGCTAGGAATC---ACTTGAGTCTGC-GGGTTCAAGACAACC
PVHY01001968.1_95636-95960       : GAAATTTATATATTAAAAAAAAAAAAAGCAGCTTGACGCGATGGTGGGCACC----TGTAATCCCAGTTACC-TGGGAGGTT-----GATGCTCGGGATC---GCTTGAGTCTGT-GGGGTTGGGACTACT
PVHY01000724.1_186353-18667      : AAAACAAATCAATTAGTTAAAAAAATTAGGGGTGGGCATGGTGGTGTATGCC----TGTAATCCCAGT--AC-TGGGAGGCT-----GAGGCCAGGGATC---ACTTGAGTCTGC-AGGTTTAGCACTACC
PVHY01017384.1_23229-23550       : GCAATATTTAAGAATTTTTTTCAGTCTAAGCCGGAAGTGGTGGTGGGTGCC----TGTAATCCC-TCTACC-TGGGAGGCT-----CAGGGTAGGGATC---ACTTGAGCCCGC-GGGTTCGAGACAAAC
PVHY01023758.1_7684-8006         : AGGGCAATAGGGAGTTAAAGTAAAGCTGAGCTGGGTGTGGTGGTGTGTACC----TGTAACCCCTGC--CT-TGGGAGGCT-----GAGGCTAAGGATC---GCTTGAGTCTGC-GTGTTTGGGACTACT
PVHY01013372.1_14164-14486       : TGACTTTTGTAGCTTTTAAAAGCAAGCTCGCTGGGCATGGTGGCAAGTGCC----TATAATCCCACCTACC--TGGAGGCT-----GAGATTAGGGATC---GCTTGAGTCTGT-GGGTTTCAGACTACC
PVHY01020839.1_22156-22479       : AATATTAAAAAACAAAATGACAAAAACAAGCTGGGTGCAGTGGTGAGCACC----TGTAATCCCAGTTAAA-TGGGAGGCT-----GAGGCTAGAAATC---ACTTGAGTGTGC-AGGGT-GAGACAACC
PVHY01011840.1_1777-2099         : GAAGTTCTTTGGCAATAACCATTAGGAAAGTTTGGCACAGTGGCGGGTGCC----TGTAATCCCAGCTACC-TGGGAGGCA-----GAGGTTAGGGACC---ACTTGAGTCTGC-AGGTTCCAGACAACC
PVHY01012998.1_32215-32544       : ATTTCCAAATGAAGATAAACATGCTATGAGCTGGGCATGGTGGTACACATC----TGTAATCCTAGTTACC-TAGAAGGTTACCTAGAAGGCTGGGATC---ACTTGAGTCTGT-GGGGTCGAGACAAGC
PVHY01048485.1_2716-3038         : GTAGTCATGGGTTTAAAATGGGTGAACTTGCCGGGCGTGGTGGCACGCACC----TATAATCCAAGC-ACT-GGGAGGCAA-----GAGGCCAGGGATC---ACTTGAGTTTGC-AGGTTCAAGGTCACC
PVHY01165340.1_247-508           : GAGCCCAGATAAAGAGTGAAAATTCCTATGCCAGGCGGGACGGCGCACGCC----TGTAATCCAAGCAC---TGGGAGGCC---AAGAGGCCTGGGATC---GCTTGAGTTCGC-AGATTCAAGACTACC
PVHY01038621.1_7263-7582         : GGGCATCATCGTCTAAAGATGGGGCTCAAGCCGGGTGTGGTGGCACGTGCC----TATAATCCAAG-TGCT-GGGAGGCAA-----GAGGCCTGGAATC---ACTTGAGTTCTT-GGGTTCGAGATGATC
PVHY01014257.1_20943-21266       : AAAAAATTACTTCTTAAAAATGAACTTCAGTCAGATGTGGTGGTACACGT-----AGTAATCCCAGCTGCC-TGGAAGGCT-----GAGGCTACGGCTC---ACTTGAGCCCAC-GCGTTTGAGACAACC
PVHY01000251.1_41103-41427       : TTAAGTGCACAAAAGAGAATACGGAAGTAGTTGAGCGTGGTGGCACATGCC----TGTGATTCCAGCTACTTGGAGAGGCT-----GAGGCTAGGGATC---ACTTGAGTCCAC-GGATTCAAGAAAACC
PVHY01291323.1_18-247            :                 TCCACAAAAATAGCTGGGCGTGGTGGCATGTGCC----TGTAATCCCAGCTACC-TGGGAAGAT-----GAGGCTGGGGACTGCAACGAGAGACTGT-GGTTCCATGGCA---
PVHY01074865.1_1107-1420         : ACTAATAGTTCAGTTTCTGTGGATAGAAACCCAGGTT---CAGCGTGTGTC----TGTAATCCCAGCTGTT-TGGGAAACT-----GAGGCTAGGAATCACCGGTTGAGTCCGC-GGGTTGGAAACTACC
PVHY01007523.1_58227-58550       : ATACATATTCATATTAAGATATCAAATAACCCAGGAGTGATGGTGTGCACC----TGTATTCCAACCT-CT-GGGAGGTAA-----GGGGCCTGGGATT---GCTTGAATTCAT-GGGTTCAAGACTAGC
PVHY01026174.1_2214-2532         : GCATATTTTAAAATGATCTTCCTGCAAATGCTGATCATGGTGGTGCACGCT----TGTAATCGCAGTTACC-TCTGG---------GAGGCTGAGGATC---ACTTGAGTCTGC-GGGTTCAAGACAA-C
PVHY01054549.1_2422-2727         : AAAACCAAGGATTAATGAGACCTAGTCTCAAAATATAGTCAGGTGCGTGCC----TGTAATCCCAGCTACC-TGG-----------AAGGCTGGGGATC---ATTTGAGTCCGC-GGGGTCAGGACTACC
PVHY01013156.1_31237-31558       : TTAAGTGCTTGTTAATTTATACAAAACTACCTGGGTGCAGTGGCATGAGCTT-----TAATCCAAGCCAC--TGGGAGGCT-----GAGGCTAGGAATC---ACTTGAGTCTGT-GGGTTTGAGACAACC
PVHY01020883.1_9691-10008        : CAAGGCAACTCATAATCAAGAGAACATTCTAAGGGCCTGGTGGCGTGGCC------TAATCCAAG-TGCT-AGGAGGCAA-----GAGTACAGGGATC---GCTTAAGTTTGC-AGGTTCAAGATTTCC
PVHY01033073.1_6531-6856         : TCTATAAAGATAAAAATCATGTATTCAAGCCAGGGCACAGTAGCTTGCACC----TGTAATCCCAACTATT-TGAAAGACT-----GAGACTGAGGCTAG------GCATCAGC-AGGTTCCAGACTATA
PVHY01017490.1_12515-12829       : GTAAAGCAAAGATAGAAAGAGAATTAGAGGCCGGGCATGGTGGCGCATGCC----TGTAA--GCGGCAGCC-TAGGAAGCT-----GTGACCAGG-------GCCAGGGATCAC--AGTTTGAGTCCCAC
PVHY01068994.1_1295-1605         : AACCAGCTCTGTTCAACAGTGGCGTGCCGGCCGGGGGCCAGGCGCGCGCC----TGTCATCCCAG-TGCC-CGGGAGGCT-----GAGGCTGGGG------TCGCGTGGTCGC-GGGCTGGAGGCTGGC
```

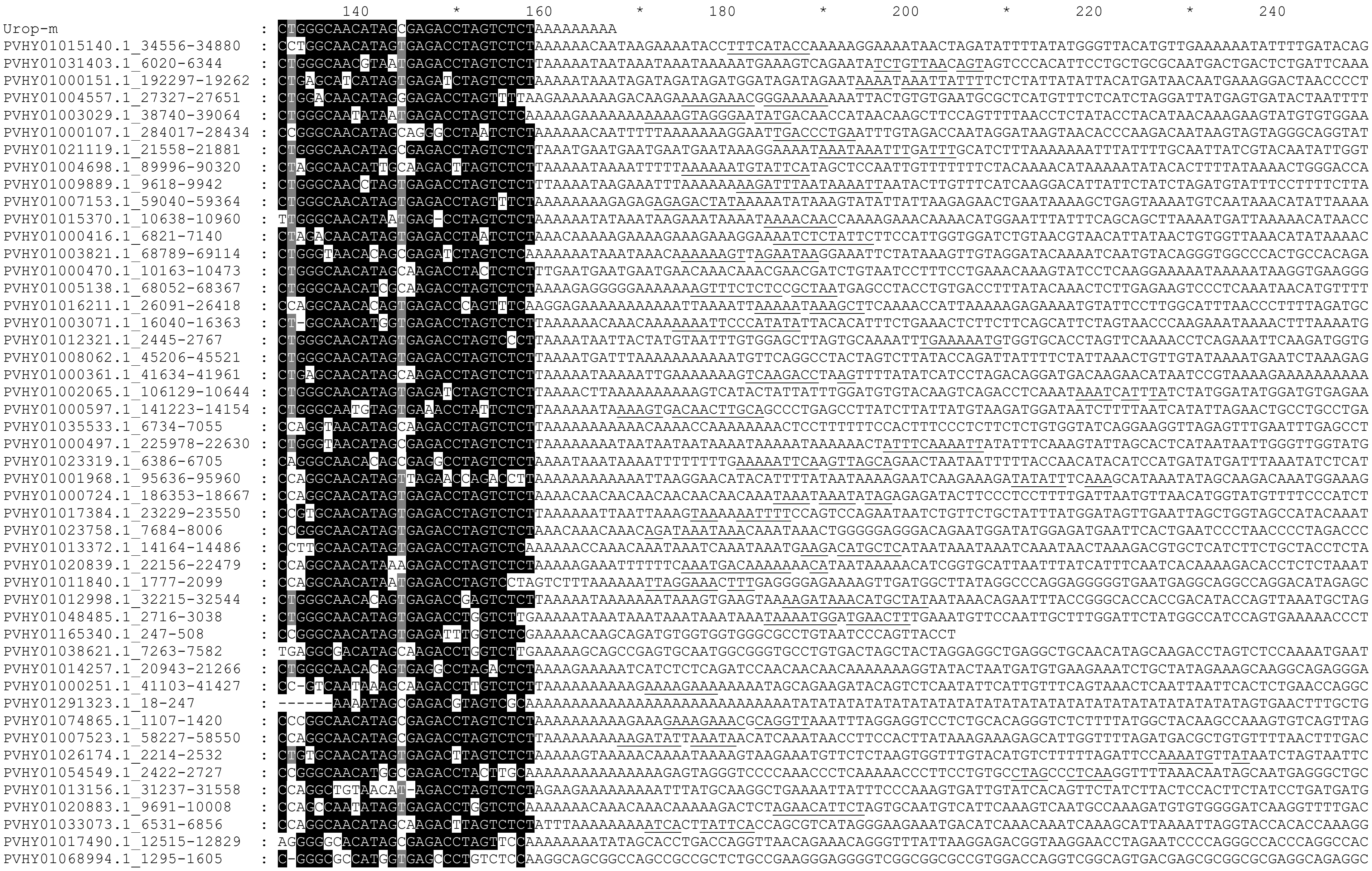

**Figure S1.** Examples of free monomer Urop elements (Urop-m) from the genome of *Uropsilus gracilis*. TSDs are underlined.
