## Supplementary material for "A Dimeric SINE Discovered in Shrew Mole is Structurally Similar to Primate Alu": Fig. S2

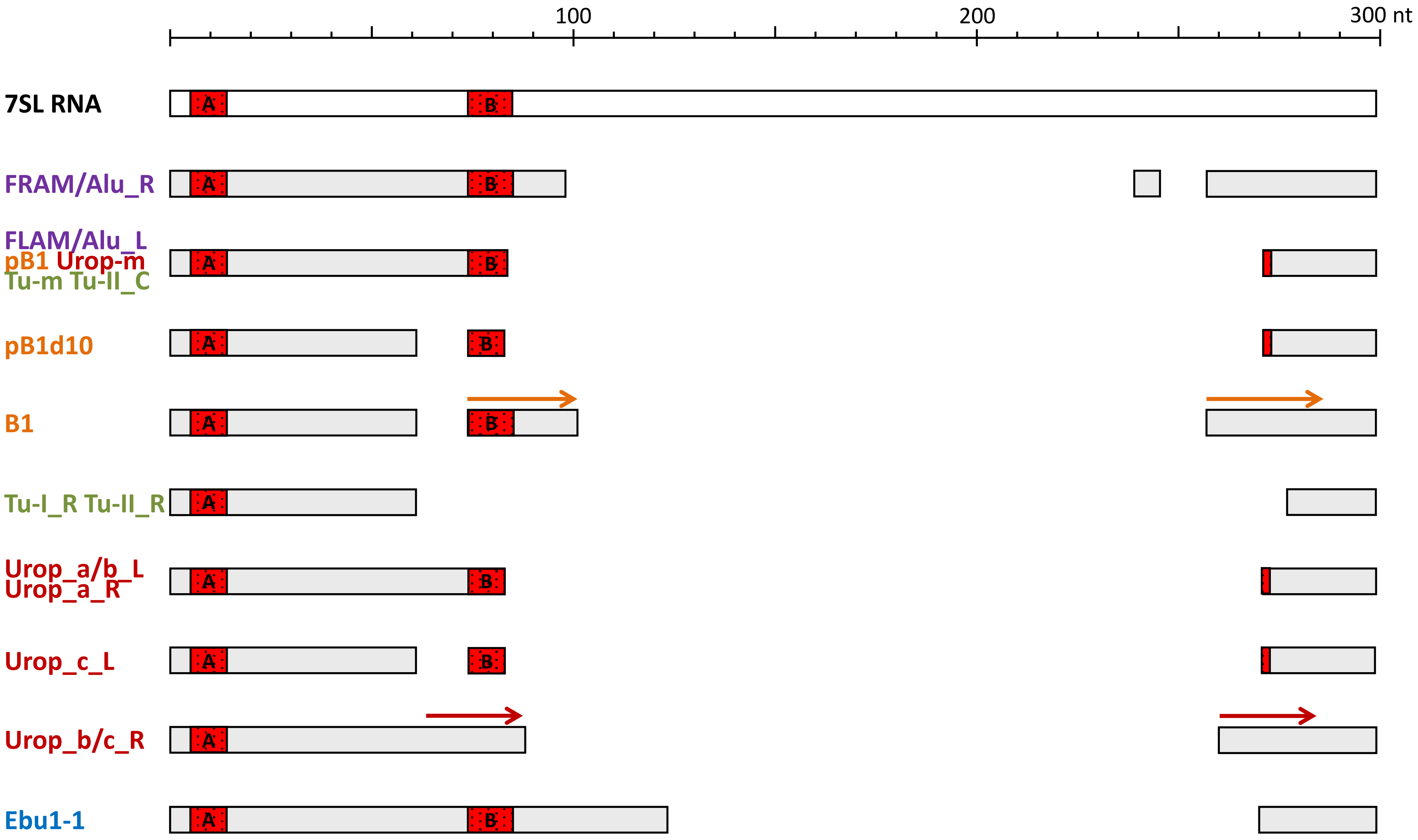


**Figure S2.** A schematic of the 7SL RNA-derived SINEs discussed in the study. 7SL RNA and SINEs are shown as rectangles. Gaps between rectangles correspond to 7SL RNA regions deleted in SINEs. Boxes A and B of pol III promoters are highlighted in red. Direct repeats (duplications) in B1, Urop_b_R and Urop_c_R are marked with arrows. The following SINEs or their monomers are shown in the scheme. Human: FLAM, FRAM, Alu_L, and Alu_R. Mouse: pB1, pB1d10, and B1. Tree shrew: Tu-m, Tu-I_R, Tu-II_C, and Tu-II_R. Shrew mole *Uropsilus gracilis*: Urop-m, Urop_a_L, Urop_b_L, Urop_c_L, Urop_a_R, Urop_b_R, and Urop_c_R. Hagfish *Eptatretus burger*: Ebu1_1.
