## Supplementary material for "A Dimeric SINE Discovered in Shrew Mole is Structurally Similar to Primate Alu": Fig. S3

```
                                                      *       20        *       40        *       60        *       80        *      100        *      120
Urop_c tribe 1                 :                                       AGGCCGGGCACGGTGGCTCGTGCTTGTAAGCTTCAGCCTATGAAGCGGCCGGGGC-TAGGGATCGCGGTTCGAGTCCCGCCT
PVHY01002401.1:48683-49143(+)  : AGAAGAAACATAATTAACAAATAAAAAGTAATAAAACTAGGCCGGGCACGGTGGCTCGTGCTTGTAAGCTTCAGCCTATGAAGCGGCCGGGGC-TAGGGATCGCGGTTCGAGTCCCACCT
PVHY01007456.1:45401-45875(+)  : ATTTTGAATACTATGTTGATTATAGAATATCTAGAACAAGGCCGGGCACTGTGGCTCGTGCTTGTAAGCTTCAGCCTATGAAGCGGCCGGGGC-TAGGGATCGCGGTTCGAGTCCCGCCT
PVHY01006771.1:32287-32748(+)  : TAAAAATAGTTGATATCAATTCAAAGAAGGTCAATTCAAGGCCGGGCACGGTGGCTCGTGCTTGTAAGCTTCAGCCTATGAAGTGGCCGGGGCTTAGGGATCGCGGTTCGAGTCCCGCCT
PVHY01045252.1:4788-5249(-)    : AATGTCCTCCCTCTTCTTTGCATTCAAACTCATCCCTAAGGCCGGGCACGGTAGCTCGTGCTTGTAAGCTTCAGCCTATGAAGCGGCCGGGGCTTAGGGATCGCGGTTCGAGTCCCGCCT
PVHY01007539.1:38627-39087(+)  : TCAACAATTTCATAAACACATTTAAAAATTCAATTTAGAGGCCGGGCACGGTGGCTCGTGCTTGTAAGCTTCAGCTTATGAAGCGGCCGGGGC-TAGGGATCGCGGTTCGAGTCCCGCCT
PVHY01037078.1:1316-1776(-)    : AACAGATTTCATACAAAGGCTCAAAAATCAGAACACTGAGGCTGGGCACGGTGGCTCATGCTTGTAAGCTTCAGCCTATGAAGCGGCCGGGGC-TAGGGATCACGGTTCGAGTCCCGCCT
PVHY01013704.1:32430-32891(-)  : CCTCCCTTCCACCCCCAACAAGAACTAGTTAATTTTTTAGGCCGGGCACGGTGGCTCGTGCTTGTAAGCTTCAGCCTATGAAGCGGCCGGGGCTTAGGGATCGCGGTTCGAGTCCCGCCT
PVHY01000998.1:68954-69416(-)  : ATGTTAATTATGTTTATTTAGATTAAAAAATGAAACACAGGCCGGGCACGGTGGCTCGTGCTTGTAAGCTTCAGCCTATGAAGCGGCTGGGGC-TAGGGATGGCGGTTCGAGTCCCGCCT
PVHY01015967.1:4675-5136(-)    : TGGACATTTTGAGCAGCACTACAAAAAATATGATCACTTGGCCGGGCACGGTAGCTCGTGCTTGTAAGCTTCAGCCTATGAAGCGGCCGGGGCTTAGGGATCGCGGTTCGAGTCCCGCCT
PVHY01008490.1:9516-9977(-)    : CGCACACTTGGTGCTGAACAATTAAGAAATAGGAAAAGAGGCCGGGCACGGTGGCTCGTGCCTGTAAGCTTCAGCCTATGAAGCGGCCGGGGC-TAGGGATCACGGTTCGAGTCCTGCCT
PVHY01003810.1:13426-13886(-)  : TTTGTTCATAGACCCCTTAGATTAAGAACACCTGCTCAAGGCTGTGCACGGTGGCTCGTGCTTGTAAGCTTCAGCCTATGAAGCGGCCGGGGC-TAGGGATCGCGGTTCGAGTCCCGCCT
PVHY01009730.1:53713-54174(+)  : AGAACATATATGAACACATACCAAAAAATATCTTAGGAAGGCCGGGCACGGTGGCTCGTGCTTGTAAGCTTCAGCCTATGAAGCGGCCGGGGC-TAGGGATCGCGGTTCGAGTCCCGCCT
PVHY01012311.1:1227-1691(-)    : GCCACCGCATAGGAAATTTACTATAAAGCTTTCAAATAAGGCTGGGCACGGTGGCTCGTGCTTGTAAGCTTCAGCCTATGAAGCGGCCGGGGC-TAGGGATCGCGGTTCGAGTCCCGCCT
PVHY01016875.1:11594-12058(+)  : CTAGCAATAACATTAAACACGTAATTATTAAAAGTCCTAGGCCGGGCACGGTGGCTCGTGCTTGTAAGGTTCAGCCTATGAAGCCGCCGGGGC-TAGGGATCGCGGTTCGAGTCCCACCT
PVHY01180168.1:0-381(-)        : ATAATGAAGTAAAATAATTTTAAAAGAAAAGAAAGACAAGGCCAGGCACGGTGGCTCGTGCTTGTAAGCTTCAGCCTATGAAGCGGCCGGGGC-TAGGGATCGCGGTTCGAGTCCCGCCT
PVHY01009930.1:15458-15918(-)  : GGGAAACAGTTTGGCAGTTCTTCAAAAAGTTAAACATAAGGCCGGGCACGGTGGCTCGTGCTTGTAAGCTTCAGCCTATGAAGCGGCCGGGGC-TAGGGATCGCGGTTCGAGTCCCGCCT
PVHY01045966.1:1383-1843(-)    : TAGTTTCCTTTTATTTTTTTAATAATAAAGAAAGATGAAGGCCGGGCACGGTGGCTCGTGCTTGTAAGCTTCAGCCTATGAAGCGGCCGGGGC-TAGGGATCGCGGTTCGAGTCCCACCT
PVHY01000878.1:41476-41936(-)  : GCTAGGCGCCCGCTATTTGTTACATTAAAAGACTATTTTAGCCGGGCACGGTGGCTCGTGCTTGTAAGCTTCAGCCTATGAAGCGGCCGGGGC-TAGGGATCGCGGTTCGAGTCCCGCCT
PVHY01009677.1:43648-44109(-)  : GGCAACATAGCGAGACCTAGTCTCAAAAAAAAGAAGAGGGGCCGGGCACGGTGGCTCGTGCTTGTAAGCTTCAGCCTATGAAGCGGCCGGGGC-TAGGGATCGCGGTTCGAGTCCCGCCT
PVHY01003949.1:35743-36203(-)  : GGACTGATTTATTCTATTGATTAAGAAACCAGTGCTTAAGGCCGGGCATGGTGGCTCGTGCTTGTAAGCTTCAGCCTATGAAGTGGCCGGGGC-TAGGGATCGCGGTTTGAGTCCCGCCT
PVHY01002269.1:76217-76677(+)  : AAAAGGACACCAGATTCAGATTTACAAAGGCTGTGCTAAGGCCGGGCACGGTGGCTCGTGCTTGTAAGCTTCAGCCTATGAAGCGGCCGGGGC-TAGGGATCGCGGTTCGAGTCCCGCCT
PVHY01094534.1:37-497(-)       : AATTAGCGAGGCATAGTCTTTTAGAAAAATAGTAATTTAGGCCGGGCACGGTGGCTCGTGCTTGTAAGCTTCAGCCTATGAAGCGGCCGGGGC-CAGGGATCGCGGTTCGAGTCCCGCCT
PVHY01000828.1:3392-3853(+)    : TTAGTATGATGCAGTATCAGCGGTGCAGAAAGGACTTTGGGCCGGGCACGGTGGCTCATGCTTGTAAGCTTCAGCCTATGAAGCGGCCGGGGCTTAGGGATCGCGGTTCGAGTCCCGCCT
PVHY01002549.1:79670-80132(-)  : TTCCCTTAAGTGGTAAAACTTTTTAAGAATCACCTTCAAGGCCGGGCACGGTGGCTCGTGCTTGTAAGCTTCAGCCTATGAAGCGGCCGGGGCTTAGGGATCGCGGTTCGAGTCCCGCCT
PVHY01004562.1:2711-3173(-)    : CCAAATTGAACATTATTGGTACCTTAGAAACTGCTTGTAGGCCGGGCACGGTGGCTCGTGCTTGTAAGCTTCAGCCTATGAAGCGGCCGGGGCTTAGGGATCACGGTTTGAGTCCCGCCT
PVHY01012448.1:22154-22616(+)  : AATCCCAAGAGAGATTACTCATCAAGACTGAGGGCCCAAGGCCGGGCACGGTGGCTCGTGCTTGTAAGCTTCAGCCTATGAAGCGGCCGGGGCTTAGGGATCACGGTTCAAGTCCCGCCT
PVHY01018318.1:7452-7914(-)    : ATGTCCTAATGTTCCTTACAATTCAAAATGGATCGTCAAGGCCAGGCACGGTGGCTCGTGCTTGTAAGCTTCAGCCTATGAAGCGGCCGGGGCTTAGGGATCGCGGTTCGAGTCCCACCT
PVHY01030613.1:7241-7942(+)    : CAAAAGCCTGCGGTTCCACAGCCAAAAATAGTAAGACCAGGCCGGGCACGGTGGCTCGTGCTTGTAAGCTTCAGCCTATGAAGCGGCCGGGGCTTAGGGATCGCGGTTCGAGTCCTGCCT
PVHY01006997.1:68467-68929(-)  : TGTAACAATAAACTATAGTTTTTTTAAAAAGTGAATCAAGGCCGGGCACGGTGGCTCGTGCTTGTAAGCTTCAGCCTATGAAGCGGCCGGGGCTTAAGGATCGCGGTTCGAGTCCCGCCT
PVHY01004607.1:35032-35493(+)  : TGACAAAACAGGATTTTAAAATGAAACATAAGTTCTGAAGGCCGGGCACGGTGGCTCGTGCTTGTAAGCTTCAGCCTATGAAGCGGCCGGGGCTTAGGGATCGCGGTTCGAGTCCCGCCT
PVHY01011287.1:22983-23444(+)  : TCTTTCAAAAGGTTTACCTGTTAAAAAGTAGCCAGTACTGGCCGGGCACGGTGGCTCGTGCTTGTAAGCTTCAGCCTATGAAGTGGCCGGGGCTTAGGGATCGCGGTTCGAGTCCCGCCT
PVHY01002465.1:6893-7355(-)    : CAAGTCATCTGGGGCTCTTGTTAAAAAGTAGTTGGTGAAGGCCGGGCACGGTGGCTCGTGCTTGTAAGCTTCAGCCTATGAAGCGGCTGGGGCTTAGGGATCGTGGTTCGAGTCCCGCCT
PVHY01002514.1:48322-48782(+)  : ATCCCAGAGCCCCTCACTGCAGTGTTTAAAGTAGCCATTAGCCGGGCACGGTGGCTCGTGCTTGTAAGCTTCAGCCTATGAAGCGGCCGGGGCTTAGGGATCGCGGTTCGAGTCCCGCCT
PVHY01009530.1:46209-46671(+)  : GGCTCTGAGACAGGTATTCTCCAAGATCTAACCCCAAAAGGCCGGGCACGGTGGCTCGTGCTTGTAAGCTTCAGCCTACGAAGCGGCCGGGGCTTAGGGATCGCGGTTCGAGTCCCGCCT
PVHY01004700.1:67022-67484(-)  : AAGGAGTATACAGGATGTTATTAAGATCTAACTTGATAAGGCCGGGCACGGTGGCTCGTGCTTGTAAGCTTCAGCCTATGAAGCGGCCGGGGCTTAGGGATCGCGGTTCGAGTCCCACCT
PVHY01001190.1:157652-158113(- : ACTGGGTAATTTCTTTCCTTCTATCAACCGGTAAATGTCGGCCGGGCACGGTGGCTCGTGCTTGTAAGCTTCAGCCTATGAAGCGGCCGGGGCTTAGGGATCGCGGTTCGAGTCCCGCCT
PVHY01000636.1:36465-36926(+)  : AACAAGTGTTAGTAAGAATGAAGAGAAAAAACAATGCTAGGCCGGGCACGGTGGCTAGTGCTTGTAAGCTTCAGCCTATGAAGCGGCCGGGGC-TAGGGATCGCAGTTCGAGTCCCGCCT
PVHY01005636.1:17251-17712(+)  : AAGAACAGGAAAGAAGACACAGTAAAGAGAAAGAGGGAAGGCCGGGCACGGTGGCTCGTGCTTGTAAGCTTCAGCCTATGAAGCGGCCAGGGC-TAGGGATCGCGGTTCGAGTCCCACCT
PVHY01018285.1:24866-25326(-)  : AGATGTATTAGAGATGATTTTAATTAAAATTCAAATTATGGCCGGGCACGGTGGCTCGTGCTTGTAAGCTTCAGCCTATGAAGCGGCCGGGGC-TAGGGATTGCGGTTCGAGTCCCGCCT
PVHY01027168.1:3480-3941(+)    : GGAAAGGGGTGGAAGAAAGGGCTAAAGAGGTGTAAGGAAGGCCGGGCACGGTGGCTCGTGCTTGTAAGCTTCAGCCTATGAAGCGGCCGGGGC-TAGGGATCGTGGTTCGAGTCCCGCCT
PVHY01054352.1:1347-1808(+)    : TTAGAGTATGTGAATTATAAATAAGAAAACAGTTGATCAGGCCGGGCACGGTGGCTCGTGCTTGTAAGCTTCAGCCTATGACGCGGCCGGGGC-TAGGAATCGCGGTTCGAGTCCCGCCT
PVHY01012201.1:12008-12468(+)  : AAAAAAGAAAAAAAAAGGTATCTTAAGAAAATATGAATAGGCCGGGCACGGTGGCTCGTGCTTGTAAGCTTCAGCCTATGAAGCAGCCGGGGC-TAGGGATCGCGGTTCGAGTCCCGCCT
PVHY01000554.1:111525-111985(- : TGTAGAGCAGGCCCCCTGGAAGCCCAAACAAGAATTTGAGGCCGGGCACGGTGGCTCGTGCTTGTAAGCTTCAGCCTATGAAGCGGCTGGGGC-TAGGGATCGCGGTTCGAGTCCCGCCT
PVHY01000922.1:123481-123941(- : GATTTAGAGAATGTAAATAGAGTAAGAAATTGGTAAACAGGCCGGGCACGGTGGCTCGTGCTTGTAAGCTTCAGCCTATGAAGCGGCCGGGGC-TAGGGATCGCGGTTCCAGTCCCGCCT
PVHY01004425.1:60288-60748(+)  : AATTATTTAAAAACATGCTGTATCAAAAAAATCAAAATAGGCCGGGCACGGTGGCTCGTGCTTGTAAGCTTCAGCCTATGAAGCGGCCGGGGC-TAGGGATCGCGGTTCGAGTCCCGCCT
PVHY01012836.1:2324-2784(+)    : AAAAAGGCAGGAAAATCTTCTACAAAGATGAGGGCATGAGGCCGGGCACGGTGGCTCGTGCTTGTAAGCTTCAGCCTATGAAGCGGCCGGGGC-TAGGGGTCGCGGTTGGAGTCCCGCCT
PVHY01067373.1:1777-2237(-)    : TACCCGGCTGAGCCAAGGTAGTGCAAGATACGGAAGTGAGGCCGGGCACAGTGGCTCATGCTTGTAAGCTTCAGCCTATGAAGCGGCTGGGGC-TAGGGATCGCGGTTCGAGTCCCGCCT
PVHY01075250.1:948-1408(+)     : GAATTAAACAAAAGAAACTAATTATTTTAAAAATTAAAAAGCCGGGCACGGTGGCTCGTGCTTGTAAGCTTCAGCCTATGAAGCGGCCGGGGC-TAGGGATCGCGGTTCGAGTCCCGCCT
PVHY01003790.1:38072-38532(+)  : AAAAAAATACAAATATGCTAACCATGAAAACTAGTGTAAGGCCGGGCATGGTGGCTCGTGCTTGTAAGCTTCAGCCTATGAAGCGGCCGGGGC-TAGGGATCGTGGTTCGAGTCCCGCCT
PVHY01012767.1:3364-3824(+)    : ATTGGTATGAAATTATACATTTGAAAAAGTACAAATGTAAGCCGGGCACGGTGGCTCGTGCTTGTAAGCTTCAGCCTATGAAGCGGCCGGGGC-TAGGGATCGCGGTTCGAGTCCCGCCT
PVHY01017727.1:1639-2100(-)    : GAAGAGGGAGCTGGTGGTGGCTTAGAAGCCATAGTGGAAGGCTGGGCACGGTGGCTCGTGCTTGTAAGCTTCAGCCTATGAAGCGGCCGGGGC-TAGGGATCGCGGTTCGAGTCCCGCCT
PVHY01007834.1:57293-57754(-)  : GGTTATGCAGAAAATAACTGATAAGAGTTATACTGTTCAGGCCGGGCACGGTGGCTCGTGCTTGTAAGCTTCAGCCTATGAAGCGGCCGGGGC-TAGGGATTGCGGTTCGAGTCCTGCCT
PVHY01042715.1:4832-5294(-)    : TTCTCAGGAAGCAGACTGAAGACTGAAAGAAGGCATGAAGGCCGGGCACGGTGGCTCGTGCTTGTAAGCTTCAGCCTATGAAGCGGCCGGGGC-TAGGGATCGTGGTTCGAGTCCCGCCT
PVHY01011504.1:12048-12511(+)  : AATATCACTTTGTAATTCAAAATAAAAATTAGATTGGAAGGCTGGGCACGGTGGCTCGTGCTTGTAAGCTTCAGCCTATGAAGCGGCCTGGGC-TAGGGATCACGGTTCGAGTCTTGCCT
PVHY01035808.1:878-1341(-)     : TTATCATATTTACACTAACATATAAGAAGGAAAATTTTCGGCCGGGCACGATGGCTCGTGCTTGTAAGCTTCAGCCTATGAAGCAGCCGGGGC-TAGGGATCGCGGTTCGAGTCCCGCCT
```

```
PVHY01016752.1:22261-22723(-)  : ATCAAGAGACATGGCATCCTTATTAAGGCCCTTGGACATGGCCGGGCACGGTGGCTCGTGCTTGTAAGCTTCAGCCTATGAAGCGGCCGGGGC-TAGGGATCGCGGTTCGAGTCCCGCCT
PVHY01072951.1:1387-1851(-)    : GGAAACAGTACGCAAGAGGGTGGGAATTGAAAATATTTGAGCCGGGCACGGTGGCTCGTGCTTGTAAGCTTCAGCCTATGAAGCGGCCGGGGC-TAGGGATCGCGGTTCGAGTCCCGCCT
PVHY01013237.1:20453-20915(+)  : TTTTGGGGACAGAATTGCCTTCACCAGAATATTCACAGCAGCCGGGCACGGTGGCTCGTGCTTGTAAGCTTCAGCCTATGAAGCGGCCGGGGC-TAGGGATCGCGGTTCGAGTCCCACCT
PVHY01005556.1:19245-19705(-)  : ATACAGAGCTTTAGATTGTTTTTGAAAGTAGTTTCCCAAGGCCGGGCATGGTGGCTCGTGCTTGTAAGCTTCAGCCTATGAAGCGGCCGGGGC-TAGGGATCGCGGTTCGAGTCCCGCCT
PVHY01008302.1:25016-25476(-)  : GGAAATTGGGCATGGAGGGGCACAAAAGACTAAGACAAGGCCGGGCACAGTGGCTCGTGCTTGTAAGCTTCAGCCTATGAAGCGGCCGGGGC-TAGGGATCGCGGTTCGAGTCCCGCCT
PVHY01020077.1:20797-21257(-)  : ATCCAATGTAACAAGTCTCATAAAAAGAGAAAAGACATAGACCGGGCACGGTGGCTCGTGCTTGTAAGCTTCAGCCTATGAAGCGGCCGGGGC-TAGGGATCGCGGTTCGAGTCCCGCCT
PVHY01016119.1:30218-30678(+)  : GTTCCTATATGAAGGCTTCATTTAAAATTTTTTTTTGAAGGCCGGGCACGGTGGCTCGTGCTTGTAAGCTTCAGCCTATGAAGCGGCCGGGGC-TAGGGATCGCGGTTCGAGTCCCGCCT
PVHY01072528.1:831-1291(+)     : TCTAGTCTTATGTTGTTTTCCACAAAGATCATTTTTGCAGGCCGGGCACGGTGGCTTGTGCTTGTAAGCTTCAGCCTATGAAGCGGCCGGGGC-TAGGGATCGCGGTTGGAGTCCGGCCT
PVHY01040643.1:4929-5390(-)    : TGCTGGAACACACATGTTTTGTTTAAAAATGAATAACAAGGCCGGGCACGGTGGCTCGTGCTTGTAAGCTTCAGCCTATGAAGCGGCCGGGGC-TAGTGATCGCGGTTCGAGTCCCGCCT
PVHY01001829.1:15787-16247(+)  : GCCATGGTGTTCATTTCTTTATTTTAAAACTAAGCCCCAGGCCGGGCACGGTGGCTCGTGCCTGTAAGCTTCAGCCTATGAAGCGGCCGGGGC-TAGGGATCGCGGTTCGAGTCCCGCCT
PVHY01004361.1:42264-42724(-)  : ATTATGTTTATCTAGGGACTTGATTAAAACTGAACTTGAGGCCGGGCACGGTGGCTCGTGCTTGTAAGCTTCAGCCTATGAAGCGGCCGGGGC-TAGGGATCGCGGTTCGAGTCCCGCCT
PVHY01023686.1:8656-9116(+)    : ATTTTATTGTGTAGCAATTTTGATTAAGAGAAACTGAAAGGCCGGGCACGGTGGCTCGTGCTTGTAAGCTTCAGCCTATGAAGCGGCCGGGGC-TAGGGATTGCGGTTCGAGTCCCACCT
PVHY01012641.1:23680-24146(-)  : TCAATTAACACAAATAACAGCATAGAAAGGAGACTGCCAGGCCGGGCACGGTGGCTCGTGCTTGTAAGCTTCAGCCTATGAAGCGGCCGGGGC-TAGGGATCGCGGTTCGAGTCCCGCCT
PVHY01015847.1:13399-13868(-)  : AGACGGAAACAAGATTTGCAGTTAAATGGTACTTTTGGAGGCCGGGCACGGTGGCTCGTGCTTGTAAGCTTCAGCCTATGAAGCGGCCGGGGC-TAGGGATTGCGGTTCGAGTCCCGCCT
PVHY01024847.1:6681-7142(+)    : CCTTCAAGAAGCAAGTGTTCCACGGTAACATGAATGTAAGGCCGGGCACGGTGGCTCGTGCTTGTAAGCTTCAGCCTATGAAGCGGCCGGGGC-TAGGGATCGCGGTTCGAGTCCCGCCT
PVHY01015071.1:21756-22218(-)  : ACTTTCCATGATAGTATTTCTTTAAAAACTAAAAGGTGAGGCCGGGCATGGTGGCTCGTGCTTGTAAGCTTCAGCCTATGAAGTAGCCGGGGC-TAGGGATCGCGGTTCGAGTCCCGCCT
PVHY01004144.1:90643-91107(-)  : CCATTAGTCTCGGCTCTTGAAAAACAAAACAATGGGTTAGGCCGGGCACGGTGGCTCGTGCTTGTAAGCTTCAGCCTATGAAGCGGCCGGGGC-TAGGGATCGCGGTTCAAGTCCCACCT
PVHY01023427.1:8943-9408(-)    : ACCAGTAAGAATGGCCCCTATCAAAAATATGGGCAACAAGGCCGGGCACGGTGGCTCGTGCTTGTAAACTTCAGCCTATGAAGCGGCTGGGGC-TAGGGATCGCGGTTCGAGTCCCGCCT
PVHY01001716.1:75005-75473(+)  : GTTCCTTTTTTATTGATTTCTTAAAAAATTTGAAGCCAAGGCCGGGCACGGTGGCTCGTGCTTGTAAGCTTCAGCCTATGAAGCGGCCGGGGC-TAGGGATCGCGGTTCGAGTCCCGCCT
PVHY01006475.1:44649-45110(-)  : CTGAATAAGAAAACACCTTTTTCAAAATATTATGTTCTAGGCCGGGCACGGTGGCTCGTGCTTGTAAGCTTCAGCCTATGAAGCGGCTGGGGC-TAGGGATCGCAGTTCGAGTCCCGCCT
PVHY01000775.1:104571-105031(- : AATTGCTGAAATTTTTCACTATTATAAGTCATACTGTGGGGCCAGGTACGGTGGCTCGTGCTTGTAAGCTTCAGCCTATGAAGCGGCCGGGGC-TAGGGATCGCGGTTCGAGTCCCGCCT
PVHY01057101.1:1698-2159(+)    : TTAAGGGGCTTATCCTTCTGGCTACTATAAACAATGATAGGCCGGGCACGGTGGCTCGTGCTTGTAAGCTTCAGCCTATGAAGCGGCCGGGGC-CAGGGATCACGGTTCGAGTCCCGCCT
PVHY01030803.1:11960-12420(-)  : AATAGAGGAAGAAACCAGGGCTTAAAAAAGACTCATCAAGGCCGGGCACGGTGGCTCGTGCTTGTAAGCGTCAGCCTATGAAGCGGCCGGGGC-TAGGGATCGCGGTTCGAGTCCCGCCT
PVHY01002408.1:33880-34341(-)  : GGGGAAGAGAGTAGAACACCAATAGAAATGTGTAACCAAGGCCGGGCACGGTGGCTCGTGCTTGTAAGCTTCAGCCTATGAAGCGGCCGGGGC-TAGGGATCGCGGTTCGAGTCCCGCCT
PVHY01015780.1:7059-7519(+)    : TCCACAGTGTTCAAAAGTCCTAAACAGAAGGCCATTTTAGGCCGGGCACGGTGGCTCGTGCTTGTAAGCTTCAGCCTATGAAGCGGCCGGGGC-TAGGGATCGCGGTTCGAGTCCCGCCT
PVHY01017180.1:14222-14682(+)  : TTAAGAACATTTGGAAAACAAAAATAAAAGTCATCCATAGGCCGGGCACGGTGGCTCGTGCTTGTAAGCTTCAGCCTATGAAGCAGCCGGGGC-TAGGGATCGCGGTTCGAGTCCCACCT
PVHY01004181.1:11239-11699(+)  : AGGCGCTGGCAGCCTTCACAGGCTAAATTGTGTTTGGTAGGCCGGGCACGGTGGCTCGTGCTTGTAAGCTTCAGCCTATGAAGCGGCCGGGGC-TAGGGATCGCGGTTCGAGTCCCGCCT
PVHY01016978.1:11379-11839(-)  : ACATTAATCCTGCATGCTCAGGATAGAATAGTCATGGTAGGCCGGGCACGGTGGCTCGTGCCTGTAAGCTTCAGCCTATGAAGCGGCCGGGGC-TAGGGATCGCGGTTCGAGTCCCGCCT
PVHY01005577.1:45995-46455(-)  : GAAAACATTAAAGTTACTTAGAATAAAATACAGAATCTAGGCCGGGCACGGTGGCTCGTGCCTGTAAGCTTCAGCCTATGAAGCGGCAGGGGC-TAGGGATCGCGGTTCGAGTCCCGCCT
PVHY01003103.1:12322-12782(-)  : GAAGAAGAAGAAGCATGAGATTAAATGACCTCAGAATGAGGCCGGGCACGGTGGCTCGTGCTTGTAAGCTTCAGCCTATGAAGCGGCCGGGGC-CAGGGATCGCGGTTCGAGTCCCGCCT
PVHY01005700.1:70829-71289(+)  : GGGAAGAAAAAAATAAGTGAGTAATATCAAGATATTTAAGGCCGTGCACGGTGGCTCGTGCTTGTAAGCTTCAGCCTATGAAGCGGCCGGGGC-TAGGGATCGCGGTTCGAGTCCCGCCT
PVHY01167867.1:29-489(+)       : TTGAGAGTAAAATTTACTAGCTAAAGAATAGGCTCTTAAGGCCGGGCACGGTGGCTCGTGCTTGTAAGCTTCAGCCTATGAAGCGGCCGGGGC-TAGGGATCGCGGTTCGAGTCCCGCCT
PVHY01013412.1:39263-39726(-)  : GACAAGAAAAAGTTAAAACTAAAAAACTGACAGAACTAAGGCCGGGCACGGTGGCTCGTGCTTGTAAGCTTCAGCCTATGAAGCGGCAGGGGC-TAGGGATCGCGGTTCGAGTCCCGCCT
PVHY01036498.1:2252-2712(-)    : CACCTAACCACCATGGTTTCTCATCAGAAAGCTACATCAGGCCGGGCACGGTGGCTCGTGCTTGTAAGCTTCAGCCTATGAAGCGGCCGGGGC-TAGGGATCGCGGTTCGAGTCCCGCCT
PVHY01004447.1:65916-66376(+)  : GTAAAATGATAGGTTATGCCTTAATAAATCTCCTAATAAGGCCGGGCACGGTGGCTCGTGCTTGTAAGCTTCAGCCTATGAAGCGGCCGGGGC-TAGGGATCGCGGTTCGAGTCCCGCCT
PVHY01032970.1:4990-5450(+)    : AAAGGCAGCTGATGTGAGTATTAAAGATGGGACATGGAAGGCCGGGCACGGTGGCTCATGCTTGTAAGCTTCAGCCTATGAAGCGGCCGGGGC-CTGGGATCGCGGTTCGAGTCCCGCCT
PVHY01024371.1:4748-5208(-)    : CTAAAACAATAACTATATAATATCTATATAATATTAGAAGGCCGGGCACGGTGGCTCGTGCTTGTAAACTTCAGCCTATGAAGCGGCCGGGGC-TAGGGATCGCGGTTCGAGTCCCGCCT
PVHY01064486.1:913-1373(+)     : TATGTTCATAAAGTGTGTTTCAAAAGTTCTCTTATACAAGGCCGGGCACGGTGGCTCGTGCTTGTAAGCTTCAGCCTATGAAGCGGCCGGGGC-TAGGGATCGCGGTTCGAGTCCCGCCT
PVHY01018181.1:27724-28184(-)  : TCCTGTCCCGAACATAATCTTCAAATAGTGATGTGCAAAGGCCGGGCACGGTGGCTCGTGCTTGTAAGCTTCAGCCTATGAAGCGGCCGGGGC-TAGGGATTGCGGTGCGAGTCCCGCCT
PVHY01037172.1:5469-5929(-)    : CTTTAATAGTAATATAATATTCTAAAATTATTATACTTAGGCCGGGCACGGTGGCTCGTGCTTGTAAGCTTCAGCCTATGAAGCGGCTGGGGC-TAGGGATCGCGGTTCAAGTCCCGCCT
PVHY01243375.1:0-398(+)        : ACATGCTTGTAGAAAAGTATGTTAAACATACAATTGTGAGGCCGGGCACGGTGGCTCGTGCTTGTACGCTTCAGTCTATGAAGCGGCCGGGGC-TAGGGATCGCGGTTCGAGTCCCGCCT
PVHY01012394.1:11145-11605(+)  : CAATAAAATTTCTAGTTGCCTTAAAAATAAATATTCCTAGGCCGGGCACGGTGGCTCGTGCTTGTAAGCTTCAGCCTATGAAGCGGCCGGGGC-TAGGGATCGCGGTTCGAGTCCCGCCT
PVHY01022271.1:16325-16786(-)  : AGCATATCACTTGATATCATTAGAACTTTGCCAGCCTGAGGCCGGGCACGGTGGCTCGTGCTTGTAAGCTTCAGCCTATGAAGCGACCGGGTC-TAGGGATCACGGTTCGAGTCCCGCCT
PVHY01024968.1:6195-6655(+)    : TTTAGTACTTGAAAACTTGGTTTAACTTTGAAACTGCAAGGCCGGGCACGGTGGCTCGTGCTTGTAAGCTTCAGCCTATGAAGCGGCCGGGGC-TAGGGATCGCGGTTCGAGCCCCGCCT
PVHY01004013.1:46042-46502(-)  : ATTCAATTTACTACTTCCCCTTTAGACTTACCAAAAGAAGGCCGGGCACGGTGGCTCGTGCTTGTAAGCTTCAGCCTATGAAGCGGCCGGGGC-TAGGGATCGCGGTTCGAGTCCCACCT
```

```
                                             *       140         *       160         *       180         *       200         *       220         *       240
Urop_c tribe 1                 : GGGGCCCGTAGCGAGACCTAGTTCC--------------AAAAAAATAGCTGGGCGTGGTGGCGCGCGCCTGTCATCCCAGCTACCTGGGAAGCTGAGGCTGGGGATTGCTTGAGTCTGT
PVHY01002401.1:48683-49143(+)  : GGGGCCCGTAGCGAGACCTAGTTCC--------------AAAAAAATAGCTGGGCGTGGTGGCGCGCGCCTGTCATCCCAGCTACCTGGGAAGCTGAGGCTGGGGATTGCTTGAGTCTGT
PVHY01007456.1:45401-45875(+)  : GGGGCCCGTAGCGAGACCTAGTTCCAAAAAAAAAAAAAAAAAAAAATAGCTGGGCGTGGTGGCGCGCGCCTGTCATCCCAGCTACCTGGGAAGCTGAGGCTGGGGATGGCTTGAGTCTTT
PVHY01006771.1:32287-32748(+)  : GGGGCACGTAGCGAGACCTAGTTCC--------------AAAAAAATAGCTGGGCGTGGTGGCGCGCGCCTGTCATCCCAGCTACCTGGGAAGCTGAGGCTGGGGATTGCTTGAGTCTGT
PVHY01045252.1:4788-5249(-)    : GGGGCCCGTAGCGAGACCTAGTTCC--------------AAAAAAATAGCTGGGCGTGGTGGCGCACGCCTGTCATCCCAGCTACCTGGGAAGCTGAGGCTGGGGATTGCTTGAGTCTGT
PVHY01007539.1:38627-39087(+)  : GGGGCCCGTAGCGAGACCTAGTTCC--------------AAAAAAATAGCTGGGCGTGGTGGTGCGCGCCTGTCATCCCAGCTACCTGGGAAGCTGAGGCTGGGGATGGCTTGAGTCTGT
PVHY01037078.1:1316-1776(-)    : GGGGCCCGTAGCGAGACCTAGTTCC--------------AAAAAAATAGTTGGGCGTGGTGGCGCGCGCCTGTAATCCCAGCTACCTGGGAAGCTGAGGCTGGGGATTGCTTGAGTCTGT
PVHY01013704.1:32430-32891(-)  : GGGGCCCGTAGCGAGACCTAGTTCC--------------AAAAAAAAGCTGGGCGTGGTGGTGCGCGCCTGTCATCCCAGCTACCTGGGAAGCTGAGGCTGGGGATTGCTTGAGTCTGT
PVHY01000998.1:68954-69416(-)  : GGGGCCCGTAGCGAGACCTAGTTCC-------------AAAAAATACAGCTGGGCGTGGTGGCGCGCGCCTGTCATCCCAGATACCTGGGAAGCTGAGGCTGGGGATTGCTTGAGTCCGT
```

```
PVHY01015967.1:4675-5136(-)      : GGGGCCCGTAGCGAGACCTAGTTCC-------------AAAAAAAATAGCTGGGCGTGGTGGTGCACGCCTGTCATCCCAGCTACCTGGGAAGCTGAGGCTGGGGATTGCTTGAGTCTGT
PVHY01008490.1:9516-9977(-)      : GGGGCCCGTAGCGAGACCTAGTTCC-------------AAAAAATACAGCTGGGCGTGGTGGCGCGCGCCTGTCATCCCAGCTACCTGGGAAGCTGAGGCTGGGGATTGCTTGAGTCCGT
PVHY01003810.1:13426-13886(-)    : GGGGCCCGTAGCGAGACCTAGTTCC--------------AAAAAAATAGCTGGGCGTGGTGGCGCGCGCCTGCCATCCCAGCTACTTGGGAAGCTGAGGCTGGGGATTGCTTGAGTCTGT
PVHY01009730.1:53713-54174(+)    : GGGGCCCGTAGCGAGACCTATTTCC-------------AAAAAAAATAGCTGGGCGTGGTGGCGCACGCCTGTCATCCCAGCTACCTGGGAAGCTGAGGGTGGGGATGGCTTGAGTCTGT
PVHY01012311.1:1227-1691(-)      : GGGGCCCGTAGCGAGACCTAGTTCC--------------AAAAAAATAGCTGGGCGTGGTGGCATGCGCCTGTCATCCCAGCTACTTGGGAAGCTGAGGCTGGGGATTGCTTGAGTCTGT
PVHY01016875.1:11594-12058(+)    : GGGGCCCATAGCGAGACCTAGTTCC--------------AAAAAAATAGCTGGGCGTGGTGGCGCGCGCCTGTCATCCCAGCTACCTGGGAAGCTGAGGCTGGGAATTGCTTGAGTCTGT
PVHY01180168.1:0-381(-)          : GGGGCCCGTAGTGAGACCTAGTCCC--------------AAAAAAATAGCTGGGCGTGGTGGCGCGCGCCTGTCATCCCAGCTACCTGGGAAGCTGAGGCTGGGGATTGCTTGAGTCTGT
PVHY01009930.1:15458-15918(-)    : GGGGCCCGTAGCGAGACCTAGTTCC--------------AAAAAAATAGCTGGGCGTGGTGGCGCGCGCCTGTAATCCCAGCTAACTGGGAAGCTGAGGCTGGGGATTGCTTGAGTCTGT
PVHY01045966.1:1383-1843(-)      : GGGGCCCGTAGTGAGACCTAGTTCC--------------AAAAAAATAGCTGGGCGTGGTGGCGCACGCCTGTCATCCCAGCTACCTGGGAAGCTGAGGCTGGGGATTGCTTGAGTCTGT
PVHY01000878.1:41476-41936(-)    : GGGGCCCGTAGCGAGACCTAGTTCC------------AAAAAAAAATAGCTGGGCGTGGTGGCGCGCGCCTGTCATCCCGGCTACCTGGGAAGCTGAGGCTGGGGATTGCTTGAGTCTGT
PVHY01009677.1:43648-44109(-)    : GGGGCCCGTAGCGAGACCTAGTTCC------------AAAAAAAAATAGCTGGGCTTGGTGGTGCGCGCCTGTCATCCCAGCTACCTGGGAAGCTGAGGCTGGGGATTGCTTGAGTCTGT
PVHY01003949.1:35743-36203(-)    : GGGGCCCGTAGCGAGACCTAGTTCC--------------AAAAAAATAGCTGGGCGTGGTGGCGCGTGCCTGTCATCCCAGCTACCTGGGAAGCTGAGGCTGGGGATTGCTTGAGTCTGT
PVHY01002269.1:76217-76677(+)    : GGGGCCCGTAGCGAGACCTAGTTCC--------------AAAAAAATAGCTGGGCGTGGTGGCGCACGCCTGTCATCCCAGCTACCTGGGAAGCTGAGGCTGGGGATGGCTTGAGTCTGT
PVHY01094534.1:37-497(-)         : GGGGCCCATAGCGAGACCTAGTCCC--------------AAAAAAATAGCTGGGCGTGGTGGCACGCGCCTGTCATCCCAGCTACCTGGGAAGCTGAGGCTGGGGATTGCTTGAGTCTGT
PVHY01000828.1:3392-3853(+)      : GGGGCCCGTAGCGAGACCTAGTTCC-------------AAAAAAAATAGCTGGGCGTGGTGGTGCACGCCTGTCATCCCAGCTACCTGGGAAGCTGAGGCTGGGGATTGCTTGAGTCTGT
PVHY01002549.1:79670-80132(-)    : GGGGCCCATAGCGAGACCTAGTTCC-------------AAAAAAAATAGCTGGGCATGGTGGTGCACGCCTGTCATCCCAGCTACCTGGGAAGCTGAGGCTGGGGATTGCTTGAGTCTGT
PVHY01004562.1:2711-3173(-)      : GGGGCCCGTAGCGAGACCTAGTTCC-------------AAAAAAAATAGCTGGGCGTGGTGGTGCACGCCTGTCATCCCAGCTACCTGGGAAGCTGAGGCTGGGGATTGCTTGAGTCTGT
PVHY01012448.1:22154-22616(+)    : GGGGCCCGTAGCGAGACCTAGTTCC-------------AAAAAAAATAGCTGGGCGTGGTGGTGCACGCCTGTCATCCCAGCTACCTGGGAAGCTGAGGCTGGGGATTGCTTGAGTCTGT
PVHY01018318.1:7452-7914(-)      : GGGGCCCGTAGCGAGACCTAGTTCC-------------AAAAAAAATAGCTGGGCGTGGTGGTGCACGCCTGTCATCCCAGCTACCTGGGAAGCTGAGGCTGGGGATTGCTTGAGTCTGT
PVHY01030613.1:7241-7942(+)      : GGGGCCCGTAGCGAGACCTAGTTCC-------------AAAGAAAATAGCTGGGCGTGGTGGTGCACGCCTGTCATCCCAGCTACCTGGGAAGCTGAGGCTGGGGATTGCTTGAGTCTGT
PVHY01006997.1:68467-68929(-)    : GGGGCCCGTAGCGAGACCTAGTTCC-------------AAAAAAAATAGCTGGGCGTGGTGGTGCACGCCTGTCATCCCAGCTACCTGGGAAGCTGAGGCTGGGGATTGCTTGAGTCTGT
PVHY01004607.1:35032-35493(+)    : GGGGCCCGTAGCGAGACCTAGTTCC-------------AAAAAAAATAGCTGGGCGTGGTGGTGCGCGCCTGTCATCCCAGCTACCTGGGAAGCTGAGGCTGGGGATTGCTTGAGTCTGT
PVHY01011287.1:22983-23444(+)    : GGGGCCCGTAGCGAGACCTAGTTCC-------------AAAAAAAATAGCTGGGCGTGGTGGTGCACGCCTGTCATCCCAGATACCTGGGAAGCTGAGGCTGGGGATTGCTTGAGTCTGT
PVHY01002465.1:6893-7355(-)      : GGGGCCCGTAGCGAGACCTAGTTCC-------------AAAAAAAATAGCTGGGCGTGGTGGTGCACGCCTGTCATCCCAGCTACCTGGGAAGCTGAGGCTGGGGATTGCTTGAGTCTGT
PVHY01002514.1:48322-48782(+)    : GGGGCCCGTAGCGAGACCTAGTTCC-------------AAAAAAAATAGCTGGGCGTGGTGGTGCACGCCTGTCATCCCAGCTACCTGGGAAGCTGAGGCTGGAGATTGCTTGAGTCTGT
PVHY01009530.1:46209-46671(+)    : GGGGCCTGTAGCGAGACCTAGTTCC-------------AAAAAAAATAGCTGGGCGTGGTGGTGCACGCCTGTCATCCCAGCTACCTGGGAAGCTGAGGCTGGGGATTGCTTGAGTCTGT
PVHY01004700.1:67022-67484(-)    : GGGGCCCGTAGCGAGACCTAGTTCC-------------AAAAAAAATAGCTGGGCGTGGTGGTGCGCGCCTGTCATCCCAGCTACCTGGGAAGCTGAGGCTGGGGATTGCTTGAGTCTGT
PVHY01001190.1:157652-158113(-  : GGGGCCCGTAGCGAGACCTAGTTCC-------------AAAAAAAATAGCTGGGCGTGGTGGTGCACGCCTGTCATCCCAGCTACCTGGGAAGCTGAGGCTGGGGATTGCTTGAGTCTGT
PVHY01000636.1:36465-36926(+)    : GGGGCCTGTAGTGAGACCTAGTTCC-------------AAAAAAAATAGCTGGGCGTGGTGGCGCGCGCCTGTCATCCCAGCTACCTGGGAAGCTGAGACTGGGGATTGCTTGAGTCTGT
PVHY01005636.1:17251-17712(+)    : GGGGCCCGTAGCGAGACCTAGTTCC-------------AAAAAAAATAGCTGGGCGTGGTGGTGCGCGCCTGTAATCCCAGCTACCTGGGAAGCTGAGGCTGGGGATTGCTTGAGTCTGT
PVHY01018285.1:24866-25326(-)    : GGGGCCCGTAGCGAGACCTAGTTCC-------------AAAAAAAATAGCTGGGCGTGGTGGCGCGCGCCTGTCATCCCAGCTACCTGGGAAGCTGAGGCTGGGGATTGCTTTAGTCTGT
PVHY01027168.1:3480-3941(+)      : GGGGCCCATAGCGAGACCTAGTTCC-------------AAAAAAAATAGCTGGGCGTGGTGGTGCGCGCCTGTCATCCCAGCTACCTGGGAAGCTGAGGCTGGGGATTGCTTGAGTCTGT
PVHY01054352.1:1347-1808(+)      : GGGGCCCGTAGCGAGACCTAGTTGC-------------AAAAAAAATATCTGGGTGTGGTGGCGCGCGCCTGTCATCCCAGCTACCTGGGAAGCTGAGGCTGGGGATTGCTTGAGTCTGT
PVHY01012201.1:12008-12468(+)    : GGGGCCCATAGCGAGACCTAGTTCC-------------AAAAAAAATAACTGGGCGTGGTGGCGCGCGCCTGTAATCCCAGCTACCTGGGAGGCTGAGGCTGGGGATTGCTTGAGTCTGT
PVHY01000554.1:111525-111985(-  : GGGGCCCGTAGCGAGACCTAGTTCC--------------AAAAAAATAGCTGGGCGTGGTGGTGCGCGCCTGTCATCCCAGCTACCTGGGAAGCTGGGGCTGGGGATTGCTTGAGTCTGT
PVHY01000922.1:123481-123941(-  : GGGGCCCGTAGCGAGACCTAGTTCC--------------AAAAAAATAGCTGGGCGTGGTGGCGTGCGCCTGTCATCCCAGCTACCTGGGAAGCTGAGGCTGGGGATTGCTTGAGTTTGT
PVHY01004425.1:60288-60748(+)    : GGGGCTCGTAGCGAGACCTAGTTCC--------------AAAAAACTAGCTGGGTGTGGTGGCGCGCGCCTGTCATCCCAGCTACCTGGGAAGCTGAGGCTGGGGATTGCTTGAGTCTGT
PVHY01012836.1:2324-2784(+)      : GGGGCCTGTAGCGAGACCTAGTTCC--------------AAAAAAATAGCTGGGCGTGGTGGCGCGCGCCTGTCATCCCAGCTACCTGGGAAGCTGAGGCTGGGGATTGCTTGAGTCTGT
PVHY01067373.1:1777-2237(-)      : GGGGCCCGTAGCGAGACCTAGTTCC--------------AAAAAAATAGCTGGGCGTGGTGGCGCGCGCCTGTCATCCCAGCTACCTGGGAAGCTGAGGCTGGGGATTGCTTGAGTCTGT
PVHY01075250.1:948-1408(+)       : GGGGCCCGTAGTGACACCTAGTTCC--------------AAAAAATTAGCTGGGCGTGGTGGCGCGTGCCTGTCATCCCAGCTACCTGGGAAGCTGAGGCTGGGGATTGCTTGAGTCTGT
PVHY01003790.1:38072-38532(+)    : GGAGCCCGTAGCGAGACCTAGTTCC--------------AAAAAAATAGCTGGGCGTGGTGGCGCGTGCCTGTCATCCCAGCTACCTGGGAAGCTGAGGCTGGGGATTGCTTGAGTCTGT
PVHY01012767.1:3364-3824(+)      : GGGGCCGGTAGCGAGACCTAGTTCC--------------AAAAAAATAGCTGGGCGTGGTGGCGCGCGCCTGTCATCCCAGCTACCTGGGAAGCTGAGGCTGGGGATTGCTTGAGTCTGT
PVHY01017727.1:1639-2100(-)      : GGGGCCCGTAGCGAGACCTAGTTCC-------------AAAAAAAATAGCTGGGCATGGTGGTGCGCGCCTGTCATCCCAGCTACCTGGGAAGCTGAGGCTGGGGATTGCTTGAGTCTGT
PVHY01007834.1:57293-57754(-)    : GGGGCCCGTAGCGAGACCTAGTTCC-------------AAAAAAAATAGCTGGGCGTGGTGGTGCACGCCTGTCATCCCAGCTACCTGGGAAGCTGAGGCTGGGGATGGCTTGAGTCTGT
PVHY01042715.1:4832-5294(-)      : GGGGCCTGTAGCGAGACCTAGTTCC-------------AAAAAATACAGCTGGGCGTGGTGGCGCGCGCCTGTCATCCCAGCTACCTGGGAAGCTGAGGCTGAGGATTGCTTGAGTCCGT
PVHY01011504.1:12048-12511(+)    : GGGGCCCGTAGCGAGACCTAGTTCC--------------AAAAAAATAGCTGGGCGTGGTGGCGCGCGCCTGTCATCCCAGCTACCTGGGAAGCTGAGGCTGGGGATTGCTTGAGTCTGT
PVHY01035808.1:878-1341(-)       : GGGGCCTGTAGCGAGACCTAGTTCC--------------AAAAAAATAGCTGGGCGTGGTGGCGCGCGCCTGTCATCCCAGCTACCTGGGAAGCTGAGGCTGGGGATTGCTTGAGTCTGT
PVHY01016752.1:22261-22723(-)    : GGGGCCCGTAGCGAGACCTAGTTCC--------------AAAAAAATAGCTGGGCGTGGTGGCGTGCGCCTGTCATCCCAGCTACCTGGGAAGCTGAGGCTGGGGATTGCTTGAGTTTGT
PVHY01072951.1:1387-1851(-)      : GGGGCCCGTAGTGAGACCTAGTTCC--------------AAAAAAATAACTGGGCGTGGTGGCGTGCGCCTGTCATCCCAGCTACCTGGGAAGCTGAGGCTGGGGATTGCTTGAGTCTGT
PVHY01013237.1:20453-20915(+)    : GGGGCCCGTAGCGAGACCTAGTTCC--------------AAAAAAATAGCTGGGCGTGGTGGCACGCGCCTGTCATCCCAGCTACCTGGGAAGCTGAGGCTGGGGATTGCTTGAGTTTGT
PVHY01005556.1:19245-19705(-)    : GGGGCCCGTAGCGAGACCTAGTTCC--------------AAAAAAATAGCTGGACGTGGTGGCGTGCGCCTGTCATCCCAGCTACCTGGGAAGCTGAGGCTGGGGATTGCTTGAGTCTGT
PVHY01008302.1:25016-25476(-)    : GGGGCCCATAGCAAGACCTAGTTCC--------------AAAAAAATAGCTGGGCGTGGTGGCGTGCGCCTGTCATCCCAGCTACCTGGGAAGCTGAGGCTGGGGATTGCTTGAGTCTGT
PVHY01020077.1:20797-21257(-)    : GGGGCCCGTAGCAAGACCTAGTTCC--------------AAAAAAATAGCTGGGCGTGGTGTCGCGCGCCTGTCATCCCAGCTACCTGGGAAGCTGAGGCTGGGGATTGCTTGAGTCTGT
PVHY01016119.1:30218-30678(+)    : GGGGCCCATAGCGAGACCTAGTTCC--------------AAAAAAATAGCTGGGCGTGGTGGTGCGCGCCTGTCATCCCAGCTACCTGGGAAGCTGAGGCTGGGGATTGCTTGAGTCTGT
PVHY01072528.1:831-1291(+)       : GGGGCCCGTAGCGAGACCTAGTTCC--------------AAAAAAATAGCTGGGTGTGGTGGCGCGCGCCTGTCATCCCAGCTACCTGGGAAGCTGAGGCTGGGGATTGCTTGAGTCTGT
PVHY01040643.1:4929-5390(-)      : GGGGCGCGTAGCGAGACCTAGTTCC-------------AAAAAAAATAGCTGGGCGTGGTGGTGCGCGCCTGTAATCCCAGCTACCTGGGAAGCTGAGGCTGGGGATTGCTTGAGTCTGT
PVHY01001829.1:15787-16247(+)    : GGGGCCCGTAGCGAGACCTAGTTCC--------------AAAAAAATAGCTGGGCGTGGTGGCGCGTGCCTGTAATCCCAGCTACCTGGGAAGCTGAGGCTGGGGATTGCTTGAGTCTGT
```

```
PVHY01004361.1:42264-42724(-)  : GGGGCCCGTAGCGAGACCTAGTTCC--------------AAAAAAATAGCTGGGCGTGGTGGTGCGCGCCTGTCATCCCAGCTACCTGGGAAGCTGAGGCTGGGGATTGCTTGAGTCTGT
PVHY01023686.1:8656-9116(+)    : GGGGCCCATAGCGAGACCTAGTTCC--------------AAAAAAATAGCTGGGCGTGGTGGCGCGCGCCTGTCATCCCAGCTACCTGGGAAGCTGAGGCTGGGGATGGCTTGAGTCTGT
PVHY01012641.1:23680-24146(-)  : GGGGCCCGTAGCGAGACCTAGTTCC-------AAAAAAAAAAAAAAATAGCTGGGCGTGGTGGTGCACGCCTGTCATCCCAGCTACCTGGGAAGCTGAGGTTGGGGATGGCTTGAGTCTGT
PVHY01015847.1:13399-13868(-)  : GGGGCCTGTAGCGAGACCTAGTTCC-----AAAAAAAAAAAAAAAAATAGCTGGGCGTGGTGGCGCGCGCCTGTCATCCCAGCTACCTGGGAAGCTGAGGTTGGGGATTGCTTGAGTCTGT
PVHY01024847.1:6681-7142(+)    : GGGGCCCGTAGCGAGACCTAGTTCC------------AAAAAAAAATAGCTGGGCGTGGTGGTGCGCGCCTGTCATCCCAGCTACCTGGGAAGCTGAGGCTGGGGATTGCTTGAGTCTGT
PVHY01015071.1:21756-22218(-)  : GGGGCCCGTAGCGAGACCTAGTTCC-----------AAAAAAAAAATAGCTGGGCGTGGTGGTGCGCGCCTGTCATCCCAGCTACCTGGGAAGCTGAGGCTGGGGATTGCTTGAGTCTGT
PVHY01004144.1:90643-91107(-)  : GGGGCCCGTAGCGAGACCTAGTTCC----------AAAAAAAAAAATAGCTGGGCGTGGTGGTGTGCGCCTGTCATCCCAGCTACCTGGGAAGCTGAGGCTGGGGATTGCTTGAGTCTGT
PVHY01023427.1:8943-9408(-)    : GGGGCCCGTAGCGAGACCTAGTTCC---------AAAAAAAAAAAATAGCTGGGCGTGGTGGTGCGCGCCTGTCATCCCAGCTACCTGGGAAGCTGAGGCTGGGGATTGCTTGAGTCTGT
PVHY01001716.1:75005-75473(+)  : GGGGCCCGTAGCGAGACCTAGTTCC------AAAAAAAAAAAAAAATAGCTGGGCGTGGTGGCGCGCGCCTGTCATCCCAGCTACCTGGGAAGCTGAGGCTGGGGATGGCTTGAGTCTGT
PVHY01006475.1:44649-45110(-)  : GGGGCCCGTAGTGAGACCTAGTTCC-------------AAAAAAAATAGCTGGGCGTGGTGGCGCGCGCCTGTCATCCCAGCTACCTGGGAAGCTGAGGCTGGGGATTGCTTGAGTCTGT
PVHY01000775.1:104571-105031(- : GGGGCCCGTAGCGAGACCTAGTTCC-------------AAAAAAAATAGCTGGGCGTGGTGACGCGTGCCTGTCATCCCAGCTACCTGGGAAGCTGAGGCTGGGGATTGCTTGAGTCTGT
PVHY01057101.1:1698-2159(+)    : GGGGCCCGTAGCGAGACCTAGTCCC-------------AAAAAAAATAGCTGGGCGTGGTGGCGCGCGCCTGTCATCCCAGCTACCTGGGAAGCTGAGGCTGGGGATGGCTTGAGTCTGT
PVHY01030803.1:11960-12420(-)  : GGGGCCCGTAGCGAGACCTAGTTCC-------------AAAAAAAATAGCTGGGCGTGGTGGCGCGCACCTGTCATCCCAGCTACCTGGGAAGCTGAGGCTGGGGATTGCTTGAGTCTGT
PVHY01002408.1:33880-34341(-)  : GGGGTCCGTAGGGAGACCTAGTTCC-------------AAAAAAAATAGCTGGGCGTGGTGGCGCGCACCTGTCATCCCAGCTACCTGGGAAGCTGAGGCTGGGGATTGCTTGAGTCTGT
PVHY01015780.1:7059-7519(+)    : GGGGCCCGTAGCGAGAACTAGTTCC--------------AAAAAAATAGCTGGGCGTGGTAGTGCGCGCCTGTCATCCCAGCTACCTGGGAAGCTGAGGCTGGGGATTGCTTGAGTCTGT
PVHY01017180.1:14222-14682(+)  : GGGGCCCGTAGCGAGACCTAGTTCC--------------AAAAAAATAGCTGGGCGTGGTGGCGCGCGCCTGTCATCCCAGCTACCTGGGAAGCTGAGGCTGGGGATTGCTTGAGTCTGT
PVHY01004181.1:11239-11699(+)  : GGGGCCCGTAGCGAGACCTAGTTCC--------------AAAAAAATAGCTGGGCGTGGTGGCACGCGCCTGTCATCCCAGCTACCTGGGAAGCTGAGACTGGGGATTGCTTGAGTCTGT
PVHY01016978.1:11379-11839(-)  : GGGGCCCGTAGTGAGACCTAGTTCC--------------AAAAAAATAGCTGGGCGTGGTGGCACGCGCCTGTCATCCCAGCTAACTGGGAAGCTGAGGCTGGGGATTGCTTGAGTCTGT
PVHY01005577.1:45995-46455(-)  : GGGGCCCGTAGCGAGACCTAGTTCC--------------AAAAAAATAGCTGGGCGTGGTGGCGCGCGCCTGTCATCCCAGCTACCTGGGAAGCTGAGGCTGGGGATTGCTTGAGTCTGT
PVHY01003103.1:12322-12782(-)  : GGGGCCCGTAGCGAGACCTAGTCCC--------------AAAAAAATAGCTGGGCGTGGTGGCGCGCGCCTGTAATCCCAGCTACCTGGGAAGCTGAGGCTGGGGATTGCTTGAGTCTGT
PVHY01005700.1:70829-71289(+)  : GGGGCCCGTAGCGAGACCTAGTTCC--------------AAAAAAATAGCTGGGCGTGGTGGCGCGCGCTTGTCATCCCAGCTACCTGGGAAGCTGAGGCTGGGGATTGCTTGAGTCTGT
PVHY01167867.1:29-489(+)       : GGGGCCCGTAGCGAGACCTAGTTCC--------------AAAAAAATAGCTGGGTGTGGTGGCGCGCGCCTGTAATCCCAGCTACCTGGGAAGCTGAGGCTGAGGATTGCTTGAGTCTGT
PVHY01013412.1:39263-39726(-)  : GGGGCCTGTAGCGAGACCTAGTTCC--------------AAAAAAATAGCTGGGCGTGGTGGCGTGCGCCTGTCATCCCAGCTACCTGGGAAGCTGAGGCTGGGGATTGCTTGAGTTTGT
PVHY01036498.1:2252-2712(-)    : GGGGCCCGTAGCGAGACCTAGTTAC--------------AAAAAAATAGCTGGGCGTGGTGGCCCGCGCCTGTCATCCCAGCTACCTGGGAAGCTGAGGCTGGGGATGGCTTGAGTCTGT
PVHY01004447.1:65916-66376(+)  : GGGGCCCGTAGCGAGACCTAGTTCC--------------AAAAAAATAGCTGGGCATGGTGGCGCGCGCCTGTCATCCCAGCTACCTGGGAAGCTGAGGCTGGGGATTGCTTGAGTCTGT
PVHY01032970.1:4990-5450(+)    : GGGGCCCGTAGTGAGACCTAGTACC--------------AAAAAAATAGCTGGGCGTGGTGGCGCGCGCCTGTCATCCCAGCTACCTGGGAAGCTGAGGCTGGGGATTGCTTGAGTCTGT
PVHY01024371.1:4748-5208(-)    : GGGGCCCGTAGCGAGACCTAGTTCC--------------AAAAAAATAGCTGGGCGTGGTGGCGCGCGCCTGTCATCCCAGCTACCTGGGAAGGTGAGGCTGGGGATTGCTTGAGTCTGT
PVHY01064486.1:913-1373(+)     : GGGGCCCGTAGCGAGACCTAGTTCC--------------AAAAAAATAGCTGGGCGTGGTGGCGCACGCCTGTCATCCCAGCTACCTGGGAAGCTGAGGCTGGGGATTGCTTGAGTCTGT
PVHY01018181.1:27724-28184(-)  : GGGGCCCGTAGCGAGACCTAGTTCC--------------AAAAAAATAGCTGGGCGTGGTGGCGTGCGCCTGTCATCCCAGCTACCTGGGAAGCTGAGGCTGGGGATTGCTTGAGTCTGT
PVHY01037172.1:5469-5929(-)    : GGGGCCCGTAGCGAGACCTAGTTCC--------------AAAAAAATAGCTGGGCGTGGTGGCGTGCGCCTGTCATCCCAGCTACCTGGGAAGCTGAGGCTGGGGATTGCTTGAGTCTGT
PVHY01243375.1:0-398(+)       : GGGGCCCGTAGCGAGACCTAGTTCC--------------AAAAAAATAGCTGGGCGTGGTGGCGCGCGCCTGTCATCCCAGCTACCTGGGAAGCTGAGGCTGGGGATTGCTTGAGTCTGT
PVHY01012394.1:11145-11605(+)  : GGGGCCCGTAGCGAGACCTAGTTCC--------------AAAAAAATAGCTGGACGTGGTGGCGCGCGCCTGTCATCCCAGCTACCTGGGAAGCTGAGGCTGGGGATTGCTTGAGTCTGT
PVHY01022271.1:16325-16786(-)  : GGGGCCCGTAGCGAGACCTAGTTCC-------------AAAAAAAATAGCTGGGCGTGGTGGCACGCGCCTGTCATCCCAGCTACCTGGGAAGCTGAGGCTGGGGATTGCTTGAGTCTGT
PVHY01024968.1:6195-6655(+)    : GGGGCCCGTAGCGAGACCTAGTTCC-------------AAAAAAAATAGCTGGGCGTGGTGGCACGCACCTGTCATCCCAGCTACCTGGGAAGCTGAAGCTGGGGATTGCTTGAGTCTGT
PVHY01004013.1:46042-46502(-)  : GGGGCCCATAGCGAGACCTAGTTCC--------------AAAAAAATAGCTGGGCGTGGTGGCGCGCGCCTGTCATCCCAGCTACCTGGGAAGCTGAGGCTGGGGATTGCTTGAGTCTGT

                                         *      260          *      280          *      300          *      320          *      340          *      360
Urop_c tribe 1                  : GGTACCCCGGGCAACGAGAGCCTGCGGTTCCACAGGC------AAAAATAGCAAGACCTAGTCTCAAAAAAAAAAAAAA
PVHY01002401.1:48683-49143(+)  : GGTACCCCGGGCAACGAGAGCCTGCGGTTCCACAGGC------AAAAATATCAAGACCTAGTCTCAAAAAAAAAAAAAAAAAAAAAAAAAAAAAAAAAAAAAAAAAAAAAAAAAAAAAAAA
PVHY01007456.1:45401-45875(+)  : GGTACCCCGGGCAACGAGAGCCTGCGGTTCCACGGGC------AAAATTAGCAAGACCTAGTCTCAAAAAAAAAAAAAAAAAAAAAAAAAAAAAAAAAAAAAAAAAAAAAAAAAAAAAAAA
PVHY01006771.1:32287-32748(+)  : GGTACCCCAGGCAACAAGAGCCTGCGGTTCCACAGGC------AAAAATAGCAAGACCTAGTCTCAAAAAAAAAAAAAAAAAAAAAAAAAAAAAAAAAAAAAAAAAAAAAAAAAAAAAAAA
PVHY01045252.1:4788-5249(-)    : GGTACCCCGGGCAACGAGAACCTGCGGTTCCACAGGC------AAAAATAGCAAGACGTAATCTCAAAAAAAAAAAAAAAAAAAAAAAAAAAAAAAAAAAAAAAAAAAAAAAAAAAAAAAA
PVHY01007539.1:38627-39087(+)  : GGTACCCCGGGCAACGAGAGCCTGCGGTTCCACGGGC------AAAAATAGCGAGACCTAGTCTCAAAAAAAAAAAAAAAAAAAAAAAAAAAAAAAAAAAAAAAAAAAAAAAAAAAAAAAA
PVHY01037078.1:1316-1776(-)    : GGTACCCCGGGCAACGAGAGCCTGCGGTTCCACAGGC------AAAAATAGCAAGACCTAGTCTCAAAAAAAAAAAAAAAAAAAAAAAAAAAAAAAAAAAAAAAAAAAAAAAAAAAAAAAA
PVHY01013704.1:32430-32891(-)  : GGTACCCCGGGCAACGAGAGCCTGCGGTTCCACAGAC------AAAAATAGCAAGACCTAATCTCAAAAAAAAAAAAAAAAAAAAAAAAAAAAAAAAAAAAAAAAAAAAAAAAAAAAAAAG
PVHY01000998.1:68954-69416(-)  : GGTACCCCGGGCAACGAGAGCCTGCGGTTCCACAGGC-----AAAAAATAGCAAGACCTAGTCTCAAAAAAAAAAAAAAAAAAAAAAAAAAAAAAAAAAAAAAAAAAAAAAAAAAAAACA
PVHY01015967.1:4675-5136(-)    : GGTACCCCGGGCAAAGAGAGCCTGCGGTTCCACAGGC------AAAAATAGCAAGACCTAGTCTCAAAAAAAAAAAAAAAAAAAAAAAAAAAAAAAAAAAAAAAAAAAAAAAAAAAATGA
PVHY01008490.1:9516-9977(-)    : GGTACCCCGGGCAACGAGAGCCTGCGGTTCCACAGGC------AAAAATAGCAAGACCTAGTCTCAAAAAAAAAAAAAAAAAAAAAAAAAAAAAAAAAAAAAAAAAAAAAAAAAGAAAT
PVHY01003810.1:13426-13886(-)  : GGTACCCCGGGCAACGAGAGCCTGCGGTTCCACAGGC------AAAAATAGCGAGACCTAGTCTCAAAAAAAAAAAAAAAAAAAAAAAAAAAAAAAAAAAAAAAAAAAAAAGAACACCT
PVHY01009730.1:53713-54174(+)  : GGTACCCCGGGCAACGAGAGCCTGCGGTTCCACAGGC------AAAAATAGCAAGACCTAATCTCAAAAAAAAAAAAAAAAAAAAAAAAAAAAAAAAAAAAAAAAAAAAATCTTAGGAAC
PVHY01012311.1:1227-1691(-)    : GGTACCCCGGGCAACGAGAGCCTGCGGTTCCACAGGC--AAAAAAAAATAGCAAGACCTAGTCTCAAAAAAAAAAAAAAAAAAAAAAAAAAAAACAAAAAAAAAAAAAAAAAAAAAAAAA
PVHY01016875.1:11594-12058(+)  : GGTACCCCGGGCAACGAGAGCCTGCGGTTCCACAGGC--AAAAAAAAATAGCAAGACCTAGTCTCAAAAAAAAAAAAAAAAAAAAAAAAAAAAAAAAATAAAAAAAAAAAAAAAAAAAAA
PVHY01180168.1:0-381(-)        : GGTACCCCGGGCAACGAGAGCCTACGGTTCCACGGGC------AAAAATAGCAAGACCTAGTCTCAAAAAAAAAAAAAAAGAAAAGAAAAGAAAAGAAAGACAACTCCTCATTAAAAAAA
PVHY01009930.1:15458-15918(-)  : GGTACCCCGGGCAATGAGAGCCTGCGGTTCCACAGGC------AAAAATAGCAAGACCTAGTCTCAAAAAAAAAAAAAAAAAGTTAAACATAGATTTACTATATGACCAGAAATTTCACT
PVHY01045966.1:1383-1843(-)    : GGTACCCCGGGCAATGAGAGCCTGCGGTTCCACAGGC------AAAAATAGCAAGACCTAGTCTCAAAAAAAAAAAAAAAAAAAGATGAGAATTGCCTGATGACTAGAGTGCAGGTTGTC
PVHY01000878.1:41476-41936(-)  : GGTACCCCGGGCAAAGAGAGCCTGCGGTTCCACAGGC------AAAAATAGCAAGACCTAGTCTCCAAAAAAAAAAAAAAGACTATTTTAGGTAGCCCTTAGGTTTTTTAGGGGCCATT
```

```
PVHY01009677.1:43648-44109(-)   : GGTACCCCGGGCAACGAGAGCCTGCGGTTCCACGGGC------AAAAATAGCAAGACCTAGTCTCAAAAAAAAAAAGAAGAGGGTCTAGCAGTTCCTCAAAATGTGCATTCAGAGTTACT
PVHY01003949.1:35743-36203(-)   : GGTACCCTGGGCAACGAGAGCCTGCGGTTCCACAGGC------AAAAATAGCAAGACCTAGTCTCAAAAAAAAAAAAAAGAAAAGAAAAGAAAAAAAAAAGAAACCAGTGCTTAGAGAGT
PVHY01002269.1:76217-76677(+)   : GGTACCCCGGGCAACGAGAGCCTGCGGTTCCACAGGC------AAAAATAGCAAGACCTAATCTCAAAAAAAAAAAAAAAATAATAAAAAATAAAAAAAAAATACAAAGGCTGTGCTAAAT
PVHY01094534.1:37-497(-)        : GGTACCCCGGGCAACGAGAGCCTGCGGTTCCACAGGC------AAAAATAGCAAGACCTAGTCTCAAAAAAAAAAAAAATTAAAAAAATTAAAAAAATTAAAAAATAGTAATTTGATTAT
PVHY01000828.1:3392-3853(+)     : GGTACCCCGGGCAACGAGAGCCTGCAGTTCCACAGGC------AAAAATAGCAAGACCTAGTCTCAAAAAAAAAAAAAAAAAAAACAAAACAAAAAAAAAAACAAAACAAAAAACTTTAA
PVHY01002549.1:79670-80132(-)   : GGTACCCAGGCAACGAGAGCCTGCGGTTCCACAGGC------AAAAATAGCAAGACCTAGTCTCAAAAAAAAAAAAAAAAAAAAAAGAATCACCTTCAATAGCAGGGGCTGACTTAAA
PVHY01004562.1:2711-3173(-)     : GGTACCCCGGGCAACGAGAGCCTGCGGTTCCACAGGC------AAAAATAGCAAGACCTAGTCTCAAAAAAAAAAAAAAAGAAACTGCTTGTAATCCTCTGTCCTCAATCATGTTTCCTG
PVHY01012448.1:22154-22616(+)   : GGTACCCCGGGCAACGAGAGCCTGCGGTTCCACAGGC------AAAAATAGCAAGACCTAGTCTCAAAAAAAAAAAAAAAAAAAAAAAGACTGAGGGCCCATTGGAGGTAATGAACCAAA
PVHY01018318.1:7452-7914(-)     : GGTACCCCGGGCAACGAGAGCCTGCGGTTCCACAGGC------AAAAATAGCAAGACCTAGTCTCAAAAAAAAAAAAAAGAAAAAAGAAAAAAAAAACAAAATGGATCGTCAAAAAATTA
PVHY01030613.1:7241-7942(+)     : GGTACCCCGGGCAACGAGAGCCTGCGGTTCCACAGGC------AAAAATAGCAAGACCTAGTCTCAAAAAAAAAAAAAAAAATATTAAGACCTAGTCTCAAATACATAAATAAATAAATA
PVHY01006997.1:68467-68929(-)   : GGTACCCCGGGCAACAAGAGCCTGCGGTTCCACAGGC------AAAAATAGCAAGACCTAGTCTCAAAAAAAAAAAATAAATAAATAAAAAAATAAAAAATAAAAAATAAAAAGTGAATC
PVHY01004607.1:35032-35493(+)   : GGTACTCCGGGCAACGAGAGCCTGCGGTTCCACAGACC------AAAAATAGCAAGACCTAATCTCAAAAAAAAAAAAAGAAAAAAAAAAAAAAAAGAAACATAAGTTCTGACTTGGACTAG
PVHY01011287.1:22983-23444(+)   : GGTACCCCGGGCAACGAGAGCCTGCGGTTCCACAGGC------AAAAATAGCAAGACCTAGTCTCAAAAAAAAAATAAATAAATAAATAAAAATAAAAAAAAAATAAAAAGTAGCCAGTA
PVHY01002465.1:6893-7355(-)     : GGTACCCCGGGCAACGAGAGCCTGCGGTTCCACAGGC------AAAAATAGCAAGACCTAATCTCAAAAAAAAAAAAAAAAAAAAAAAAAGTAGTTGGTGAGAGCTGGGCATGGTGGCAC
PVHY01002514.1:48322-48782(+)  : AGTACCCCGGGCAACGAGAGCCTGCGGTTCCACAGGC------AAAAATAGCAAGACCTAGTCTCAAAAAAAAAAAAAAAATAAATAAATAAAAAAATAAAGTAGCCATTAATGCACTTG
PVHY01009530.1:46209-46671(+)   : GGTACTCCGGGCAACGAGAGCCTGCGGTTCCACAGGC------AAAAATAGCAAGACCTAGTCTCAAAAAAAAAAAAAAAAAAAAAAAAGATCTAACCCCAAGAGCTCACATCTCCCCCT
PVHY01004700.1:67022-67484(-)   : GGTACCCCGGGCAACGAGAGCCTGCGGTTCCACAGAC------AAAAATAGCAAGACCTAATCTCAAAAAAAAAAAAAAAAAAGATCTAACTTGATAGACTTTTCATATTACTGCAAATG
PVHY01001190.1:157652-158113(- : GGTACCCCGGACAACGAGAGCCTGCGGTTCCACAGGC------AAAAATAGCAAGACCTAGTCTAAAAAAAAAAAAAAAAAAAAAAAAAATCAACCGGTAAATGTCAATTTGGAAGAGCAGA
PVHY01000636.1:36465-36926(+)   : GGTACCCCGGGCAACGAGAGCCTGCGGTTCCACAGGC------AAAAATAGCAAGACCTAGTCTCAAAAAAAAAAAAAAAGAAAAAAAAAACAATGCTCATACAATGCTGTTAATGCTG
PVHY01005636.1:17251-17712(+)   : GGTACCCCGGGCAACGAGAGCCTGCGGTTCCACGGGC------AAAAATAGCAAGACCTAGTCTCAAAAAAAAAAAAAAAAAAAAAAAAGAGAAAGAGGGAAGCGCGACTTAGAAGCCAGA
PVHY01018285.1:24866-25326(-)   : GGTACCCCGGGCAACGAGAGCCTGCGGTTCCACAGGC------AAAAATAGCAAGACCCAGTCTCAAAAAAAAAAAAAAAAAAAAAATTCAAATTATGTATCATAAACAAGAGTTATCTATG
PVHY01027168.1:3480-3941(+)     : GGTACCCTGGGCAACGAGAGCCTGCGGTTCCACAGGC------AAAAATAGCCAGACCTAGTCTCAAAAAAAAAAAAAAAAAAAAAAAAAAAAAAAAAGAGGTGTAAGGGACACTGTCGTG
PVHY01054352.1:1347-1808(+)     : GGTACCCCGGGCAACGAGAGCCTGCGGTTCCACAGGC------AAAAATAGCAAGACCTAGTCTCAAAAAAAAAAAAAAAAAAAAGAAAACAGTAGATCGACTTCACACATAATATAGGTA
PVHY01012201.1:12008-12468(+)   : GGTACCCCGGGCAACGAGAGCCTGCGGTTCCACAGGC------AAAAATAGCAAGACCTAGTCTCAAAAAAAAAAAAAGAAAATATGAATATAGCTCACAATTAAAGTTAGCATCTAAGC
PVHY01000554.1:111525-111985(- : GGTACCCCGGGCAAAGAGAGCCTGCGGTTCCACAGGC------AAAAATAGCAAGAGCTAGTCTCAAAAAAAAAAAAAAAAAAAAAAAAAAAAAAGAATTTGCTTTATCAATCAGTGCGATG
PVHY01000922.1:123481-123941(- : GGTACCCCGGGCAACGAGAGCCTGCGGTTCCACAGGC------AAAAATAGCAAGACCTAGTCTCAAAAAAAAAAAAAAAGAAATTGGTAAGCATGTCTAAAAACCTCCAAAAAGATAGT
PVHY01004425.1:60288-60748(+)   : GGTACCCCGGGCAACGAGAGCCTGCGGTTCCACAGGC------AAAAATAGCAAGACCTAGTCTCAAAAAAAAAGAAAAAAAAATTAAAAAAAAATCAAAATAAAACATGTTAAATCTTT
PVHY01012836.1:2324-2784(+)     : GGTACCCAGGCAACGAGAGCCTGCGGTTCCACAGGC------AAAAATAGCAAGACCTAGTCTCAAAAAAAAAAAAAAGATGAGGGCATGAAGAAATCTGTTTAGTGGCAATGGGTTGT
PVHY01067373.1:1777-2237(-)     : GGTACCCCGGGCAACGAGAGCCTGCGGTTCCACGGGC------AAAAATAGCAAGACCTACTCTCAAAAAAAAAAAAAAAAAAAGATACGAAAGTGATTTTTATTCTGAGCTGACCCGTTA
PVHY01075250.1:948-1408(+)      : GGTACCCCGGGCAACGAGAGCCTGCGGTTCCACAGGC------AAAAATAGCAAGACCTAGTCTCAAAAAAAAAAAAAAAATTAAAAAAGGCAAATGTGTTTACTTGAATGAACAAATGA
PVHY01003790.1:38072-38532(+)   : GGTACCCCGGGCAACGAGAGCCTGTGGTTCCACAGGC------AAAAATAGCAAGACCTAGTCTCAAAAAAAAAAAAAAAAAAAAAAAGAAAACTAGTGTAATTTCCTAAACTATAGGTC
PVHY01012767.1:3364-3824(+)     : GGTACCCCGGGCAACGAGAGCCTGCGGTTCCACAGGC------AAAAATAGCAAGACCTACTCTCAAAAAAAAAAAAAAAAGAAAAAGTACAAATGTATTGTTTAAAATATTTGTGCATT
PVHY01017727.1:1639-2100(-)     : GGTACCCCGGGCAACGAGAGCCTGCGGTTCCACAGAC------AAAAATAGCAAGACCTAATCTCAAAAAAAAAAAAAAAAAAAAAAAAGAAGCCATAGTGGGATAGTGACTGGCAGAGA
PVHY01007834.1:57293-57754(-)   : GGTACCCCGGGCAACGAGAGCCTGCGGTTCCACAGGC------AAAAATAGCAAGACCTAGTCTCAAAAAAAAAAAAAAAAAAAAAAAAAAAAAGAGAGTTATACTGTTCAGTAAGTTATAC
PVHY01042715.1:4832-5294(-)     : GGTACCCCGGGCAACGAGAGCCTGCGGTTCCACAGGC-----AAAAAATAGCAAGACCTAGTCTCAAAAAAAAAAAAAAAAAAAAAAAAAAAAAAGAAGGCATGAAATGCTACTTTCAAAA
PVHY01011504.1:12048-12511(+)   : GGTACCCCGGGCAACGAGAGCCTGCGGTTCCACAGGC--AAAAAAAAATAGCAAGACCTAGTCTCAAAAAAAAAAAAAATTAGATTGGAGAACATCTGATCAAGTAGAAAATGAATTCTGA
PVHY01035808.1:878-1341(-)      : GGTACCCCGGGCAACAAGAGCCTGTGGTTCCACAGGC--AAAAAAAAATAGCAAGACCTAGTCTCAAAAAAAAAAAAAAAAGGAAAATTTTCTTTCCCGCCATGTCGTTCGTGGACAGCG
PVHY01016752.1:22261-22723(-)   : GGTACCCCGGGCAACGAGAGCCTGTGGTTCCACAGGC---AAAAAAAATAGCAAGACCTAGTCTCAAAAAAAAAAAAAAAAAAAAAAAAAAAAAATTAAGGCCCTTGGACAAAGAAAATACAG
PVHY01072951.1:1387-1851(-)     : GGTACCCCGGGCAACAAGAGCCTGCGGTTCCACAGGCAAAAAAAAAAATAGCAAGACCTAGTCTCAAAAAAAAAAAAAAGAAAATATTTGAATGTCTGGAATAATTACCCTGAATAAGCA
PVHY01013237.1:20453-20915(+)   : GGTACCCCGGGCAACGAGAACCTGCGGTTCCACAGGC--AAAAAAAAATAGCAAGACCTAGTCTCAAAAAAAAAAAAAAAAAAATTCACAGCATCCCAAACTTGCTTCACTGAGTTGCTC
PVHY01005556.1:19245-19705(-)   : GGTACCCCGGGCAACAAGAGCCTGTGGTTCCACAGGC------AAAAATAGCAAGACCTAGTCTCAAAAAAAAAAAAAAAAAAAAAAAAAAAGTAGTTTCCCTAAGGAGATATTTAAGCCA
PVHY01008302.1:25016-25476(-)   : GGTACCCAGGCAACGAGAGCCTGCGGTTCCACAGGC------AAAAATAGCAAGACCTAGTCTCAAAAAAAAAAAAAAAAAAAAAAAAGACTAAGACAATGGTTCATGTGTTATAAGGTGAT
PVHY01020077.1:20797-21257(-)   : GGTACCCCGGGCAACGAGAGCCTGTGGTTCCACAGGC------AAAAATAGCAAGACCTAGTCTCAAAAAAAAAAAAAAAAAGAAAAGACATGAAGACAGGTACAGAGAGAAAAGGGGGTC
PVHY01016119.1:30218-30678(+)   : GGTACCCTGGGCAACGAGAGCCTGCGGTTCCACAGGC------AAAAATAGCCAGACCTAGTCTCAAAAAAAAAAAAAAAAAAAAGTAATTGAACTAAATTTTTTTTTGTTTTTTGTTTGTT
PVHY01072528.1:831-1291(+)      : GGTACCCCGGGCAACGAGAGCCTGCGGTTCCACAGGC------AAAAATAGCAAGACCTAGTCTCAAAAAAAAAAAAAAAAAAAAAAAAAAGATCATTTTTGCCTCCATTTCTTTCAGTA
PVHY01040643.1:4929-5390(-)     : GGTACCCCGGGCAACGAGAGCCTGTGGTTCCACAGGC------AAAAATAGCAAGACCTAGTCTCAAAAAAAAAAAAAAAAAAAAAAATGAATAACAGCTAAGGCATCAACAAATTGTTC
PVHY01001829.1:15787-16247(+)   : GGTACCCCGGGCAACGAGAGCCTGCGGTTCCACGGGC------AAAAATAGCGAGACCTAGTCTCAAAAAAAAAAAAAAAGAATAAAACTAAGCCCCAAGTGCTACTGAATTTTGGCACT
PVHY01004361.1:42264-42724(-)   : GGTACCCCGGGCAAAGAGAGCCTGCGGTTCCACAGGC------AAAAATAGCAAGACCTAGTCTCAAACAAAAAAAAAAACAACAAAAAAAAACTGAAGTTGGTAAAAAGTCTTGGAAAA
PVHY01023686.1:8656-9116(+)     : GGTACCCCGGGCAACGAGAGCCTGCGGTTCCACAGGC------AAAAAAAGCAAGACCTAGTCTCAAAAAAAAAAAAAAAAAAAGAGAGAGAAACTGAAAGTAGCCAAAAACAATTGTACT
PVHY01012641.1:23680-24146(-)   : GGTACCCCGGGCAACGAGAGCCTGCAGTTCCACGGGC------AAAAATAGCAAGACCTAGTCTCAAAAAAAAAAAAAGAAGAAGAAAGGAGACTGCCAGCTTTGTGGTCAGAATCTCAA
PVHY01015847.1:13399-13868(-)   : GGTACCCCGGGCAACGAGAGCCTGCGGTTCCACGGGC------AAAAATAGCAAGACCTAGTCTCAAAAAAAAAAAAAAAAAAATGGTACTTTTGCTCCCCTCAGCGTCCCACTTACGTGG
PVHY01024847.1:6681-7142(+)     : GGTACCCCGGGCAACGAGAGCCTGTGGTTCCACGGGC------AAAAATAGTAAGGCCTAGTCTCAAAAAAAAAAAAAACAACAAAAAAACAAAAAACATGAATGTACTACAAGAAATCATA
PVHY01015071.1:21756-22218(-)   : GGTACCCCGGGCAACGAGAGCCTGCGGTTCCACAGGC------AAAAATAGCAAGACCTAGTCTCAAAAAAAAAAAAACTAAAGGTGGCCGGGCACGGTGGCTCGTGCCTATACACAGC
PVHY01004144.1:90643-91107(-)   : GGTACCCCGGGCAACGAGAGCCTGCGGTTCCACGGGC------AAAAATAGCAAGACCTAGTCTCAAAAAAAAAAAAAAAAACAACAACAAAACAATGGGTTGATAGCACAAGGTCTGTTC
PVHY01023427.1:8943-9408(-)     : GGTACCCCGGGCAACGAGAGCCTGCGGTTCCACAGGC------AAAAATAGCAAGACCTAGTCTCAAAAAAAAAAAAAAAAAAAAATGGGCAACAAGTGCTGGCCTGGATGTGGAGAGAAAG
PVHY01001716.1:75005-75473(+)   : GGTACCCTGGGCAACGAGAGCCTGCGGTTCCATGGGC------AAAAATAGCAAGACCTAGTCTCAAAAAAAAAAAAAAAAAAAAATTGAAGCCAGCATTGGTTTTATTTTAATAAGGAAA
PVHY01006475.1:44649-45110(-)   : GGTACCCCGGGCAACGAGAGCCTGCGGTTCCACGGGC------AAAAATAGCAAGACCTAGTCTCAAAAAAAAAAAAAAGAAAAACCAAAATATTATGTTCTAGCAACTTTTGGACTTTT
```

```
PVHY01000775.1:104571-105031(- : GGTACCCCGGGCAACGAGAGCCTGCGGTTCCACAGGC------AAAAATAGCAAGACCTAGTCTCAAAAAAAAAAAAAAAAAAAGTCATACTGTGATAAGCAATGTGAATTTGTTGCTAT
PVHY01057101.1:1698-2159(+)    : GGTACCCCGGGTAACGAGAGCCTGCGGTTCCACAGGC------AAAAATAGCAAGACCTAGTCTCAAAAAAAAAAAAAAAAAAAACAATGATGAGGACGCTTGTGTACAGTTTTTGGAGC
PVHY01030803.1:11960-12420(-)  : GGTACTCCGGGCAACGAGAGCCTGCGGTTCCACAGGC------AAAAATAGCAAGACCTAGTCTCAAAAAAAAAAAAAAAAAAGACTCATCACTTATCCACGATCACTTAGCCTTTCAAT
PVHY01002408.1:33880-34341(-)  : GGTACCCCGGGCAATGAGAGCCTGCGGTTCCACAGGC------AAAAATAGCGAGACCTAGTCTCAAAAAAAAAAAAAAAAAGAAATGTGTATATTAGATTAAGCATGATACAATAGTAA
PVHY01015780.1:7059-7519(+)    : GGTACCCTGGGCAAAGAGAGCCTGCGGTTCCACAGGC------AAAAATAGCAAGACCTAGTCTCAAAAAAAAAAAAAAAAAGACGGCCATTCATCCAAGAAGGCATTGTCCCATCAACA
PVHY01017180.1:14222-14682(+)  : GGTACCCCGGGCAACGAGAGTCTGTGGTTCCACAGGC------AAAAATAGCAAGACCTAGTCTCAAAAAAAAAAAAAAAAAAAGTCATCCATAAAGGTGAAGGTGATCATTGGAGCTAA
PVHY01004181.1:11239-11699(+)  : GGTACCCCGGGCAACGAGAGCCTGCGGTTCCACGGGC------AAAAATAGCGAGACCTAGTCTCAAAAAAAAAAAAAAAAAATTGTGTTTGGTGAGACACTTTATTTTAAATCACCTGA
PVHY01016978.1:11379-11839(-)  : GGTACCCCGGGCAACGAGAGCCTGCGGTTCCACAGGC------AAAAATAGCGAGACCTAGTCTCAAAAAAAAAAAAAAGAAAAAAAATAGTCATGGTGTTCTGATGCTGATAGATGCTG
PVHY01005577.1:45995-46455(-)  : GGTACCCCGGGCAACGAGAGCCTGCGGTTCCACGGGC------AAAAATAGCGAGACCTGGTCTCAAAAAAAAAAAAAAATACAGAATCTGCCTAATTTTCCAATTTCACCTCTTTTCAG
PVHY01003103.1:12322-12782(-)  : GGTACCCCGGGCAACGAGAGCCTGCGGTTCCACAGGC------AAAAATAGCAAGACCTAGTCTCAAAAAAAAAAAAAAAGACCTCAGAATGAAATGTTATATTCAGTAACTAAAACCTA
PVHY01005700.1:70829-71289(+)  : GGTACCCCGGGCAACGAGAGCCTGCGGTTCCATGGGC------AAAAAGAGCAAGACCTAGTCTCAAAAAAAAAAAAAAAAATCAAGATATTTAAAATATTTTTTAAAAAAAGAAAGTAA
PVHY01167867.1:29-489(+)       : GGTACCCCGGGCAACAAGAGCCTGCGGTTCCACGGGC------AAAAATAGCAAGACCTAGTCTCAAAAAAAAAAAAAACTTAAAGGACTTTAATTCCCAATTTGAGAAGCAAAATTAAT
PVHY01013412.1:39263-39726(-)  : GGTACCCCGGGCAACGAGAGCCTGCGGTTCCACAGGC---AAAAAAAATAGCAAGACCTAGTCTCAAAAAAAAAAAAAAAACTGACAGAACTAAAACAAAGAGAGGGGGATGGCCGGGTA
PVHY01036498.1:2252-2712(-)    : GGTACCCCGGGCAACGAGAGCCTGCGGTTCCACGGGC------AAAAATAGCAAGACTTAGTCTCAAAAAAAAAAAAAAAAAAAAAGCTACATCAAAATCATATTCCTTCACTTGAGGAA
PVHY01004447.1:65916-66376(+)  : GGTACCCCGGGCAACGAGAGCCTGCGGTTCCACAGGC------AAAAATATCAAGACCTAGTCTCAAAAAAAAAATAAATAAATAAATCTCCTAATAGAGATTTCTCTGATGGCCTTGTC
PVHY01032970.1:4990-5450(+)    : GGTACCCCGGGCAACGAGAGCCTGCGGTTCCACAGGC------AAAAATAGCAAGACCTAGTCTCAAAAAAAAAAAAAAAAAAAGATGGGGCATGGGATGGGCTTAATTTACATATAAAG
PVHY01024371.1:4748-5208(-)    : GGTACCCCGAGCAACAAGAGCCTGCGGTTCCATAGGC------AAAAATAGCAAGACCTAGTCTCAAAAAAAAAAAAAAACACTATATAATATTAAGAACAGCAGCTATACTTATTCATC
PVHY01064486.1:913-1373(+)     : GGTACCCAGGCAACGAGAGCCTGTGGTTTCACGGGC------AAAAATAGCAAGACCTAGTCTCAAAAAAAAAAAAAAAAGTTCTCTTATACAAAGTCACTGGAATCATGTCAGGACAC
PVHY01018181.1:27724-28184(-)  : GGTACCCCGGGCAACGAGAGCCTGCGGTTCCACAGGC------AAAAATAGCAAGACCTAGTCTCAAAAAAAAAATATATATATAGTGATGTGCAAAGTGTTGGCTTTTAAAATAAT
PVHY01037172.1:5469-5929(-)    : GGTACCCTGGGCAGCGAGAGCCTGCGGTTCCACAGGC------AAAAATAGCAAGACCTAGTCTCAAAAAAAAAAAAAAACCAAAATTATTATGCTTTCCCCATTGATGCCATTCAGGCT
PVHY01243375.1:0-398(+)       : GGTACCCCGGGCAACAAGAGCCTGCGGTTCCACAGGC------AAAAATAGCAAGACCTAGTCTCAAAAAAAAACCAAAAACATACAATTGTGAAGAAAACGCACGTGCTCTCAGCTCTG
PVHY01012394.1:11145-11605(+)  : GGTACCCCGGGCAACAAGAGCCTGCAGTTCCACAGGC------AAAAATAGCGAGACCTAGCCTCAAAAAAAAAAAAAATCCCTCTTCATGCATCCTGACTGTCTCCAGGAATAGTACTT
PVHY01022271.1:16325-16786(-)  : GGTACCCCGGACAACGAGAGCCTGCGGTTCCACAGGC------AAAAATAGCAAGACCTAGTCTCAAAAAAAAAAAAAAAAAAAACCAACTTTGCCAGCCTGATTGGTACAAAATGTCTA
PVHY01024968.1:6195-6655(+)    : GGTACCCCGGGCAACGAGAGCCTGCGGTTCCACGGGC------AAAAATAGCAAGACCTAGTCTCAAAAAAAAAAAAACAAACTTTGAAACTGCAAAGCTTGTAAGTCTTCTGGAGACTG
PVHY01004013.1:46042-46502(-)  : GGTACCCCGGGCAACGAGAGCCTGCAGTTCCACAGGC------AAAAATAGCAAGACTTAGTCTCAAAAAAAAAAAAAAATAGACTTATCAAAAGAAATAAAAATATACACACATTCTATA
```

```
                                         *        380         *        400         *        420

PVHY01002401.1:48683-49143(+)  : AAAAAAAAAAAAAAAAAAAAAAAAAAAAAAAAAAAAAAAAAAAAGTAATAAAACTGCTATG
PVHY01007456.1:45401-45875(+)  : AAAAAAAAAAAAAAAAAGAATATCTAGAACATTCTATCCTAATCATAATATATCTATGAAA
PVHY01006771.1:32287-32748(+)  : AAAAAAAAAAAAAAAGGTCAAATAAAAAGAATTCAATTCAAGGCCGCTGCACTTCATTTCC
PVHY01045252.1:4788-5249(-)    : AAAAAAAAAACAACTCATCCCTAACCTGCTCTAATAACTACCAGACTTTCCATTCAGGCAG
PVHY01007539.1:38627-39087(+)  : AAAAAAAATCAATTTAGAATTCCAAAAACAAATGCTGCTCCAAGGAGTCCCTTTGCCCCTG
PVHY01037078.1:1316-1776(-)    : AATCAGAACACTGAGATTCTTCGTAGCAACATTGTTAGAAATACAGTGAAACAAGGCTTTC
PVHY01013704.1:32430-32891(-)  : AACTAGTTAATTTTTAAATAGTCTTTAGGCAGGTGTGAAGCCTGCTTGAAAATTAATTTCA
PVHY01000998.1:68954-69416(-)  : CTAGAAATATTTTTCCTATACTTTACTGCACACTCCATTCCTTCATACATACAAGTAAAAC
PVHY01015967.1:4675-5136(-)    : TCACATAAGGAATATGCAGTTTTTCCTAATGTGCATGGAACGTTTGTAAAAATTCAACATG
PVHY01008490.1:9516-9977(-)    : AGGAAAAGAGAATCTCTTGGAGTACACCAAGCATAAGATTGAAGGTCAAGGTGAGATGTTC
PVHY01003810.1:13426-13886(-)  : GCTCAAGAATCACCTCTTCCCCTCAAGCTCTATTACTTAACTGGTTTGTAATCAATAATGA
PVHY01009730.1:53713-54174(+)  : CAATGACCTTATTCTAGAAGCATACAGAAAGCTAACTAAAATTTAAATGACTAATAAAAAT
PVHY01012311.1:1227-1691(-)    : AAAAAAAAACCTTTCAAATAGGCTTTAAAACAAATGATCAAAAATGTGATGGATAGTATGA
PVHY01016875.1:11594-12058(+)  : AAAAGTCCTGGTCATTGCACCAAGGCAAGGAAAAAACCAGACATAAAAACTGAGCAGTAAC
PVHY01180168.1:0-381(-)        : AAATTCCAGCTTAATATTAAAAACCTTGAA
PVHY01009930.1:15458-15918(-)  : CCTACTTATAAACTCAAGAGAAATGAGAACGTATGTTCACACACAAAAAAGGGGAAACAAC
PVHY01045966.1:1383-1843(-)    : TGTAGGTTGTCTCCTAAGCTTTGGAAGCTTTGCTTCTCTTAATCACCCATTACTGTCTATC
PVHY01000878.1:41476-41936(-)  : AACGAGAGAGGTTTTTTTTTTAATCTTGTCTGGCATGACTCTTTTGGAATGCAAATGGCGC
PVHY01009677.1:43648-44109(-)  : GCATGAGCCATCAACTCCACTCATGTACACTTACCCGAAAGAAATGAAAATGTAGGTTCAC
PVHY01003949.1:35743-36203(-)  : AATAGCAACGTGCAACGTGCCCAAGATTATTTGAAAGGTGCCAGAATCAGATTTCACCCTA
PVHY01002269.1:76217-76677(+)  : GGCAATGCTTGTAATAAATGGGAATGAAATAATAAGTGATCATCCAATTTGAACATTTACT
PVHY01094534.1:37-497(-)       : TGAGAGAAGGTAATTGGGAATTGGACTTTGGAGATAAGTTGAGAAACTTTCTTCTTTGTGT
PVHY01000828.1:3392-3853(+)    : TGGTGAATTTAATGTTGCGTACCCACATACTGATTTTTAATGATGCACATCTGAAACTTAT
PVHY01002549.1:79670-80132(-)  : TTTTCTATAAGAGTACCCTGTAGTAAATATTTTTGGCTTTGCCAGGCTACATAAAGTCTTT
PVHY01004562.1:2711-3173(-)    : TCCTTTCCACCAGAGGTAATGATTATCCTGACCTAAAATAATTTTTTAATTGAAGTGAAAT
PVHY01012448.1:22154-22616(+)  : AGGCTAATTTATTAACAAAAGAAAGGGGTATTTATAGAGACACAGGACGAGGGTCTGGTGA
PVHY01018318.1:7452-7914(-)    : ATTTTAAGCATAGCTAGGTTAAAAATTACCTAGGATTTTTTTGATGAAGATGTACATATGC
PVHY01030613.1:7241-7942(+)    : AAATAAAATAAAATGCAGCTTCTGCCGCTCAGATTCCCTCACAGACGCATAGCAGTTGCCA
```

```
PVHY01006997.1:68467-68929(-)  : AGCTAAGACTGGGAGTCAATGTATACAGTAATAACAGAATATTTCTTGCAGTAACTGATTT
PVHY01004607.1:35032-35493(+)  : GGGTGGAGAGGAGAGCACAATGGACCATGGCGGAGGACTATGGGTACTTGGCAGTAAGAGA
PVHY01011287.1:22983-23444(+)  : CAATATAGGGCCTTACAGGGCTGAGAAAAGATTTGTACCAATTTTATTTCTTGCAGCTTTT
PVHY01002465.1:6893-7355(-)    : ACCTGTAATCCCAGCTATTTGGGAGGCTGAAGCTAGGAATCACTTGAGTCCGCAGGTTTGA
PVHY01002514.1:48322-48782(+)  : ATCTACTGTTAATATGAGATTTTTAAGAATAAAGTTGCCTAATAATTTTATTGGCTCTTCC
PVHY01009530.1:46209-46671(+)  : GGCTCCAGGGCCTGATGGTCTCGGCATCTCTTAAAAAACAAAAAGCAAGGATAATATTTTT
PVHY01004700.1:67022-67484(-)  : TGGCTACTAGTACTAACTTTATATTGTTTCCCTGTACAAACTAAAAGTATTATGAATAGTG
PVHY01001190.1:157652-158113(- : AACTTATCTTAGAAAATCAGGAACCAGATTACTGAGATTTATATCTGGGGCCATAAATTTT
PVHY01000636.1:36465-36926(+)  : TTGTAGACTGCTCCATTTCCTAGGTAAAGCAGTGTGGAGGTTTATAAAAGTATTGAAAAAT
PVHY01005636.1:17251-17712(+)  : ACAAGAACTCAGCAAAACAGGGTAGAGACCTGAGGACTGGACAGGAAAAGGGGCAGGGCCC
PVHY01018285.1:24866-25326(-)  : GAACACTTCAAGGAGGCAGTACGGCTTCAGAGCAGTGGAAACACCTACAAGAAGAGACTTA
PVHY01027168.1:3480-3941(+)    : GGAGAAAAAAAAAAAAAATCCATGGAGAAATAGAAAGAGCTAACCCTCCACCCCCAAAAAAG
PVHY01054352.1:1347-1808(+)    : GCATGACTTTACATGTCCTAGCTACATCATTCTAATGTGTTGTCCTGATTGATTTACAGTA
PVHY01012201.1:12008-12468(+)  : CAGGCATCATGGCAGGCACCTGTAATCCCAGCTACCTGGGAGGCTGAGCTAGGAATAACTT
PVHY01000554.1:111525-111985(- : GCTTTATGCTGGGGAACAGAGAGAAGGAAGTTTGATGGGGGAAGGGCATAACTGCATTGAA
PVHY01000922.1:123481-123941(- : AAGTTTCTAATTATGTATGAAACCAATATATCTTCATTAAAAGAAGCAAAAAACATTAAAA
PVHY01004425.1:60288-60748(+)  : TGCTTATGAAAAAATTCATTAAATGAATAGTATTGGACATTAAAAAATATTGCTAATGTAT
PVHY01012836.1:2324-2784(+)    : CCAGTAGGGCATAGGATGGTGGTAAGAGGGTATGAAGGTGTCCTTGAGAAACATAAGCAGG
PVHY01067373.1:1777-2237(-)    : GGCAACACAACAACAGCTGACTAGAACAACGGTCTCCGAACCTCCACAACAGGGGAGATGG
PVHY01075250.1:948-1408(+)     : GCAAGAGCCTGGGTTCAGGCTAGAATTGCAAACTCCATCACAAAAAAGTAGGGAGGTGACC
PVHY01003790.1:38072-38532(+)  : AAGTTTGGTTTTGAGCTTCCAGGCAGCAGGAACAGATAAAAGAAAAGTGAGATGTGTGGAC
PVHY01012767.1:3364-3824(+)    : AAATACTCTCATTAAGTAGTTTATAGAGGGATGCATGGATTTATTTATTTTATTATTATTA
PVHY01017727.1:1639-2100(-)    : CGTAGCCAGTGTGGAGTCAGATCCCAGAATGGCAGGCTGAGGGTGACCAGGGGCTGCTGAA
PVHY01007834.1:57293-57754(-)  : CTTTGAATTGACTTGAAAATCTTTTTATGAAAGGTTCTTTAGTATTTCATCTCTGAAAACT
PVHY01042715.1:4832-5294(-)    : TCAACTGACATGACTGAAGCAAAGAAAGAATTAAAGAATGAATTACACCCTGATATTGAAG
PVHY01011504.1:12048-12511(+)  : AGCAATTTAGTGCAATCTTTTAAGAACAATTCAAAATAAATGGAAGAGATAGATTAATGAC
PVHY01035808.1:878-1341(-)     : GAAGGCGCTCGGCTCTTCAACGCCACCGCATAGGTACCCCTGCCCTGCTCGCTGGGCCGTT
PVHY01016752.1:22261-22723(-)  : GCTGAGTTCAACTAGTCCTAAATCAGGCTGGTAGATCACAGAGGCCTGATTTGCAATTACC
PVHY01072951.1:1387-1851(-)    : GGGCAAGTTTCAGGCAGGGGTGAGTCCCTGTCTATGTAAATAAGGACAAATATCCCTATGA
PVHY01013237.1:20453-20915(+)  : TCTCTGATATACTCTTTAAAGGAGTGCAAAGAAAATTCACTATCCAATGCAGCCTACTCAT
PVHY01005556.1:19245-19705(-)  : AAATGCAAAAAGGTGGCAGGTGTACTTCTGGTGTAGATATTCTAGGTATATGGAACAACCA
PVHY01008302.1:25016-25476(-)  : GGGAGAATCCAGGGATAGATAATGATGTAAAATAAAAGAACTCAAGTTGTGGCAATAAATG
PVHY01020077.1:20797-21257(-)  : CTGGGATGACAGAAATGGGAGTTACAATCCCAAGGACACTGGAGTCACTAGATAGTGGAAG
PVHY01016119.1:30218-30678(+)  : TGTTTGTTTGTTTTGAGACTAGTTCTCGCTATTTTTGCCCATGGAACTGCAGGCTCACGTT
PVHY01072528.1:831-1291(+)     : ACCCAAACAATAGCTATTTTTAATGATCAGTTCTCAAATCTCAACTTTGCAGTGTTTCTGC
PVHY01040643.1:4929-5390(-)    : CTGAATTTCCAGATCTAATGTTACAATTTTTCAAAAGAAACCAGAAACCTGGACTTTCAAG
PVHY01001829.1:15787-16247(+)  : TTATTGTTTAGCTATTAATACCATGATCTTTTTTGGTCATCTACGTATATTGTAAAAGTGC
PVHY01004361.1:42264-42724(-)  : TCTGATCTAAGTGGAATGTAATGAATATATATCCCGTCTAAGAAAAAGTGAGAATTTTATT
PVHY01023686.1:8656-9116(+)    : AGGACTTATTGTTCTAAGCCTTAGCCAGAAAGCACTATTTTAAGGTTTGCAGCTGCATTAC
PVHY01012641.1:23680-24146(-)  : CCCTTTACGGGTTCTTAAAAGGGTTAATAGAGATTAATGATTTTTTTTCCAACAAAAGATT
PVHY01015847.1:13399-13868(-)  : TGGTTTTAACTCTGATTTTCTCCTGTTGCACAGTTCGGCCGCCTCTCAGGGGCACATAGAA
PVHY01024847.1:6681-7142(+)    : TTGGCAGAAACAAGTCAGAAGGAAAAAGACAGAAACCAAATGGTTTCACTAATATATGGAC
PVHY01015071.1:21756-22218(-)  : AACCTAGGAAGCCATGGCAGGAACCAGGCATCACGGTTCAAGTCCTGCTCGTGTCAACACA
PVHY01004144.1:90643-91107(-)  : ATAGTTGCTCAGTAAACAACTGGTGATGAACTCAGTCAAGGAGCGGAGCTCTCTCCAAAAG
PVHY01023427.1:8943-9408(-)    : ACACTTTTACTTGCTGGTGGGAATGCAAACTGGTCTAGCCACTATGGAAAGCTGTGTGGAG
PVHY01001716.1:75005-75473(+)  : ATGTGACAATGTCAAAAACACATTTTGGTTGACCTGTTTTCTGCATTTTGATTTCTGCATT
PVHY01006475.1:44649-45110(-)  : ATTTTCTTATGCTTTTTTGAATACTTGAATTCCTAAGCAACTCCCATTCTTTTCTCTTATA
PVHY01000775.1:104571-105031(- : ATGTGCCTATTTTCTTGGAATCCATCCAGCCATGCCAGTGGTTACATTTTAAGTAGGAATG
PVHY01057101.1:1698-2159(+)    : AAATACATGGTTCACATATCTGAGGTACCCAGAGGTTACAAAAGCTCGAATCTAAAAATGC
PVHY01030803.1:11960-12420(-)  : ATCACAATCAGGGTTTGAACCCCTGATCTGACCTTTACCCAATTCACTCGTGATTAAACAG
PVHY01002408.1:33880-34341(-)  : GGGTTGCTGGCAGAATGGACAAAGTGGAGCTCGATTGGGAAGATTTTGTATATAGATGGTT
PVHY01015780.1:7059-7519(+)    : ATGGCAGCCACCAGCAAGCAGTAGCCCAAAACCCAACTCACCACTCACTGCAGCTGGAGGT
PVHY01017180.1:14222-14682(+)  : GCATGGTGGCGCATGCCTGTAAGCGGCAGCCTAGGAAGCCATGGCGGAGACCAGGGATCGC
PVHY01004181.1:11239-11699(+)  : CTCTGTTTCCTTTGACACCTCTTGAAATTATTACCCAAGGAGAACTTCCTACCTAGAACTG
PVHY01016978.1:11379-11839(-)  : CTGCTCTGCAGATGGATGAGTCACTCAGGTATGGTAATTGGCTCTTAGCAGGCTATTAGAA
PVHY01005577.1:45995-46455(-)  : TTCCACTTATACATCCTCTGTTCTGGCTACACCAAACTACTGGGATCAGTCCAAGCATTTT
PVHY01003103.1:12322-12782(-)  : GTCAATGCACACTTCTAAGGAAGCCAAAGTCAGCAGATTGCCACACTCTGAAACCTTTCAT
```

```
PVHY01005700.1:70829-71289(+)  : ACACAAGTCAGATGCTGCTAAGAAGGAGAGTGAGATGAGGACTAGGAAGAATCCAAGAGAT
PVHY01167867.1:29-489(+)       : GCAAAGTGCACAGTCCTGTGTTTTGACAGAGGAGAGGGCTGGGCTGTTTATCCACAGTTAG
PVHY01013412.1:39263-39726(-)  : TGGTGACATGTGCTTGTAAGCAACAGCTTATGGAGTCACAGTGGGGTCTGGGGATAGAGTT
PVHY01036498.1:2252-2712(-)    : GATTTGAAGGCATCTTCTTGGCTTCTTGGTGTGGGTGGAGGCCTATGAAAATATGGACGGG
PVHY01004447.1:65916-66376(+)  : GGTGTTATTGTTCTAAGCCAAATCAATTCAAATAAAATCACTGTGCCACATTCTGAGATCT
PVHY01032970.1:4990-5450(+)    : ATCTTTGTGATAATATAATTAAATTCCAAATACATTTGAACATGTACATAGTTATGCATTC
PVHY01024371.1:4748-5208(-)    : AGTTCCTAATACAGTGCTTCCTCAGTTCCTAATACAGTGCCTACATCATATGATAAGTGCT
PVHY01064486.1:913-1373(+)     : CTATGTTTTTGACTCAGCACTGGCACTGTTCAGCTGCTTTTCTGCAATTCAAAAAGAGCTT
PVHY01018181.1:27724-28184(-)  : TTCTTTGGACCTCTGATCATTTGTCACTAATTTGGAATAGACATGCTACATTTCTTAACTA
PVHY01037172.1:5469-5929(-)    : TTTCTCCTTTTTTCCTCACTACAAGCCATATCACAATCATTATTACTGTCCGTCTACTTCT
PVHY01243375.1:0-398(+)        : GCTGCAGCGCACAGCTAGGCACCCCCCAACCCCACATGTGCCCGCGAGCA
PVHY01012394.1:11145-11605(+)  : TAGTATTCCTGGCAGTGACTTCTGTCTAATCCTTCCTAATGGTGAGAAAAGGAAGCTGATT
PVHY01022271.1:16325-16786(-)  : CTTTTTTTTTGTTTTTGTTTTTGAGACTAGGTCTTTTTATTTTTGCCTGTGGAACCACAGG
PVHY01024968.1:6195-6655(+)    : GAAAGGAAAACTGAACTCTAGGTAAGAGGCACAGCCAACTCGAGGTGTTCTTATTTTCATG
PVHY01004013.1:46042-46502(-)  : ATGTTCCATTAAAACTCAGGCCACTAAGCTTGAAAAGGGTCGTGTCCATTCTTTGTACACT
```

**Figure S3**. Copies of Urop_c from tribe 1. The alignment shows 100 out of 269 copies of the tribe. TSDs are underlined. Adenosine residues in tails are highlighted in yellow.
