## Supplementary material for "A Dimeric SINE Discovered in Shrew Mole is Structurally Similar to Primate Alu": Fig. S4

### Subfamily Urop_c.

PVHY01079167_1:308-653(+) $\mathbf{A_{21}}$
tagaccttaccatcggaggctaggtactgaccctgaaagggaaaacttcaGGCCGGGTACGGTGGCTCGTGCTTGTA
AGCTTCAGCCTATGAAGCAGCCAGGGCTAGGGGTCGTGGTTCGAGTCCCTCCTGGGGCCCGTAGTGAGACCTAATTC
CAAAAAAATAGCTGGGCGAGGTGGCGCACGCCTGTCATCCCAGCTACCTGGGAAGCTGAGGCTGGGGATTACTTGAG
TCTGTGGTACCCCGGGCAACGAGAGCCTGCGGTTCCACAGGCAAAAATAGCAAGACCTAGTCTCAAAAAAAAAAAAA
AAAAAaaaggggaaaacttctggagtgacagtgatgt

PVHY01039115_1:4066-4410(+) $\mathbf{A_{25}}$
gagaacatctcttggtgttttcacacgttctcctcacttcattcagttcaGGCTGGACACAGTGGCTCGTGTCTGTA
AGCTTCAGCCTATGAAGTGGCCAGGGCTAGGGATCGTGGTTCGAGTCCCACTTGGGGCCTGTAGTGAGACCTAGTTC
CAAAAAATAGCTGGGCGTGGTGGCGCACGCCTGTAATCCCAGCTATGTGGGAAGCTGAGGCTGGGGATGGCTTGAAT
CTGTGGTACCCCAGGCAACGAGAACCTGTGGTTCCATGGGCAAAAATAGCAACACCTAGTCTCAAAAAAAAAAAAAA
AAAAAAAAAAacttcattcagttcactagtatgagc

PVHY01003829_1:18915-19259(+) $\mathbf{A_{26}}$
taatgatatatactgttgacgtgtatctctttaacaataaatcacttcaaCCGAACTCGGTGCCTCCCTGCTGTAAG
CTTCAGTTTATGAAGCGGCTGGGGCTAGGGATCGCGGTTCGAGTCCTGCTTGGGGCACGTAGTGAGACCTAGTCCCC
CCAAAATATAACTTGGCGTGGTGGCGCATGCTTATAACCCCAGCTACCTGGGAAGCTGAGGCTGGGGATTGCTTGAG
TCTGGGGTACCCTGGGCAACGTGAACTGCGGTTCCACAGGAAAAAATAGCGAGATCTAGTCTCAAAAAAAAAAAAAA
AAAAAAAAAaaatcacttcaaagaagtgatttccaactacagcatccttctcaccatg

PVHY01014482_1:24069-24423(-) $\mathbf{A_{26}}$
tgggtactgaactttattaatttcttactttcgaaagctagtgtttcccaGGCCGGGCACGGTGGCTCGTGCTTGTA
AGCTTCAGCCTATAAAGTGGCCGGGGCTAGGGATCGCGGTTCAGTCCCGCCTGGGGCCTGTAGCGAGACCTAGTTCC
AAAAAAATAGCTGGGCTCGGTGGTACATGCCTGTAATCCCAGCTACCTGGGAAGCTGAGGCTGGGGATTGCTTGAGT
CTGTGGTACCCCGGGCAACGAGAGCTTGGGGTTCCACGGGCAAAAAATAGGGAGACCTAGTCTCAAAAAAAAAAAAA
AAAAAAAAAAAAAgaaagctggtgttttcccattttcaatccagggaaggaaatgcttta

PVHY01003705_1:57154-57501(+) $\mathbf{A_{27}}$
agaaaggaaaaatagaatggactgcatcaaaattgaaagctcatgtttcaGGCTGGGCACGGTGGCTCGTGCTTGTA
AGCTTCAGCCTATGAAGCGGCCGGGGCTAGGGATCGCTGTTCGAGTCCCGCCTGGGGCCCGTAGCGAGACCTAGTTC
CAAAAAAAATAGGTGGGCGTGGTGGTATGCACCTGTAATCCTAGCTACCTCGGAAGCTGCGGCTGGGGATTGCTTGA
GTCTGTGGTACGCCGGGCAACGAGAGCCTGGGGTTCCATGGGCAAAAAATAGCAAGACCTAGCCTCAAAAAAAAAAA
AAAAAAAAAAAAAaaagctcatgtttcaaatgacagtatgaacaaagtgaacagaatgggagaaa

PVHY01001329_1:101916-102275(+) $\mathbf{A_{37}}$
tatgaagccatcactactttgaaatactacctgaccaagatcttaccaatgcaGCCGGGCATGGTGGTGCATGCCTG
TAAGCGGCAGCCTACGAATCCACGGCAGGGACCAGGGATCACGGTTTGAGTCCTGCTCCAGGCAACGTAGTGAGGCC
TAGTTCCAAAAAAAATAGCTGGGCGTGGTGGCAAGTGCCTGTAATACCAGCTACCTGGGAAGCTGAGGCTGGGAATT
GCTTGAGTCCTGGTACCCCAGGCAACCTGAGCCTGCGGTTCCATGGGCAAAATAGCGAGACCTAGTCTCAAAAAAAA
AAAAAAAAAAAAAAAAAAAAAAAAAAAAtcttaccaatgcaattataatctaatttcttttgatatagctttaac

PVHY01000332_1:85378-85721(-) $\mathbf{A_{45}}$
ctgaagtaaggtcagctgtggtttgttccaagattcatttgtttgtttctGCCAGGCACCATGCTTGTGCCTGTAAG
CTTCAGGCTATGAAGCGGCAGGGGCTAGGGATGGCGGTTCGAGTCCCACCTGGGGCCTGTAGTGAGACTTAGTTCCA
AAAAAATAGCTGGGCGTGATGGTGTGTACCTGTAATCCCAGCTACCTGGGAAGCTGAGGCTGGGGATTGCTTGAGTT
TGTGGTAACCTGGGCAACGAGAGCCTGCGGTTCCACGGGCAAAAATAGCAAGACCTAGTCTCAAAAAAAAAAAAAAA
AAAAAAAAAAAAAAAAAAAAAAAaaagatccatttgtttgtttccgagtggagggcaaattttaa

PVHY01008294_1:47328-47679(-) $\mathbf{A_{53}}$
aacccaaacaaaggttcctgatgagtgcttggtaaagactccagtttgcagGCCGGGCATGGTGGCGCATGCCTGTA
AGCGGCAGCCTAGGAACTCGTGGCAGGGGCCAGGGATCATGGTTCGAGTCCCATCTGGGGCACATAGCGAGACCTAG
TTCCAAAAAAATATAACTGGGCACGGTGGCGCGCGCCTGTAACCCCAACTACCTGGGAAGCTGAGGCTGGGGATTGC
TTGAGTTCGTGGTACCCCCGGCAGTGTGAGCCTGCGGCACAAATAGGGAGACCTAAGTCTCAAAAAAAAAAAAAAAA
AAAAAAAAAAAAAAAAAAAAAAAAAAAAAAAAAAACCCAAAaaactccggtttgcagcagaggcacctccagagg

PVHY01000312_1:73138-73481(-)        $\mathbf{A}_{56}$

aacagttagaggtgatcacacgaggcttgacaacagaaagacttgatagaGCCGAGCAGGTGGCGCGTCCCTGTAAG
CTTCAGCCTATGAAGCGGCAGGGGCCAGGGATTGCAGTCCAAGTCCCGCCAGGGGCACGTAGTGAGACCTAGTTCCA
AAAAAATAGCTGGGTGTGGTGGCATGTGCCTGTAATCCCAGCTACCTGGAAAGCTGAGACTGGGGATTGCTTGAGTC
TGTGGTACCCTGGGCAACATGAGCCTGCAGTTCCACAGGCAAAAATAGCAAGACCTAGTCTCAAAAAAAAAAAAAAA
AAAAAAAAAAAAAAAAAAAAAAAAAAAAAAAAAAAAAAAAAgacttgctacctctccactcacaagttgtgtgattt
gcacattgttttagatccttgccttagtttttgct

PVHY01006787_1:60621-60970(+)        $\mathbf{A}_{57}$

ttgaattatttttgactttcaggaagcaaaaaaaaaaaaaaaaaaatcaaGACCGGGCACTGTGGCACGTTCCTGTA
AGAGGCAGCCTAGGAAGCCACGGTGGGACTGGGGATCGTGGTTTGAGTTCCAGTAGGGGCAAAGTAGTGAGACCTAG
TTCCAAAAAAATAGCTGGGTGTGTTGGCACATGCCTGTAATCCCAGCTACCTGGGAAGCTGAGGCTGGGGATTGCTT
GAGTCCGTGGTACTCACAGGCAATGTGAGCCTGCGGTTCCATGGGCAAAAATAGCAAGACCTAGTCTCAAAAAAAAA
AAAAAAAAAAAAAAAAAAAAAAAAAAAAAAAAAAAAAAAAAAAAAAAAtcaacactgaatgggattataaccaaatg
tgtaagcaattaaggaaacttgtttttttaaa

PVHY01009329_1:34967-35319(+)        $\mathbf{A}_{59}$

cactcagatatagaaggcaaatattaacagaaataaaaggaagaaaaaagGGCCGGGCACGGTGGCTCGTGTCTGTA
AGCTTCAGCCTATGAAGTGGCCAGGGCTAGGGATCGCGGTTCGAGTCCCACCTGGGGCCCATAGTGAGACCTAGTTC
CAAAAAATAGCTGGGCGTGGAGGCTTACGCCTGTAATCCCAGCTACCTGGGAAGCTGAGGCTGGGGATTGCTTGAGT
CTGTGGTACCCCAGGCAACAAGAGCCTGTGGTTCCTGGGCAAAAATAGCGAGACCTAGTCTCAAAAAAAAAAAAAAA
AAAAAAAAAAAAAAAAAAAAAAAAAAAAAAAAAAAAAAAAAAAAcaaacaaaaaaaaagaaataaaaggaagaaaaa
aaaaagtgagcagtgtaatactagttgaagacattaataagaatgaacaaatcttccagacacaggac

PVHY01000061_1:336419-336772(+)        $\mathbf{A}_{61}$

taaagcacttgtttgtcattctctgaaggagttgcaaatctggcaattgtGCCGGGCACGGTGGCTCGTGCTTGTAA
GCTTCAGCCTATGAAGCGGCCGGGGCCAGGGATCGCGGTTCGAGTCCTGCCTGGGGCCCGTAGTGAGACCTGGTTCC
AAAAAAACAGCTGGGCGTGGTGGCACACGCCAGTAATCCCAGCTACCTGGGAAGCTGAGGCTGGGGATGGCTTGAGT
CCGTGGTACCCCGGGCAACGAGAGCCTGCGGTTCCACAGGCAAAAATAGCGACACCTAGTCTCAAAAAAAAAAAAAA
AAAAAAAAAAAAAAAAAAAAAAAAAAAAAAAAAAAAAAAAAAAAAAAtctggcaattgttggcttgaacttgtggta
aacattgagccaacatttcatccattacaaatttccctgggagattactttatgatacccat

PVHY01007010_1:2582-2940(-)        $\mathbf{A}_{80}$

tgtggagacaaggaagtcgctgtgcactctgggtaagaatgtgagttgggGGCTGGGCACGGTGGTGCATGCCTATA
AGCTGCAGCCTAGGAAACCGTGGCGGGAACCAGGAATCGCGGTTCGAGTCCTGCCAGGGGCACCGTAGCGAGACCTA
GTTCCAAAAAATTAGCAGGGCTTGGTGGCGCGTGCCTGTAATCCCAGCTACCTGGGAAACTGAGGATGGGGATTGCT
TGAGTCTGTGGTACCCCAGGCAACATGAGCCTGCGGTTCCACAGGCAAAAATAGCAAGATCTAGTCTCAAAAAAAAA
AAAAAAAAAAAAAAAAAAAAAAAAAAAAAAAAAAAAAAAAAAAAAAAAAAAAAAAAAAAAAAAAAAAAAAAgtgagt
tggtacagccaccagggaaaacaataaagaagcgcttca

PVHY01010040_1:34589-34952(+)        $\mathbf{A}_{88}$

tatctctgtttataaccacttcatttggttattaaaatgcattagtactaGGCCGGGCACGGCGGCTCGTGTCTGTA
AGCTTCAGCCTATGAAGCGGCCTTCATAATGGTGCTAGGCATCGCAGTTCGAGTCCTGCCTGGGGCCCATAGCGAGA
CCTAGTTTCAAAAAATATAGCTGGGTGTGGTGGTGCACACCTGTAATCCCAGCTACCTGGAAAGCTGAGGCTGGGGA
TTGCTTGAGTCTGCGGTACCCCAGGCAACGAGAGCCTGTGGTTCCATGAGCAAAAACAGTGAGACCTAGTCTCAAAA
AAAAAAAAAAAAAAAAAAAAAAAAAAAAAAAAAAAAAAAAAAAAAAAAAAAAAAAAAAAAAAAAAAAAAAAAAAAAA
AAAAAAAgcattagcAAAGTAATCATGTgtgaattaaaaaattgtttatttctgtATGTTAATGTTTTCTATTCTAC

PVHY01000966_1:124831-125176(+)        $\mathbf{A}_{89}$

AgttctctattcaacccttaggacacaaaggtaaaagaagggaaccaGACCTGGACACGGTGGCACGTGCCTGTAAG
CGTCAGCCTATGGAGCATGGCTGGAACAGGGGATCGGCGGTTTGAGTCCCACCCAGGATAACATAGTGAGACCTAGT
TCCAAAAAAATAGCTGGGTGTGGTGGTGCGCATCTGTAATCCCAGCTACCTGGGAAGCTGAAGCTGGGGGTTGTTTG
AGTCTGTGGTACCCCGGGCAATGTGAGTCTGTGGTTCCAAGAGCAAAAATAGCGGAACTTAGTCAAAAAAAAAAAAA
AAAAAAAAAAAAAAAAAAAAAAAAAAAAAAAAAAAAAAAAAAAAAAAAAAAAAAAAAAAAAAAAAAAAAAAAAAAAg
aaaggaaaaaaggaaacagatttaataggcttcatctgatgagcctaggat

PVHY01033845_1:9058-9541(+). **$A_{90}$**

```
ttatctgcacttaagtgtataataagatttgcttcaaaacagtggcaacaGCCGGGCACGGTGGCTCGTGCCTGTAA
GCTTCAGCCTATGAAGCGGCCGGGCCTAGGGATCACGGTTCGAGTCCCGTCTGGGGCCCGTAGCGAGACCTAGTTCC
AAAAAATAGCTGGGCGTGGTGGCGCGCGCCTGTCATCCCAGCTACCTGGGAAGCTGAGGCTGGGGATTGCTTGAGTC
TGTGGAACCCCGGGCAACGAGAGCCTGCGGTTCCACGGGCAAAATAGCGAGACCCAGTCTCAAAAAAAAAAAAAAA
AAAAAAAAAAAAAAAAAAAAAAAAAAAAAAAAAAAAAAAAAAAAAAAAAAAAAAAAAAAAAAAAAAAAAAAaaaaca
gtggcaacaaatctctctagaaagctcaacagaacctactatgtagtagcaatcacat
```

### Subfamily Urop_b.

PVHY01026755_1:13314-13669(+) **$A_{21}$**

```
taatagccaaaaaatgaaagacaaaatgttcatcaatatgtgagcaaatgGGCTGGGCACGATGGCATGTGCCTTTA
ATTGCAGCCTCCAAAGTGAGGCCAAGACTAGGAATTGCTGGAGTCCGCGGTTGGAGACCCACCCGAGGCAGCATAGC
AAGACCTAGTCTCAAAAAATAGCTAGGCGTGGTGGCGTGTGCCTGTAATCCCAGCTGCCTGGGAAGCTGAGACTGGG
GATCACTTGAGTTCGTGGTACCCCAGGCAACGTGAGCCTGTGGTTCTCTGGGGAAAATAGTGAAACCTAGTCTCAAA
AAAAAAAAAAAAAAAAAAgtgagtaaataaattaattaaaatacaat
```

PVHY01003356_1:14114-14468(-) **$A_{29}$**

```
aggacgctgacctaactattgaactgtaactttttaaaaatactgaggataGTCAGTCATGGTGGCATGCGGATTTAA
TCCCAGCCCAGGACAGGAAGCTGAGGCTAGGAATCACTTGCGGCCACTGGTTCAAGACCTGCCTGGGGCAACATAGC
GAGACCTAGTCTCAAAAAATAGCTAGGCATGGTGGTGCGTGCCTGTAATCCCAGCTACCTGGGAGACTGAGACTAGG
GATCATTTGAGTCCAGGGTACCTCGGGCAACAAGACTTTATGGTCCCTGGGCAACATCGCGAGACCTAACCTCAAAA
AAAAAAAAAAAAAAAAAAAAAAAAAtactgaggatagtgtttgctt
```

PVHY01005495_1:44137-44492() **$A_{61}$**

```
gaagccagatcatgagataggactctctgataaaaggaaatagcttcccaGCCGACTGTGGTGACACACAGGTTTAA
TCTCAGCCTAGGACAGGCGACTGAGACTGGGAATTACTTAAGTCTGCAGGTTTGAGACCCACCCAGGGCAACATAGC
GAGACCTAGTCTCAAAAAATAGCTGGGTGTGGTGGTATGCACCTATAATCCCAGCTACCTGGAAGGCTGACACTGGG
GACCGCTTGAGTCTGGGATATCCTGGGCAACGAGAGTCTGTGGTTCCCAAGGCAACAAAGTGAGACCTAGCCTCAAA
AAAAAAAAAAAAAAAAAAAAAAAAAAAAAAAAAAAAAAAAAAAAAAAAAAAAAAAAAAggaaacagcccCCTGGATA
AATGACTGAATCTGGGACTGGGCAGGAGAAACACAACATGAACTTGATATGTCGTT
```

PVHY01006540_1:70565-70916(+) **$A_{65}$**

```
aaccatgaaaactgcctaaatctctttcagtggatcaaggagattatgtACCAGGCACAGTGGCACATACCTTTAAG
CACAGCCTAGAGCGAAGCTGAGACTAGGAATCACTTGAGTCCGTGGGTTCCATACTGCCCGGGGCAACATAATGAGA
CCTATTCTCAAAAAATAGCTGGGCTTGGTGATGTGCGCCTGTAATCCCAGCTACCTGGGAAACTGAGGCTGGGGATT
GCTTGAGTCCGTAGTATTCTGACCAACGTGAGCCTGTGGTTCCCCAGGCAAAATAGCGAGACCTAGTCTCAAAAAAA
AAAAAAAAAAAAAAAAAAAAAAAAAAAAAAAAAAAAAAAAAAAAAAAAAAAAAAAAAAggagattatgcatatacac
accatggaatattatgttactataagaaaaaatgggatgtaaattgtagggaatttgagagaatta
```

PVHY01014619_1:15438-15794(+) **$A_{158}$**

```
cagtaacctacgaaacgcagtctagggggtctagaaaaacaactgtataGCCAGGCACAGTGGCACGTGCCTTTGA
GCGCAAACCTATGGAGCGCAGCTGGGACTGTGAATTGCTTGAGTCTGAGGGTTTGAGACCCGCCCAGGCAACATAGT
GAGACCTAGTCCCAATAAATAGCTGGGTGTTGTGGCACGCACCTGTAATCCCAGCTACCCGGGAAGCTGAGGCTGGG
GATTGCTTGAGTCTGTGGTACCCCCAGCAACGTGAGCCTGCAGTTCCCCAGGCAAAAATAGTGAGACCTATTCTCAA
AAAAAAAAAAAAAAAAAAAAAAAAAAAAAAAAAAAAAAAAAAAAAAAAAAAAAAAAAAAAAAAAAAAAAAAAAAAAA
AAAAAAAAAAAAAAAAAAAAAAAAAAAAAAAAAAAAAAAAAAAAAAAAAAAAAAAAAAAAAAAAAAAAAAAAAAAAA
AAGAAAAGAAAAAGAAaaagtaaaacaacggtttagaaaaacaacccatgactactgactcactttaatggtatgta
atcctgtttctccttg
```

**Figure S4.** Examples of Urop_c and Urop_b copies with long poly(A) tails. Flanking sequences are shown in lowercase. TSDs are underlined. Sequences corresponding to the promoter boxes A and B are highlighted in blue and yellow, respectively. The length of pure poly(A) sequence is specified after the sequence name.
