## Supplementary material for "A Dimeric SINE Discovered in Shrew Mole is Structurally Similar to Primate Alu": Fig. S5

PVHY01002910.1:36035-36780(+). $\mathbf{A}_{140}$

agttcccaacaacagatgatcagtttaaaaagatgtggtatataGGCCAGGCATGGTGGTGCATGCCTGTAAGCGGC
AGCCTAGCAAGCCACGGCTGGAGCTAGGGATCGCGGTTCGAGTCCCCCTGGGGGCACATAGCAAGACCTAGTCCCAA
AAAATAGCTGGGTGTGGTGGTGTAGTGGTGCATGCCTGTAATTCCAGTTACCTGGGAAGCTGAGGCTGGGGATTGCT
TGAATCTGTGGTACCCTGGGCAACGTGAGCCTGTGATTCCACAGGCAAAAATAGTGAGACCTAGTCTCAAAAAAAAA
AAAAAAAAAAAAAAAAAAAAAAAAAAAAAAAAAAAAAAAAAAAAAAAAAAAAAAAAAAAAAAAAAAAAAAAAAAAAA
AAAAAAAAAAAAAAAAAAAAAAAAAAAAAAAAAAAAAAAAAAAAAAAAAaaaaagatgtggtatataaacacaatgg
aataatggaatattattcaaatataaaatacaagagaaatcctgcctatgtgacaacgtggatgaac

PVHY01006121.1:55550-56292(-). $\mathbf{A}_{140}$

atgtgcaattcagaaatcatttatttttaaaaaagaaaagcttGGGCCAGGCATGGTGGTGCATGCCTATAAGTGGC
AGCCTAGGAAGCCATGGCAGGGGCCAGGGATCACGGTTCGAGTCCTGCTGGGGGAGCATAGTGAGACCTAGTTCCAA
AAAATATATAGCTGGGTGTGGTGGCATGAGTCTGTAATCCCAGCTACCTGGGAAGCTGAGGCTGGGGATTGCTTGAG
TCTTTGGTACTCTGGGCAATGTGAGCCTGTGGTTCCACAGGCAAAAGTAGCAAGACCTAGTCTCAAAAAAAAAAAAA
AAAAAAAAAAAAAAAAAAAAAAAAAAAAAAAAAAAAAAAAAAAAAAAAAAAAAAAAAAAAAAAAAAAAAAAAAAAAA
AAAAAAAAAAAAAAAAAAAAAAAAAAAAAAAAAAAAAAAAAAAAAAAAAAAGAAAGaaagaaaagcttatggggtaa
aatccaaaagaacaataaaggtaaaaac

PVHY01009249.1:41389-42125(-). $\mathbf{A}_{170}$

aattccaataaaaatttcaatggcatttcttaagaaatagaacaGGCCGGGCATGGTCACGCATGCCTGTAAGCAGC
AACCTaagaaaCTGCGGCCAGATCCAGGGACTGTGGTTCGAGTCCTGCCAAGGGTATGTAATGAGACCTAGTTCCAA
AAAAATATAGCTGGGTGTGGTGGCATGCGCCTAGAATCCCACCTACCTGGGAAGCTGAGACTGGGGATTGCTTGAGT
CTGTGGCACCCTGGGCAATGTGAACCTGCGGTTTCACAGGCAAAAATAGCGAGACCTAGATCAAAAAAAAAAAAAAA
AAAAAAAAAAAAAAAAAAAAAAAAAAAAAAAAAAAAAAAAAAAAAAAAAAAAAAAAAAAAAAAAAAAAAAAAAAAAA
AAAAAAAAAAAAAAAAAAAAAAAAAAAAAAAAAAAAAAAAAAAAAAAAAAAAAAAAAAAAAAAAAAAAAAAAAAAAa
agaaagaaaaagaacaaatattactgaaattcatatggaataataaaaaaacttaaatagtc

PVHY01007196.1:33009-33744(+). $\mathbf{A}_{159}$

tcttgatgatgattatttacattgaacatcaaaagcaaaggcaaGGCCAGGCACGGTGGCTCGTGCTTGTACGCTTC
AACCTATGAAGCAGCCGGGGCTAGGGATCGCAGTTCGAGTCCCGTGTGGGGCCAGTAGTGAGACCTAGTTCCAAAAA
AATAGCTGGGCACGGTGGTGCGCGCCTGTGATCCCAGCTACCCGGGAAGCTGAGGCTGGGGATTGCTTGAGTCTGTG
GTACCCCCGGCAACGAGAGCCTGTGGTTCCACGGGCAAAAATAGCCAGACCTAGTCGCAAAAAAAAAAAAAAAAAAA
AAAAAAAAAAAAAAAAAAAAAAAAAAAAAAAAAAAAAAAAAAAAAAAAAAAAAAAAAAAAAAAAAAAAAAAAAAAAA
AAAAAAAAAAAAAAAAAAAAAAAAAAAAAAAAAAAAAAAAAAAAAAAAAAAAAAAAAAAaaaagcaaaggcaacaga
agcaaaaacaaacaaataaactaaaaaacttcacatcaaaggaaaccatcaattaaa

PVHY01040674.1:6218-6951(+). $\mathbf{A}_{170}$

aaacagtctaagctaagatctgacttaaaaaatggactaatctGGGCCGGGCACAGTGGCTCATGCCTGTAAGCTTC
AGCCTATGAAGCGGCCGGGGCTAGGGATCACGGTTCAAGTCCTGCCTGGGGTCCGTAGTGAGACCTAGTTCCAAAAA
ATAGCTGGGTGTGGTGGCGCACGCCTGTAATCCCGGCTACCTGGGAAGCTGAGGCTGGGGATTGCTTGAGTCTGTGG
TACCCTGGGCAACGAGAGCCTGCGGTTCCACAGGCAAAAATAGCAAGACCTAATCTAAAAAAAAAAAAAAAAAAAAA
AAAAAAAAAAAAAAAAAAAAAAAAAAAAAAAAAAAAAAAAAAAAAAAAAAAAAAAAAAAAAAAAAAAAAAAAAAAAA
AAAAAAAAAAAAAAAAAAAAAAAAAAAAAAAAAAAAAAAAAAAAAAAAAAAAAAAAAAAAAAAAAAAAAAAAggact
aatctcctgtcttatctgagaccagaaataggcaaataaggaaagttgtttgtcaa

PVHY01000814.1:58317-59052(-). $\mathbf{A}_{169}$

aaaattaccgtaaatatgtttatatttaaaaaacaggaaaattggGCCGGGCACGGTGGCTCATGCCTGTAAGATTC
AGCCTATGAGGCGGGTGGGTCTATGGATCACGGTTTGAGTCCCGCCAGGGGCCCGTAGTGAGACCTAGTTCCAAAAA
GATAGCTGGGTGTGGTGGTGTGCGCCTGTAATCCCAGCTACCTGGGAAGCTGAGGCTGGGGATTGCTTGAGTTCGTG
GTACCCAAGGCAACGAGAGCCTGCGGTTCCACTGGCAAAAATAGCGAGACCTAGTCTCAAAAAAAAAAAAAAAAAAA
AAAAAAAAAAAAAAAAAAAAAAAAAAAAAAAAAAAAAAAAAAAAAAAAAAAAAAAAAAAAAAAAAAAAAAAAAAAAA
AAAAAAAAAAAAAAAAAAAAAAAAAAAAAAAAAAAAAAAAAAAAAAAAAAAAAAAAAAAAAAAAAAAAAAAAAGAAA
AAATcaggaaaattggtttttattagtttggtaatttccaagaaatatgagatgaaa

PVHY01011285.1:17940-18674(-). $\mathbf{A}_{154}$
tgacaaactttactttttgtggagaataaaagtagtaacaaagGGCCGGGCACGGTGGCTCTTGCTTGTAAGCTTC
AGCCTATGAAGCGGCCGGGGCTAGGGATCGCAGTTTGAGTCCCGCCTGGGGCCCGTAGTGAGACCTAGTTCCACAAA
ATAGCTGGGCGTGGTGGCGCGCGCCTGTAATCCCAGCTCCCTGGGAAGTTGAGGCTGGGGATTGCTTGAGTCTGTGG
TACCCCGGGCAACAAAAGCCTGCGGCTCCACAGGCAAAAATAGCAAGACCTAGTCTCAAAAAAAAAAAAAAAAAAAA
AAAAAAAAAAAAAAAAAAAAAAAAAAAAAAAAAAAAAAAAAAAAAAAAAAAAAAAAAAAAAAAAAAAAAAAAAAAAA
AAAAAAAAAAAAAAAAAAAAAAAAAAAAAAAAAAAAAAAAAAAAAAAAAAAAAaaaagtagtaacaaagtctggccc
ccaaatcactgttaacatagtagaatagcattaagaatttgagcaaaaatttcaaa

>15 PVHY01026942.1:6682-7417(+). $\mathbf{A}_{159}$
acaccagtactgggggccacagccccctaacagaagcagcaacaGGCCGGGCACGGTGGCTCGTGCCTGTAAGTTTC
AGCCTATGAAGCAGCCAGGGCTAGGGATTGCGGTTCGAGTTCCGCCTGGGGCCTGTAGTGAGACCTAGTTCCAAAAA
AATAGCTGGGCGTGATGGCGCGCGCCTGTGATCCCAGCTACCTGGGAAGTTGAGGCTGGGGATTGCTTGAGTCTGTG
GTACCCCAGGCAACATGATCCTGCGGTTCCACAGGCAAAAATAGCCAGACCTAGTCTCAAAAAAAAAAAAAAAAAAA
AAAAAAAAAAAAAAAAAAAAAAAAAAAAAAAAAAAAAAAAAAAAAAAAAAAAAAAAAAAAAAAAAAAAAAAAAAAAA
AAAAAAAAAAAAAAAAAAAAAAAAAAAAAAAAAAAAAAAAAAAAAAAAAAAAAAAAAAAAAAAGAAAAAGAAAAGAA
AAagaagcagcaacaaaaataacaaactggatgttaccaaaatgaaaaactattatg

PVHY01001821.1:7336-8074(+). $\mathbf{A}_{153}$
agctagaaaagtttgggagaaagaattaaaaagaagaaaagtaaGGCCGGGCACGGTGGCTCGTGCCTATAAGCTTC
AGCCTATGAAGCGGCCTGGGCCAGGGATCGCGGTTCGAGCCCCGCCTGGGGTCCGTAGCGAGACCTAGTTCCAAAAA
AAAAATAGCTGGGTGTGGTGGTGTGCTCCTGTAATCCCAACTACCTGGGAAGCTGAGGCTGGGGATTGCTTGAGTCT
GTGGTACCCCGGGTAACGAGAGCCTGCGATTCCACGGGCAAAAATAGCAAGACCTAGTCTCAAAAAAAAAAAAAAAA
AAAAAAAAAAAAAAAAAAAAAAAAAAAAAAAAAAAAAAAAAAAAAAAAAAAAAAAAAAAAAAAAAAAAAAAAAAAAA
AAAAAAAAAAAAAAAAAAAAAAAAAAAAAAAAAAAAAAAAAAAAAAAAAAAAAAAAAAAAGAAGAAGaagaagaaaa
gtaaaggagagggtgaggagaagcctggaggacgcacatggagcagaaacatgaagaaaa

PVHY01011921.1:34023-34759(+). $\mathbf{A}_{170}$
taaacataggagttaaaacagtgcataaaaatcataaaataCCAGGCCGGGCATGGTGGCTCGTGCCTGTAAGCTTC
AGCCTATGAAGTGGCTGGGGCTAGGGATAGAAGTTTGAGTCCTGCCCGGGGCCTGTAGCGAGACCTAGTTCCAAAAA
ATACAGCTGGGCGTGGTGGTGTGTGTCTGTAATCCCAGCTAACTGGGAAGCTGAGGCTGAGGACTGCCTGAGTCTGT
GGTACCCAGGGCAACGAGAGCCTGCGGTTCCACGGGGAAAAATAGCTAGACCTGGTCTCAAAAAAAAAAAAAAAAAA
AAAAAAAAAAAAAAAAAAAAAAAAAAAAAAAAAAAAAAAAAAAAAAAAAAAAAAAAAAAAAAAAAAAAAAAAAAAAA
AAAAAAAAAAAAAAAAAAAAAAAAAAAAAAAAAAAAAAAAAAAAAAAAAAAAAAAAAAAAAAAAAAAAAAAAAAAGA
AAAAAAGAAAAaaaaatcataaaataactcaaattgaaagacaaaggcacattcaaac

PVHY01012450.1:32753-33487(-). $\mathbf{A}_{143}$
gtaccttcttataagaagctatgatagtaaacactagctattaatGTCAGGCACGGTGGCTCGTGCCTGTAAGCTTC
AGCCTATGAAGCGGCCAGGTTAGGGATCGCGGTTCGAGTCCTGCCTGGGACCCGTAGTGAGACCTAGTTCCAAAAAA
ATAGCTGGGCGTGGTGGCACGTGCCTGTAATCCCAGCTACCTGGGAAGCTGAGGCTAGGGATTGCTGGAGTCTGTGG
TACCCCAGGCAATGTGAGCCTGCGGTTCCACCGGCAAAAATAGCAAAACCTAGTCTCAAAAAAAAAAAAAAAAAAAA
AAAAAAAAAAAAAAAAAAAAAAAAAAAAAAAAAAAAAAAAAAAAAAAAAAAAAAAAAAAAAAAAAAAAAAAAAAAAA
AAAAAAAAAAAAAAAAAAAAAAAAAAAAAAAAAAAAAAAAAAAaaacactagctattaatcatttattgtacttaat
aaaaaacaaatcagaaataataaacagcttctagtccaatttatgtaatactaaaat

PVHY01009322.1:55395-56130(+). $\mathbf{A}_{150}$
gaaagagagagaaaaaccaatttgaagttaaaactgaaggggaaGGCTGGTTACGGTGGCTCATGCCTGTAAGCTTC
AGCCTATGAAGCGGCCGGGGCCAGGGATCGCGGTTCAAGTCCTGCCTGGGGCCCGTAGTGAGACCTAGTTCCAAAAA
AACAGCTGGGTGTGGTGGCGCTCGCCTGTCATCCCAGCTACCTGGGAAGCTGAGGCTGGGGGTTGCTTGAGTCTGTG
GTACCCCAGGCAACGAGAGCCTGCGGTTCCACGGGCAAAAATAGCAAGACCTAGCCTCAAAAAAAAAAAAAAAAAAA
AAAAAAAAAAAAAAAAAAAAAAAAAAAAAAAAAAAAAAAAAAAAAAAAAAAAAAAAAAAAAAAAAAAAAAAAAAAAA
AAAAAAAAAAAAAAAAAAAAAAAAAAAAAAAAAAAAAAAAAAAAAAAAAAaaaactgaaggggaaaataatttacaa
taagacaagtaagataaattatatggggaaaaaacttgacaagaaatgtgtgagatg

PVHY01008365.1:7625-8363(+). **A**$_{156}$

ccaaactagtcctccttgaaatttacaattctaaagctacagggGGCTGGGCATGGTGGTTCATGCCTGCAAGCGGC
AGCCTAGGAAGCCATGGCAGGGGCCAGGGATTGTGGTTCAAATCCTACTGGGGTCACGCAGTGAGACCTAGCCCCCA
AAAAATAGCTGGGTGTGGTGGTGCATGCCTTTAATCCCAGCTACCTGGCAAGCTGAGGCTGGGGATTGCTTGAGTCT
GTGGTACCCCAGGCAACATGATCCTGCAGTTCCATAGGCAAAAATAGCAAGACCTAGCCTCAAAAAAAAAAAAAAAA
AAAAAAAAAAAAAAAAAAAAAAAAAAAAAAAAAAAAAAAAAAAAAAAAAAAAAAAAAAAAAAAAAAAAAAAAAAAAA
AAAAAAAAAAAAAAAAAAAAAAAAAAAAAAAAAAAAAAAAAAAAAAAAAAAAAAAAAAAAaaagctacagggaagta
attacattctcctagccattatcaatatatgtatcagttatgttaatggcaagcaccact

PVHY01018992.1:7438-8175(+). **A**$_{159}$

tttttgtgatagtgtatttttgtcctagttaagaaattattgAGGCTGGGCATGGTGGCACATGCCTGTAGGTGGC
AGCCTAGGAAGCCATGGCAGGAGCCAGGGATCGTGGTTTGAATCCCACCGGGGCACGTAGCAAGACCTAGTCCCCAA
AAAATAGCTGAGTATGGTGGCGCATGCTTGTAATCCCAGCTATCTGGGAAGCTGAGGCTGGGGATTCTTTGAGTCTG
TGGTACCCCAGGCAACGTGAACCTGCAGTTCCATGGGTAAAAATAGCGAGACCTAGTTTCCAAAAAAAAAAAAAAAAA
AAAAAAAAAAAAAAAAAAAAAAAAAAAAAAAAAAAAAAAAAAAAAAAAAAAAAAAAAAAAAAAAAAAAAAAAAAAAA
AAAAAAAAAAAAAAAAAAAAAAAAAAAAAAAAAAAAAAAAAAAAAAAAAAAAAAAAAAAAAAAAAAAGAAATAAAAG
AAAAGAAAATAAAaagaaattattgcctattgcctgttataagaagggacctactttttt

PVHY01000850.1:160789-161520(+). **A**$_{164}$

ctaacaaaatgaaatacagaccaacaataaagaaataagtaaacaGCCAGGCATGGTAGCTTGTGGCTGTAAGCGTC
AGCCTATGAAGCAGCCGAGGCCAGGGATCACGGTTCAAGTCCTGCCTGGGGCGTATAGAGAGACCTAGTTCCAAAAA
AATAGCTGGGCATGGTGGTGCGCGCCTGTAATCCCAGTTACCTGGGAAGCTGAGGCTGGGGATTGCTTGAGTCTGTG
GTACCGCAGGCAATGAGAGCCTGCAGTTCCATGGGCAAAAATAGCGAGTAGTCTCAAAAAAAAAAAAAAAAAAAAAA
AAAAAAAAAAAAAAAAAAAAAAAAAAAAAAAAAAAAAAAAAAAAAAAAAAAAAAAAAAAAAAAAAAAAAAAAAAAAA
AAAAAAAAAAAAAAAAAAAAAAAAAAAAAAAAAAAAAAAAAAAAAAAAAAAAAAAAAAAAAAAAAGAAAGAAAATta
agtaaacatagagttgtttcttcaaaaagcacaataaaattcccatcttctcaa

PVHY01012762.1:37864-38604(+). **A**$_{144}$

ttagccaaactcacacagaacacaaaatcagaaaggaaagagaGGGCTGGGCACGGTAGCTCGTGCCTGTAAGCGTC
AGCCTATGAAGCGGCCAGGGCGAGGGATCGCGGTTCGAGTCCCAACTGGGGCCCGTAGCAAGACCTAGTTCCAAAAA
AATATAACTGGGCGTGGAGGTGGTGCGCGCCTGTAATCCCAGCTACGTGGGAAGCTGAGGCTGGGGATGGCTTGAGT
CTGTGGTACCCCAGGAAACAAGAGCCTGCGGTTCCACGGGCAAAAACAGCAAGACCTAGTCTCAAAAAAAAAAAAAA
AAAAAAAAAAAAAAAAAAAAAAAAAAAAAAAAAAAAAAAAAAAAAAAAAAAAAAAAAAAAAAAAAAAAAAAAAAAAA
AAAAAAAAAAAAAAAAAAAAAAAAAAAAAAAAAAAAAAAAAAAAAAAAAAAAagaaaggaaagagaaattacaagag
atactacagaaatgagaaggattttaagagagtaattccacatgtcaacacattggacaatc

PVHY01013167.1:17612-18347(+). **A**$_{147}$

atgctaaattaatgttcagattctttactaaaagtaagtgcattGGCCGGGCACGGTGGCTCGTGCTTGTAAGCTTC
AGCCTATGAAGCGGCCGGGGCCAGGGATCGCGGTTCGAGTCCCGCCTTGGGCCCATAGTGAGACCTAGTCCCAAAAA
AACAGCTGGGCGTGGTGGCGCACGCCTGTAATCCCAGCTACCTGGGAAGCTGAGGCTGGGGATGGCTTGAGTTCGTG
GTACCCCGGGCAACAAGAGCCTGCGGTTCCACAGGCAAAAATAGCGACACCTAGTCTCAAAAAAAAAAAAAAAAAAA
AAAAAAAAAAAAAAAAAAAAAAAAAAAAAAAAAAAAAAAAAAAAAAAAAAAAAAAAAAAAAAAAAAAAAAAAAAAAA
AAAAAAAAAAAAAAAAAAAAAAAAAAAAAAAAAAAAAAAAAAAAAAAaaaagtaagtgcattatttattaactcgat
gaaatcagctgtcttggaaatatgttgaattccacaaccccttttcttaggcaagga

PVHY01011283.1:21875-22611(+). **A**$_{165}$

taagagcagaagtgaacaaaatagagatcaaaaagatcatttcaGGCTGGGCATGGTGGCGCATGCCTGTAAGCAGC
AGCTTAGGAAGCTGTGGCCGGGGCCAGGGATTGCGGTTTGAGTCCCACTGGGGGCACATAATGAGACCTAGTCCCAA
AAAAATAGCTGGGCACGGTGGCGCATGCCTGTAATCTCAGCGACCTGGAAAGCTGAGGCTGGGGATTGCTTGATCTG
TGGTACCCTGGGCAATGTGAGCCTGCGTGCCACAGGCAAAAATTGCAATACATAGTCTCAAAAAAAAAAAAAAAAAA
AAAAAAAAAAAAAAAAAAAAAAAAAAAAAAAAAAAAAAAAAAAAAAAAAAAAAAAAAAAAAAAAAAAAAAAAAAAAA
AAAAAAAAAAAAAAAAAAAAAAAAAAAAAAAAAAAAAAAAAAAAAAAAAAAAAAAAAAAAAAAAAaaaaagatcatt
tcaaatatcactgaaaccaaaagatgcttcattgaaaaaactgacaaaacagatacac

PVHY01001295.1:147414-148147(+). **A145**

agcgagcaggaccaactggatttcctccgtcaagcaggtaagctgGGCGGGCATGGTGGCTCGTGCTTGTAAGCTTC
AGCCTATGAAGCGGCCGGAGCTAGGGATCGCAGTTCGAGTCCTGCCTGGGGCACGTAGCGAGACCTAGTCCCAAAAA
AATAGCTGGGCATGGTGGCGCGCGCCTGTCATCCCAGCTACCTAGGAAGCTGAGGCTGAGGATCGCTTGAGTCTGTG
GTACCACGGGCAATGAGAGCCTGCGGTTCCACGGGCAAAAATAGCGAGACCTAGTCAAAAAAAAAAAAAAAAAAAAA
AAAAAAAAAAAAAAAAAAAAAAAAAAAAAAAAAAAAAAAAAAAAAAAAAAAAAAAAAAAAAAAAAAAAAAAAAAAAA
AAAAAAAAAAAAAAAAAAAAAAAAAAAAAAAAAAAAAAAAAAAAAaagcaggtaagctgggaaggagagagcatatg
acagtcagagtgggaactactgattagcccagagggaggaccttaaagctgacccac

PVHY01000291.1:22510-23241(-). **A161**

tgggctaaaagaagaaaacttagaaggttgaaaagagaagatgaaGCCGGGCATGGCGGCTCGTGCCTGTAAACTTC
AGGCTATGAAGCGGCTGAGGCTAGGGATCATGGTTCGAGTACTGTTCGGGGCAACACAGAGAGACCAAGTCACAAAA
AATAGCTGGGTGTGGTGGCGCACGCCTGTTATCCCAGCTACCTGGGAAGCTGAGGCTGGGGATTGCTTGAGTCTGTG
GTACCCTGGGCAACGTGATACTGCGGTTCCATGAGAAAATAACGGGACCTGTCTCCAAAAAAAAAAAAAAAAAAAAA
AAAAAAAAAAAAAAAAAAAAAAAAAAAAAAAAAAAAAAAAAAAAAAAAAAAAAAAAAAAAAAAAAAAAAAAAAAAAA
AAAAAAAAAAAAAAAAAAAAAAAAAAAAAAAAAAAAAAAAAAAAAAAAAAAAAAAAGGAAAAAGaaagagaagatga
aaaggtcaaaactggattatgagttaataaaaagagaaaattaagcataaacgat

PVHY01004644.1:82873-83600(-). **A178**

tcatataaagtatgtcaggagttttaacagaatgggcaccctggGGCCGGGCACCATGGCTCGTGCCTGTAAGCTTC
AGCCTATGAAGTGGCCAGGGCTAGGGATCGCGGTTTGAGTCCCGCCTGGGGCCCGTAGCGAGACCTAGTTCCAAAAA
AATAGCCGGGCATGGTGGTACGTGCCTGTAATCCCAGCTACCTGGGAAGCTGAGGCTGGGGATTGCTTGAGTCTGTG
TTACCCTGGGCCTGCAGTTCCATGGGCAAAAAATAGGGAGACCTAGTCTCAAAAAAAAAAAAAAAAAAAAAAAAAAA
AAAAAAAAAAAAAAAAAAAAAAAAAAAAAAAAAAAAAAAAAAAAAAAAAAAAAAAAAAAAAAAAAAAAAAAAAAAAA
AAAAAAAAAAAAAAAAAAAAAAAAAAAAAAAAAAAAAAAAAAAAAAAAAAAAAAAAAAAAAAAAAAAAAAAAAAggc
accctgggataatgtgatataataaatgtaaatttggcaacattgatta

PVHY01006420.1:49701-50435(+). **A139**

agtaactgcgatcccctgctagaacctcaaaaaaagctctcaaggGCCGGGCACAATGGCTCATGCCTGTCAGCTTT
AGCCTATGAAGCGGCTGGGGCTAGGGACCATGGTTTGAGTCCCGCCTGGGGCCCATAGCATGATCTAGTTCCAAAAA
AATAGCTGGGCGTGGTGGTACATGCCTGTAATCCCAACCATCTGGGAAGCTGAGGCTGAGGATTGCTTGAGTCTGTG
GTACCCCGGGCAATGAGAGCCTGCGGTTCCATGGGCAAAAATAGCGAGAACTAGTCTTAAAAAAAAAAAAAAAAAAA
AAAAAAAAAAAAAAAAAAAAAAAAAAAAAAAAAAAAAAAAAAAAAAAAAAAAAAAAAAAAAAAAAAAAAAAAAAAAA
AAAAAAAAAAAAAAAAAAAAAAAAAAAAAAAAAAAAaaaaaaagctctcaaggtcctaaataataagtcttataacc
ttgagacttccatgctataagtaaacccaaacttgctgtgtggggaggccacatac

PVHY01020753.1:10561-11303(+) . **A158**

cattctatgtatcttgttctggtgactcaaagctatgagcatagAGCCGGACACAGTGGCGTGTGCCTGTAAGTGCC
AGCCTATGGAATGCGGATGAGACTGGGGACCTTGGTTTGAGTCCCGCCCGGAGCAACATAGCGAGACCTAGTTCCAA
AAAAATATAGCTGGACATGGTGGCGTGCACTATAATCCCACCTATATGGGAAGCTGAGGCTGGGGATTGCTTGAGTC
TGTGGTACCCTGGGCAACATGAGCCTGTGGTTCCATGAGCAAAAATAGTGAGACCTACCTAGTCTCAAAAAAAAAAA
AAAAAAAAAAAAAAAAAAAAAAAAAAAAAAAAAAAAAAAAAAAAAAAAAAAAAAAAAAAAAAAAAAAAAAAAAAAAA
AAAAAAAAAAAAAAAAAAAAAAAAAAAAAAAAAAAAAAAAAAAAAAAAAAAAAAAAAAAAAAAAAAAaaagctatga
gcatccttgcacaggttgtgattattttgtaatagtggctatttcttgataaaattccaattaa

**Figure S5**. Examples of Urop_c copies with extra-long poly(A) tails (from $A_{139}$ to $A_{178}$). Flanking sequences are shown in lowercase. TSDs are underlined. Sequences corresponding to the promoter boxes A and B are highlighted in blue and yellow, respectively. The length of pure poly(A) sequence is specified after the sequence name.
