## Supplementary material for "A Dimeric SINE Discovered in Shrew Mole is Structurally Similar to Primate Alu": Fig. S6

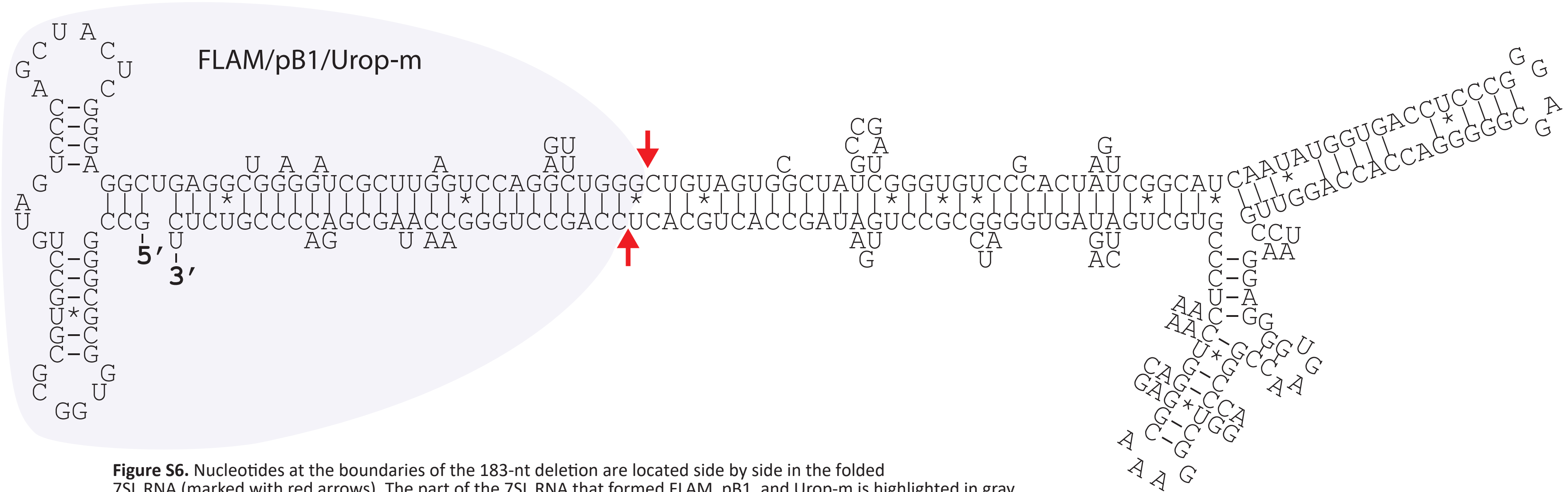


**Figure S6.** Nucleotides at the boundaries of the 183-nt deletion are located side by side in the folded 7SL RNA (marked with red arrows). The part of the 7SL RNA that formed FLAM, pB1, and Urop-m is highlighted in gray. The secondary structure of the human 7SL RNA was established by digestion with nucleases specific to single- and double-stranded RNA regions (Gundelfinger ED, Di Carlo M, Zopf D, Melli M. Structure and evolution of the 7SL RNA component of the signal recognition particle. EMBO J. 1984:3(10)2325-2332. doi: 10.1002/j.1460-2075.1984.tb02134.x).
